# Effects of chronic exposure to fluoxetine, eicosapentaenoic acid, and lipopolysaccharide on behavior and hippocampal transcriptome in the rat model of prolonged chronic unpredictable stress

**DOI:** 10.1101/2021.12.05.471274

**Authors:** Konstantin A. Demin, Tatiana O. Kolesnikova, David S. Galstyan, Nataliya A. Krotova, Nikita P. Ilyin, Ksenia A. Derzhavina, Maria Seredinskaya, Yuriy M. Kositsyn, Dmitry V. Sorokin, Maria O. Nerush, Abubakar-Askhab S. Khaybaev, Sofia A. Pushkareva, Elena V. Petersen, Murilo S. de Abreu, Alexey Masharsky, Allan V. Kalueff

**Affiliations:** Almazov National Medical Research Centre, St. Petersburg, Russia; Institute of Translational Biomedicine, St. Petersburg State University, St. Petersburg, Russia; Neuroscience Program, Sirius University of Science and Technology, Sochi, Russia; Core Facility Centre for Molecular and Cell Technologies, St. Petersburg State University, St. Petersburg, Russia; Moscow Institute of Physics and Technology, Moscow, Russia

**Keywords:** chronic unpredictable stress, behavior, antidepressants, rats, transcriptome

## Abstract

Animal models are widely used to study stress-induced affective disorders and associated with them neuroinflammation and other neuroimmune processes. Here, we examined rat behavioral and hippocampal transcriptomic responses to prolonged chronic unpredictable stress (PCUS), as well as following a 4-week treatment with a classical antidepressant fluoxetine, an anti-inflammatory agent eicosapentaenoic acid (EPA), a pro-inflammatory agent lipopolysaccharide (LPS) and their combinations. Overall, PCUS evoked an anxiety-like behavioral phenotype in rats (corrected by chronic fluoxetine alone or combined with other drugs), EPA was anxiolytic and LPS promoted anxiety in this model. PCUS evoked pronounced transcriptomic responses in rat hippocampi, including >200 differentially expressed genes. While pharmacological manipulations did not affect hippocampal gene expression markedly, *Gpr6*, *Drd2* and *Adora2a* were downregulated in stressed rats treated with fluoxetine+EPA, suggesting G protein-coupled receptor 6, dopamine D2 receptor and adenosine A2A receptor as potential evolutionarily conserved targets in chronic stress. Overall, these findings support the validity of rat PCUS paradigm as a useful tool to study stress-related affective pathologies and calls for further research probing how various conventional and novel drugs modulate behavioral and neurotranscriptomic biomarkers of chronic stress.

## Introduction

While stress potently activates sympatho-adrenomedullary, hypothalamic-pituitary-adrenal^1,2^, metabolic and immune systems^3,4^, chronic stress evokes lasting behavioral and physiological pathological responses^5,6^, including neuroendocrine and neuroimmune deficits^7–11^. In humans, this often triggers anxiety, depression and other affective disorders^12–15^ - widespread, severely debilitating and poorly understood mental illnessess^16–20^. Various animal models, especially rodents and zebrafish (*Danio rerio),* are commonly used to study the effects of stress on brain and behavior^21–23^. Typically utilizing chronic unpredictable stress (CUS) protocols^24–28^, such studies expose rodents to varying stressors for several weeks^26,28–31^, evoking anxiety- and depression-like states^32–34^ that resemble those observed in humans^35^.

However, clinically relevant chronic stress usually lasts much longer than 5 weeks, and conventional antidepressants take several weeks to act, thus making short-term CUS models less optimal for paralleling clinical setting^36,37^. To address this problem, we have recently developed a novel prolonged chronic unpredictable stress (PCUS) model in zebrafish, based on >10-week stress and >3-week antidepressant treatment^38^. Capitalizing on this model, here we attempted to translate it into a rat chronic stress battery, developing a 12-week PCUS rat protocol, which we also modulated using 4-week treatment by a serotonergic antidepressant fluoxetine, a neuroprotective omega-3 polyunsaturated fatty acid (PUFA) eicosapentaenoic acid (EPA), and a pro-inflammatory bacteria-derived lipopolysaccharide (LPS) alone, or combined with fluoxetine. Our rationale for combining the drugs was based on the idea that a combination of antidepressant fluoxetine with anti-inflamatory EPA here may be synergetic and more beneficial, targeting both neurochemical and proinflammatory disturbances, respectively, which are both observed under chronic stress. In contrast, testing LPS effects and combining it with fluoxetine meant to assess the efficacy of antidepressant treatment in an adverse scenario when chronic stress is ‘potentiated’ by neuroinflammation. The scope of the present study was to assess behavioral and hippocampal transcriptomic responses to PCUS in rats, and their modulation by a 4-week treatment with fluoxetine, EPA and LPS alone or administered jointly.

## Materials and methods

### Animals

A total of 140 young male Wistar rats (2-2.5 months) were obtained from the Center for Preclinical and Translational Research of Almazov National Medical Research Centre (St. Petersburg, Russia). Prior to and during testing, rats were kept under standard conditions (20–22°C, 55% humidity, food and water *ad libitum*, 12:12 h light/dark cycle; lights on at 08:00 am). All rats were from the same population and were randomly allocated into experimental groups using a random number generator (https://www.random.org/). All experimental animal manipulations were approved by the Ethics committee of the Institute of Experimental Medicine at Almazov National Medical Research Center (approval number 20-14PZ#V2). All animals tested were included in final analyses, without removing the outliers. All experiments were performed as planned, and all analyses and endpoints assessed were included without omission.

Rodents are widely used as a tool for CNS pathology modeling, including stress-related pathological states, due to the high homology of the core stress mechanisms with humans. The Wistar strain was chosen here as the best-studied and widely used rat strain in biomedical modeling and preclinical studies^39^, and because studies with this strain are highly reproducible in neurogenetics and CNS disorders modeling, given genetic stability of this stain^39^. Only male rats were used in the present study, chosen based on overt sex differences in rat chronic stress assays^40–42^ and in order to mitigate the impact of the estrous cycle on female rat behavior^43^ which may unpredictably affect the results.

### Prolonged Chronic Unpredictable Stress (PCUS) procedure

Experimental rats were exposed for 12 weeks to various stressors daily, similar to ^44,45^, including crowding, smell, novel objects, flashing light, water/food deprivation, shaking, swimming, novelty, day/night inversion, predator exposure, darkness and light for 24 h, intermittent and stroboscopic lighting, cage tilt, noise (drill sound), social isolation, and sleep deprivation (Tables 1 and 2, Fig. 1). The duration of stress exposure was chosen here by extending previous rodents CUS protocols^45^ and adapting the recently developed PCUS procedure in zebrafish^38^.

**Figure 1.**
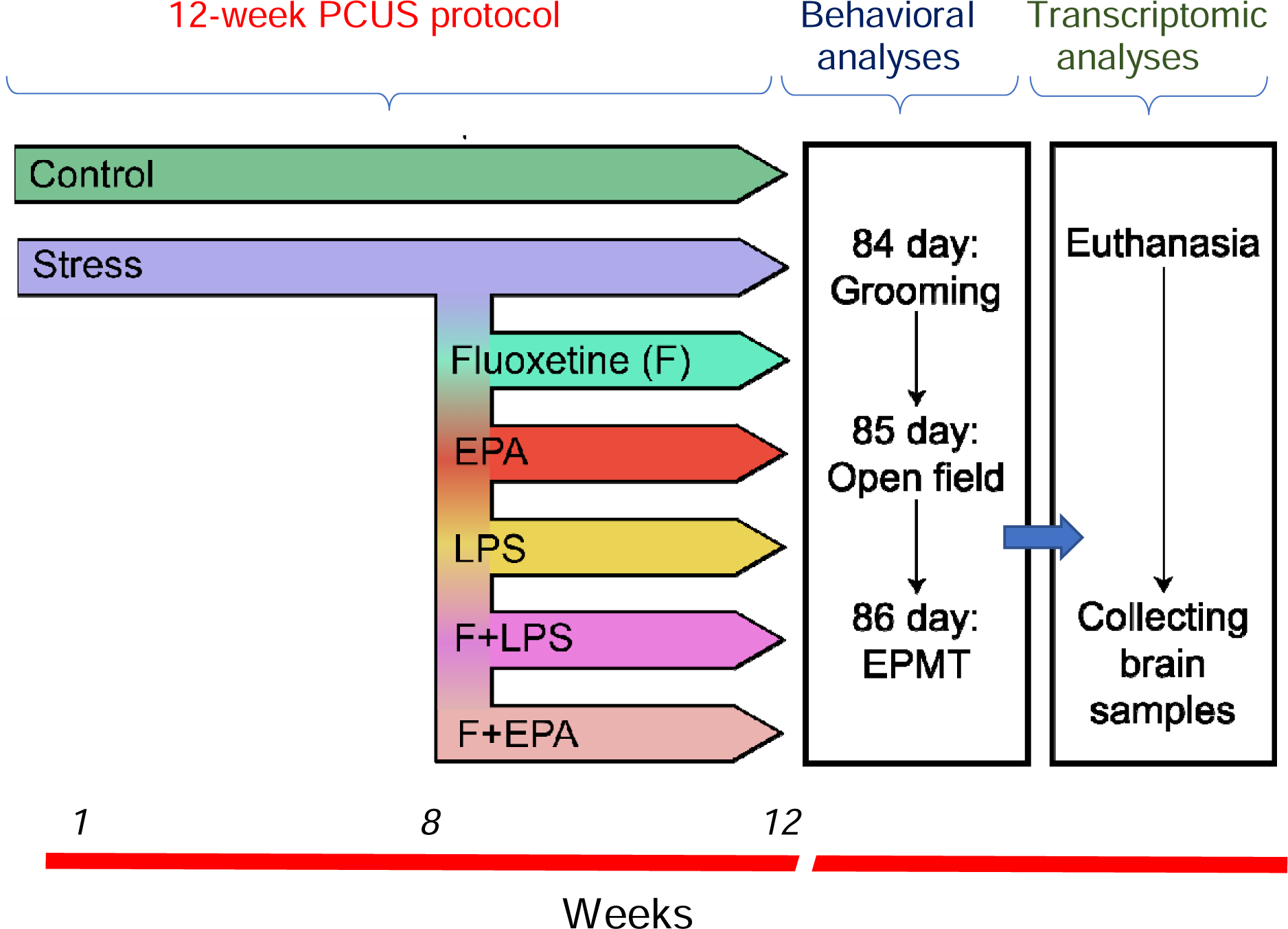
Brief diagram summarizing the study experimental design. PCUS – prolonged chronic unpredictable stress, EPA – eicosapentaenoic acid, LPS – lipopolysaccharide, FEPA – fluoxetine + EPA, FLPS – fluoxetine +LPS, EPMT – elevated plus maze test.

**Table 1.** Summary of the rat PCUS protocol used in the present study (see. Table 2 **for details of specific stressors)**

| Day | Stress procedures |
| --- | --- |
| 1 | Strong smell + noise 30 min + bright light (150 Lx) 24 h |
| 2 | Flashing light 4 h + crowding 6 h + water deprivation 12 h |
| 3 | Social isolation 6 h + cage tilt 24 h |
| 4 | Intermittent lighting + swimming 20 min + food deprivation |
| 5 | Shaking 20 min + noise 4 h + day/night inversion |
| 6 | Novelty stress + predator exposure 24 h + darkness 24 h |
| 7 | Stroboscopic lighting (150 Lx) 6 h+ novel objects + strong smell |
| 8 | Intermittent lighting + cage tilt 24 h + day/night inversion |
| 9 | Crowding 8 h + shaking 30 min |
| 10 | Food and water deprivation (with empty bottles) 12 h + light 24 h |
| 11 | Flashing light 8 h + noise 2 h + sleep deprivation |
| 12 | Swimming 30 min + shaking 10 min + predator exposure 24 h |
| 13 | Novel objects 24 h + day/night inversion |
| 14 | Strong smell + noise 4 h + intermittent lighting |
| 15 | Social isolation 10 h + cage tilt 24 h |
| 16 | Novelty stress + predator 24 h + light 24 h |
| 17 | Noise 6 h + flashing light 4 h + food deprivation 24 h |
| 18 | Swimming 30 min + water deprivation 12 h + day/night inversion |
| 19 | Crowding 8 h + strong smell + light 24 h |
| 20 | Flashing light 10 h + cage tilt 24 h + predator 24 h |
| 21 | Stroboscopic lighting 8 h + novel objects 24 h |
| 22 | Strong smell + intermittent lighting + sleep deprivation 24 h |
| 23 | Noise 8 h + food/water deprivation + darkness 24 h |
| 24 | Flashing light 8 h + novel objects and cage tilt 24 h |
| 25 | Swimming 30 min + stroboscopic lighting 6 h + predator exposure 24 h |
| 26 | Crowding 6 h + food deprivation 10 h + darkness 24 h |
| 27 | Shaking 40 min+ flashing light 8 h |
| 28 | Social isolation 8 h with noise 6 h + day/night inversion |
| 29 | Novelty stress + stroboscopic lighting with novel objects 8 h + predator exposure 24 h |
| 30 | Crowding 8 h + water deprivation (with empty bottle) 12 h + cage tilt 24 h |
| 31 | Shaking 35 min + darkness 24 h with flashing light (4 h) |
| 32 | Noise 6 h + intermittent lighting + sleep deprivation |
| 33 | Social isolation 8 h + light 24 h + food deprivation 24 h |
| 34 | Swimming 25 min + intermittent lighting + cage tilt 24 h |
| 35 | Flashing light 10 h + novel objects 24 h + noise 2 h |
| 36 | Shaking 30 min + predator 24 h + light 24 h |
| 37 | Noise 4 h + food deprivation + day/night inversion |
| 38 | Crowding 6 h + intermittent lighting |
| 39 | Shaking 40 min + novel objects + stroboscopic lighting 6 h |
| 40 | Swimming 20 min + predator 24 h + water deprivation 24 h |
| 41 | Social isolation 10 h + darkness 24 h + food deprivation 24 h |
| 42 | Noise 6 h + flashing light 8 h + cage tilt 24 h |
| 43 | Novel objects 24 h + strong smell + light 24 h |
| 44 | Swimming 5 min + stroboscopic lighting 8 h + predator 24 h |
| 45 | Crowding 10 h + noise 12 h + food deprivation 24 h |
| 46 | Novelty stress + shaking 20 min + darkness 24 h |
| 47 | Noise 4 h + intermittent lighting + predator 24 h |
| 48 | Water deprivation 10 h + cage tilt 24 h + day/night inversion |
| 49 | Social isolation 8 h + noise 2 h + stroboscopic lighting 2 h |
| 50 | Shaking 30 min + predator 24 h + darkness 24 h |
| 51 | Strong smell + intermittent lighting + novelty stress |
| 52 | Noise 6 h + crowding 12 h + sleep deprivation |
| 53 | Swimming 30 min + flashing light 4 h + predator 24 h |
| 54 | Social isolation 6 h + food deprivation 24 h + light 24 h |
| 55 | Strong smell + novel objects 24 h + cage tilt 24 h |
| 56 | Stroboscopic lighting 10 h + noise 5 h + day/night inversion |
| 57 | Social isolation 8 h + food deprivation 24 h + darkness 24 h |
| 58 | Shaking 20 min + predator 24 h + flashing light 6 h |
| 59 | Novel objects + intermittent lighting+ water deprivation with empty bottle 24 h |
| 60 | Strong smell + noise 6 h + novel objects 24 h |
| 61 | Crowding 10 h + cage tilt 24 h + light 24 h |
| 62 | Swimming 20 min + intermittent lighting + predator 24 h |
| 63 | Water deprivation with empty bottle 12 h + strong smell + day/night inversion |
| 64 | Social isolation 8 h + noise 8 h + food deprivation 24 h |
| 65 | Flashing light 6 h + light 24 h + predator exposure 24 h |
| 66 | Shaking 30 min + crowding 10 h |
| 67 | Noise 2 h + stroboscopic lighting 6 h + cage tilt 24 h |
| 68 | Predator 24 h + darkness 24 h with flashing light (4 h) |
| 69 | Shaking 20 min + noise 8 h + novel objects 24 h |
| 70 | Social isolation 8 h + intermittent lighting + food and water deprivation 24 h |
| 71 | Swimming 20 min + predator 24 h + darkness 24 h |
| 72 | Strong smell + novel objects 24 h + flashing light 10 h |
| 73 | Social isolation 8 h + light 24 h |
| 74 | Noise 8 h + intermittent lighting + food deprivation 24 h |
| 75 | Crowding 10 h + shaking 30 min + predator exposure 24 h |
| 76 | Water deprivation (with empty bottle) 12 h + cage tilt 24 h + day/night inversion |
| 77 | Noise 6 h + stroboscopic lighting 8 h + sleep deprivation 24 h |
| 78 | Predator 24 h + intermittent lighting 8 h |
| 79 | Social isolation 8 h + strong smell 24 h + darkness 24 h |
| 80 | Shaking 40 min + crowding 8 h |
| 81 | Swimming 30 min + stroboscopic lighting with novel objects + noise 4 h |
| 82 | Novelty stress + cage tilt 24 h + day/night inversion |
| 83 | Social isolation 6 h + noise 6 h + sleep deprivation 24 h |
| 84 | Flashing light in darkness 6 h + predator exposure 24 h |
| 85 | Behavioral testing Day 1: grooming test (GT) |
| 86 | Behavioral testing Day 2: the open field test (OF) |
| 87 | Behavioral testing day 3: the elevated plus-maze test (EPMT) |
| 88 | Sacrificing the animals, collecting and storing brain samples |

**Table 2.** Summary of PCUS manipulations used in the present study (see. Table 1 **for details)**

| Stressors | Description | References |
| --- | --- | --- |
| Crowding | Twice increased cage density (to 8 rats per cage) | 165 |
| Strong smell | Sponge soaked with essential oil of lemon and bergamot placed 5-10 cm from the home cage for 24 h. These essential oils were chosen based on their persistent and pungent odor, known to be aversive for rats | 166 |
| Novel objects | Two plastic toys (6-13 cm) placed into the homecage for 24 h | 167 |
| Flashing light | 40-m flashing Christmas lights (5 Hz, 150 Lx) placed on the top of homecages | 168 |
| Water deprivation | Removing water bottle or placing empty bottle into the cage | 169,170 |
| Food deprivation | Removing food from the feeder | 169,170 |
| Shaking | Rolling a cart (100 x 100 x 150 cm) with rat cages on an uneven surface at a constant speed of 6 km/h | 169 |
| Swimming | Rats individually placed in 20-L plastic box filled with water (22-24°) with legs and tail not touching the bottom | 171 |
| Novelty stress | Rats individually placed into novel/unfamiliar plastic boxes (39 x 28 x 28 cm, 56 x 39 x 28 cm or 56 x 39 x 42 cm) for 5 min | 66 |
| Day/night inversion | Darkness in the morning and turning light on during the night | 169,170 |
| Predator exposure | The predator (cat) urine applied to a sponge and placed near the home cage (5-10 cm) | 166 |
| Darkness or light | Light or darkness in the animal homecages for 24 | 170 |
| Intermittent lighting | Turning the lights on and off in the animal room every 2 h for 12 h | 170 |
| Cage tilt | Cages individually tilted at a 45-degree angle | 170,172 |
| Noise (drill sound) | A 50-db drill online sound (www.youtube.com/watch?v=7Xomg2zge-8) | 173 |
| Social isolation | Isolation of rats in a separate plastic box (30 x 20 x 20 cm) with bedding | 174 |
| Sleep deprivation | A cylindrical wooden plinth (6 cm in diameter and 5 cm in height) placed on the floor of the cage opposite the food and water compartment. The cage is filled with water (22°) to a depth of 3 cm, allowing animals to stand, but not sit comfortably or sleep | 175 |
| Stroboscopic lights | Flashing light (160 Lx) exposure produced by a stroboscope at 60 flashes/min 2 m from homecages | 170 |

Control rats were housed similarly to the experimental cohort but remained experimentally naïve for the entire duration of the study. On Day 57, the stressed rodent cohort was divided into several groups (chronic stress alone and chronic stress with chronic fluoxetine, EPA, LPS, as well as fluoxetine+EPA and fluoxetine+LPS) for the final 4 weeks of the study. Since LPS was injected intraperitoneally once a week, control animals were also similarly injected intraperitoneally with sodium chloride solution. Fluoxetine (Biocom Ltd., Stavropol, Russia) is a commonly used antidepressant^46–48^ widely tested in various animal models, including rodents^49–52^. Rats were given 0.5 mL distilled water containing 5 mg/kg fluoxetine or drug-free water using oral syringes. The duration of treatment, its dose and route of administration were selected based on the previous studies in stress-related models^38,53^. LPS from *Escherichia coli O55:B5* (Sigma Aldrich, St. Louis, MO, USA), was chosen here for its ability to induce inflammation^54^ following 0.5-ml weekly injections containing 0.1 mg/kg LPS. The dose was chosen based on rodent LPS studies^55–59^, and adjusted for chronic exposure. EPA was used here for its anti-inflammatory properties^60^, obtained from Tokyo Chemical Industry Co. Ltd. (Tokyo, Japan) and administered by oral syringes in 0.5 mL sunflower oil containing 275 mg/kg of EPA, based on earlier rodent studies^61–64^. Animals from all groups were tested in parallel. On each testing day, all animals were assessed in behavioral tests in the same way, in similar conditions and by the same highly trained experimenters and were euthanized using the same approved procedures.

### Behavioral testing

Following a 12-week PCUS protocol, rat behavioral phenotypes were assessed in a battery of assays consisting of the open field (OFT), elevated plus-maze (EPMT) and grooming tests (GT). Behavioral assays were performed in order of increasing stress intensity, aiming to reduce the impact of the preceding testing experience, according to standards of rodent neurophenotyping. Prior to testing, the rats were kept for 2 h in a testing room for acclimation and were returned to the holding room after testing. Behavioral testing was performed between 10.00 and 17.00 h and was recorded with a SJ4000 action camera (SJCAM, Ltd., Shenzhen, China) at 60 frames/s. Experimenters were blinded to the treatments during behavioral testing and statistical and video analyses, using individual ID codes for rats and rat groups. Manual analysis (where applicable) of behavioral data was performed by two highly-trained observers (blinded to the groups) with inter- and intra-rater reliability of□>□0.85, as assessed by Spearman correlation as part of the laboratory’s standard operating procedure (SOP).

The OFT apparatus was a gray-colored plastic square box 97 length□×□97 width × 40 height, cm (OpenScience Ltd., Krasnogorsk, Russia), mounted on a mobile cart at 55-cm height. The illumination of the arena was 90 Lx. The animals were placed in the center of the apparatus facing the opposite direction to the researcher. The arena was cleaned by sponge with 70% ethanol after each animal and dried with a rag to remove olfactory cues. Each rat was recorded separately, immediately after being taken from the home cage, by a ceiling-mounted SJ4000 action camera, for 5 min, assessing horizontal (total distance traveled, cm) and vertical exploratory activity (total number and duration (s) of supported (paws on the wall) and unsupported (paws in the air) vertical rears), as well as duration and the latency of freezing (s)^65,66^, using Noldus EthoVision XT11.5 (Noldus IT, Wageningen, Netherlands) for automated scoring and RealTimer (OpenScience) software for manual scoring. Freezing was defined in this and other tests here as a period of immobility > 2 s.

The EPMT apparatus consisted of 4 cross-connected gray-colored plastic arms (50 length□×□14 width, cm, OpenScience) placed on 55-cm tall cart, as in ^67^. The two ‘open’ arms had 1-cm edges, and the two ‘closed’ arms had 30 cm-tall walls. The apparatus was illuminated using 65-wt light bulbs, directed to open arms (400 Lx), whereas the testing room and closed arms were dimly lit (30 Lx). During the testing, animals were placed in the central area of the apparatus facing from the researcher for 5 min and their behavior were recorded and analyzed using the RealTimer software, scoring vertical motor activity (total number and duration (s) of supported and unsupported rearing behavior), the number and duration (s) of freezing bouts, as well as the latency (s) to enter and total time spent (s) in open and closed arms^67^. The animal was considered entering the respective arm when hind legs crossed the virtual line between the central platform and the arm. GT was used to characterize in-depth rat self-grooming behavior and its complex behavioral patterns, according to ^68–70^. For this, rats were individually placed in the transparent glass cylindrical jar (20 cm in diameter, 45 cm high)^69^ and their grooming behavior was recorded by a SJ4000 action camera for 10 min, assessing the duration of total, rostral (paw, face and head) and caudal (body, tail and genital) grooming bouts (s)^69,71–73^, analyzed manually offline after the testing by two highly trained observers. We have further visually analyzed grooming microstructure using ethograms and compared the duration (s) for each type of grooming behavior individually, as well as the percent of incorrect grooming transitions for animals with total grooming time >10 s^69,71–73^. Incorrect grooming transitions were defined as any transition between grooming stages that violated normal cephalo-caudal progression (paws->face->head->body->tails/genitals), according to ^69,71–73^.

### Sampling and RNA-sequencing

Brain hippocampal samples for transcriptomic analyses were collected without pooling (i.e., 1 brain per sample) one day after the last behavioral test, between 11:00 and 16.00 h. The 1-day interval was used here to minimize concomitant immediate genomic effects of behavioral testing and/or handling^74^. Hippocampal samples were selected here for its well-studied involvement in depression pathogenesis ^75–77^ and pronounced reduction in both neurogenesis and neuroplasticity that is observed in this area following inflammatory ^78^ and stressful conditions ^79,80^ that is also potentially contributing to affective phenotype. Rats (n=3) for RNA-sequencing analysis were chosen from the groups using a random number generator (https://www.random.org/). Rats were quickly euthanized in a small animal inhalation anesthesia chamber (SomnoSuite, Kent Scientific, CT, USA) using 5 % isoflurane, and their brains were dissected on ice and stored in liquid nitrogen for analyses. RNA isolation was performed using TRI-reagent (MRC, Catalog # 118), according to manufacturer instructions. RNA quality was verified with Quantus, electrophoresis, and QIAxel.

PolyA RNA was purified with Dynabeads mRNA Purification Kit (Ambion, Naugatuck, USA)^74^. Illumina library was made from polyA RNA with NEBNext Ultra II Directional RNA Library Prep Kit for Illumina (NEB) according to the manufacturer’s manual^74^. Sequencing was performed on Illumina HiSeq4000 with 151 bp read length, with at least 20 million reads generated for each sample. Resulting FASTA files and gene counts are available on NCBI’s Gene Expression Omnibus (GEO) ^81^ with accession number GSE205325 (www.ncbi.nlm.nih.gov/geo/query/acc.cgi?acc=GSE205325), according to the necessary guidelines and standards in the field ^82^.

### Statistical analyses and data handling

The study used Generalized Linear Models (GZLM) to analyze behavioral data, similar to ^74^. Briefly, GZLM is a widely used method of statistical analyses^83–85^ of variables with non-normal distributions, thus making it suitable for both nonparametric and parametric data^83,84,86–88^. We performed the Wald chi-square (χ²) analysis of variance (ANOVA, Type II) for GZLM fits, followed by Tukey’s post-hoc testing for significant GZLM/Wald pair-wise comparison data. To count for potential effects of testing day we used test-day, group and their interaction effects to construct GZLM model. However, we further analyzed and discussed only the group effects as the only one relevant to the study aims. To choose optimal GZLM distribution and link functions (goodness of fit) for each endpoint, we compared, where applicable, the Akaike information criterion (AIC) levels^89,90^ of Gaussian distribution (identity link), Poisson distribution (with log or squared root links), Gamma distribution (inverse and log links) and Inverse Gaussian distribution (with inverse and log links), choosing the least AIC score (indicating the model most likely to be correct)^91^, similar to^74^. GZLM analyses were performed using the R software^92^.

Unless specified otherwise, all data were expressed as mean ± standard error of mean (S.E.M.), and P set as < 0.05 in all behavioral analyses. Analyses of all data were performed offline without blinding the analysts to the treatments, since all animals and samples were included in analyses, data were analyzed in a fully unbiased automated method, and the analysts had no ability to influence the results of the experiments, as in^93^. The study experimental design and its description here, as well as data analysis and representation, adhered to the ARRIVE guidelines for reporting animal research and the PREPARE guidelines for planning animal research and testing.

### Differential Gene Expression (DE) and Gene Set Enrichment Analysis (GSEA)

To analyze differential expression (DE) of the genes, reads were mapped to the rat Rnor_6.0 reference genome using STAR spliced aligner^94^ and further processed using featureCounts^95^ to obtain raw gene counts (usegalaxy.org). A total of 32883 genes were used for analyses using the R software^92^ with the Bioconductor^96^ and DESeq2^97^ packages. This method was chosen as an efficient tool to analyze experiments with 12 or fewer replicates per condition, that is stable even within 0.5 fold-change thresholds, and generally consistent with other tools^98^. First, all rows without counts or only with a single count across all samples were removed from analyses, yielding 23684 genes. The Principal Components (PCs) Analysis (PCA) of the regularized log (rlog)-transformed^97^ data counts were used as a preprocessing tool to tackle any outlier samples using pcaExplorer R package^99^. For the PCA analyses, 500 most varied genes were used, and the outliers were determined graphically, using the PC1-PC2 plot, identifying LF3 (LPS-Fluoxetine sample number 3) and K1 (Control sample number 1) as outliers that were excluded from further analysis (Supplementary Fig. S1). PCA of the remaining samples revealed more closely ordered samples with no obvious outliers (Supplementary Fig. S2), but elliptic grouping of the samples with 0.95 CI did not reveal any clear clusters. PC1 and PC2 together determined more than half (58.7%) of the sample variance, whereas PC3-8 each determined less than 15% (Fig. 5). Finally, we identified top 10 down- and up-regulated genes loadings for PC1 and PC2 (genes with largest impact on PC).

DE gene analyses on the Negative Binomial (Gamma-Poisson) distribution were next performed by estimating size factors, dispersion, and negative binomial generalized linear models and Wald statistics using the DESeq function^74,100^. The p-values were adjusted using the Benjamini-Hochberg correction. We further adjusted p-value and false discovery rate (FDR) for multiple comparison using the Bonferroni correction, thus finally setting FDR 0.01(6) for pair-wise comparisons vs. control group and 0.02 for pair-wise comparisons vs. chronically stressed PCUS group. We next compared resultant DE genes between groups identifying uniquely represented as well as co-represented genes. Venn diagrams were constructed using the VennDiagram R package^101^.

The Gene Set Enrichment Analysis (GSEA) is a popular method to assess gene expression data arranged in molecular sets from curated databases, allowing for a better detection of molecularly relevant changes^102–105^. However, original GSEA approaches have some intrinsic limitations, including the inability to handle datasets of different sizes and complex experimental designs^106^. A novel type of GSEA, the Generally Applicable Gene Set Enrichment (GAGE) for the pathways analysis addressed these limitations^106^, enabling to choose independent pathways databases to be analyzed depending on research goals and consistently outperforming classical GSEA methods^106^. The KEGG enrichment analyses were performed on normalized and log2-transformed counts using the GAGE package^106^ and two-sample Student’s t-test for unpaired group comparison of differential expression of gene sets. The FDR cut-off was set at 0.01(6) for pair-wise comparisons vs. control group and 0.02 for pair-wise comparisons vs. chronically stressed group, similarly to the DE analysis. The resultant sets were compared between groups, similarly to DE analysis, and visualized by Venn diagrams, as in ^107^.

## Results

Behavioral analyses in the OFT support the validity of PCUS as an anxiogenic/stressogenic model, revealing overt anxiety-like behaviors in rats (Table 3, Fig. 2 and Supplementary Tables S1-S2), as stressed rats reduced the frequency and duration of vertical exploratory rearing behavior, and increased freezing frequency and duration (p<0.05 vs. control, Tukey test). In contrast, fluoxetine treatment during the last 4 weeks of PCUS normalized most rat OFT behaviors (p<0.05, Tukey test), thus showing a predictable anxiolytic/anti-stress-like response. While EPA, similarly to fluoxetine, also normalized most behavioral endpoints (e.g., bringing rearing and freezing frequency to control levels, p>0.05 vs. control, Tukey test), it did not normalize freezing duration vs. control (p<0.05, Tukey test), suggesting somewhat lesser efficacy in alleviating PCUS than fluoxetine. Fluoxetine combinations with EPA or LPS were similarly effective in the OFT, increasing the duration and the frequency of vertical rearing vs. the PCUS group (p<0.05), likely due to the action of fluoxetine that was prevailing.

**Figure 2.**
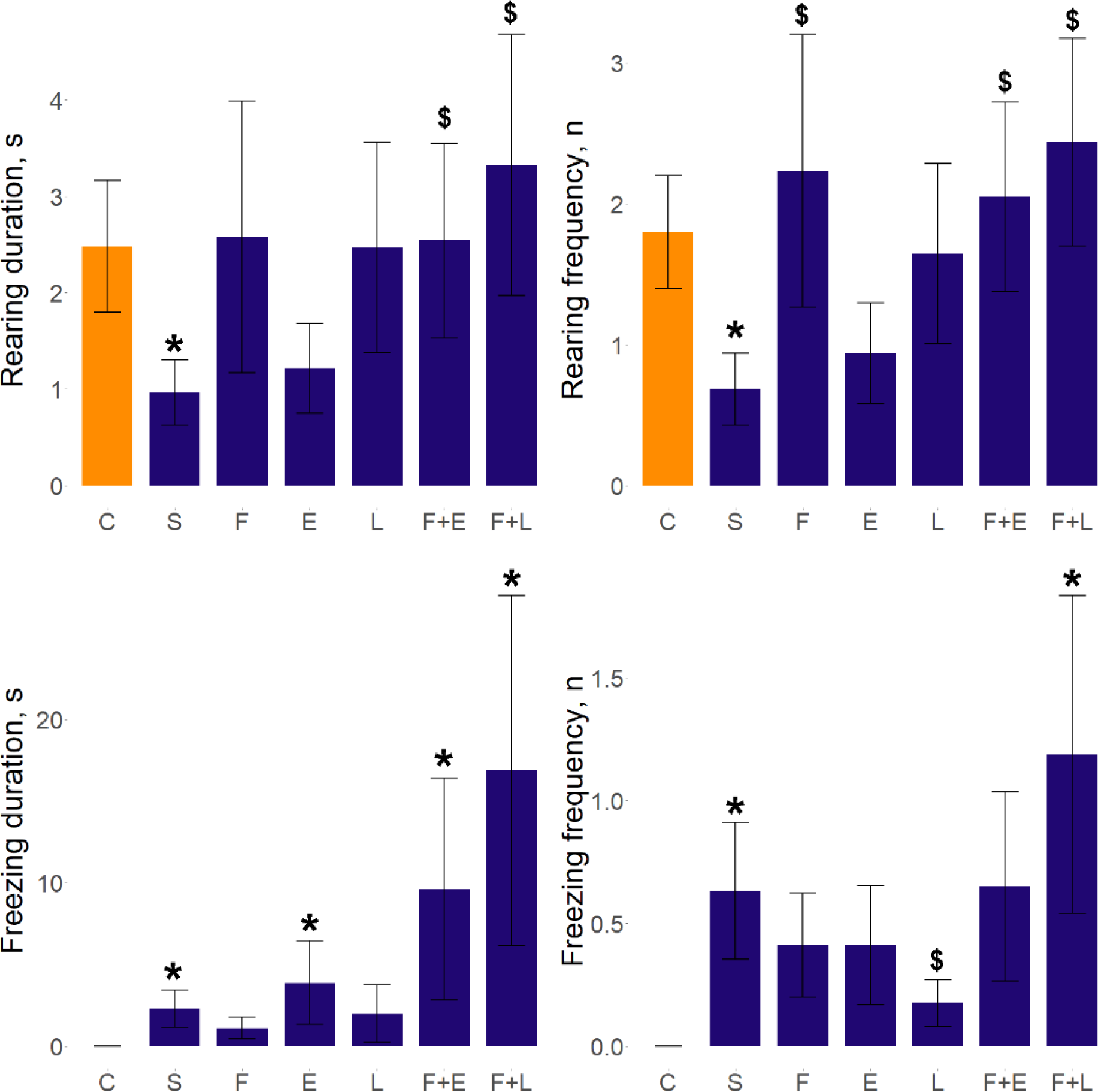
Behavioral effects induced by prolonged chronic unpredictable stress (PCUS) exposure and fluoxetine, eicosapentaenoic acid (EPA) or lipopolysaccharide (LPS) treatment in rat assessed in the open field test. Data are presented as mean ± SEM (n=16-20). *p<0.05 vs. control, $p<0.05 vs. PCUS, post-hoc Tukey test for pair-wise comparison of significant Wald Chi-square (χ²) analysis of variance (ANOVA Type II) for GZLM fits data. Graphs were constructed using the ggplot2 R package^176^, also see Table 3 and Supplementary Tables S1-S2 for statistical details. C - control, S – PCUS, F – fluoxetine, E – EPA, L - LPS groups.

**Table 3.**
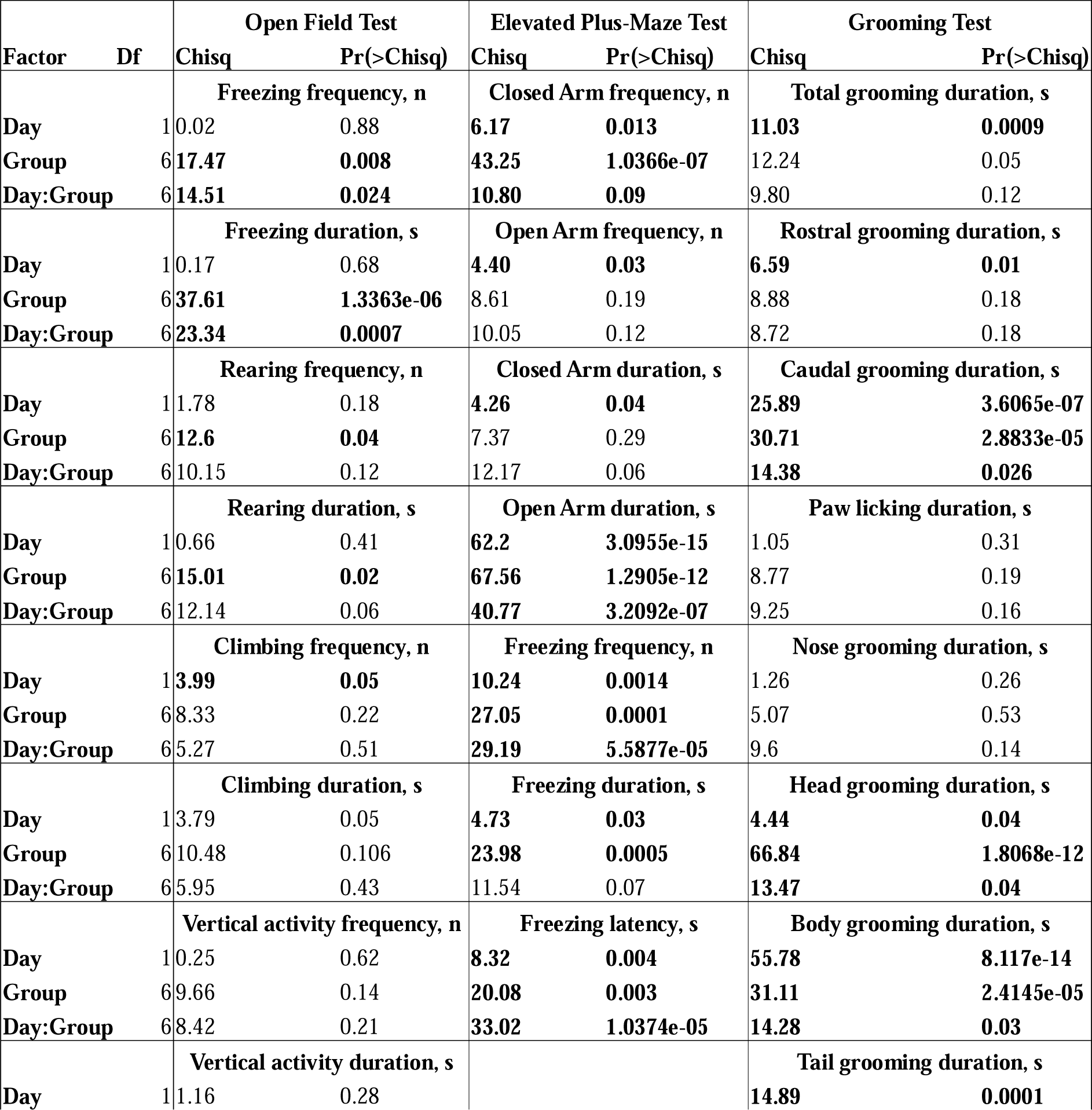
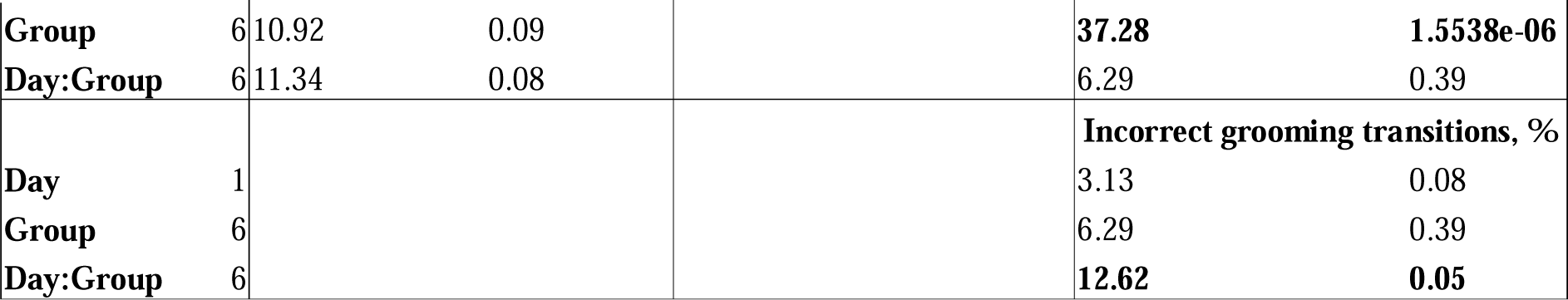
Summary of the Wald Chi-square test results (ANOVA Type II) for generalized linear model (GZLM; Supplementary Table S1) using group, testing day and their interaction effects as ‘predictors’, to compare behavior of experimental rat groups (also see Supplementary Table S2 for Tukey test pair-wise Group comparison data). Note that while we used day, group and interaction to perform GZLM and ANOVA test, here we did not discuss day or interaction effects, and used them only to minimize any potential testing day effects on group factor in the models. Bolded text corresponds to significant ANOVA Type II treatment effects for corresponding endpoints (p<0.05, ANOVA Type II).

In the EPMT, PCUS-exposed rats predictably spent less time in ‘aversive’ open arms than controls (p<0.05, Tukey test vs. control, Table 3, Fig. 3 and Supplementary Tables S1-S2), whereas both fluoxetine or EPA normalized this behavioral deficit in the PCUS group (p>0.05, Tukey test). Moreover, fluoxetine+LPS treatment increased closed entries (p<0.05, Tukey test), potentially suggesting anxiogenic-like effects of LPS. No other overt effects were observed in the number of open arm entries, time spent in closed arms and freezing endpoints in the EPMT in all experimental groups (p>0.05, Tukey test, Table 3, Fig. 3 and Supplementary Tables S1-S2).

**Figure 3.**
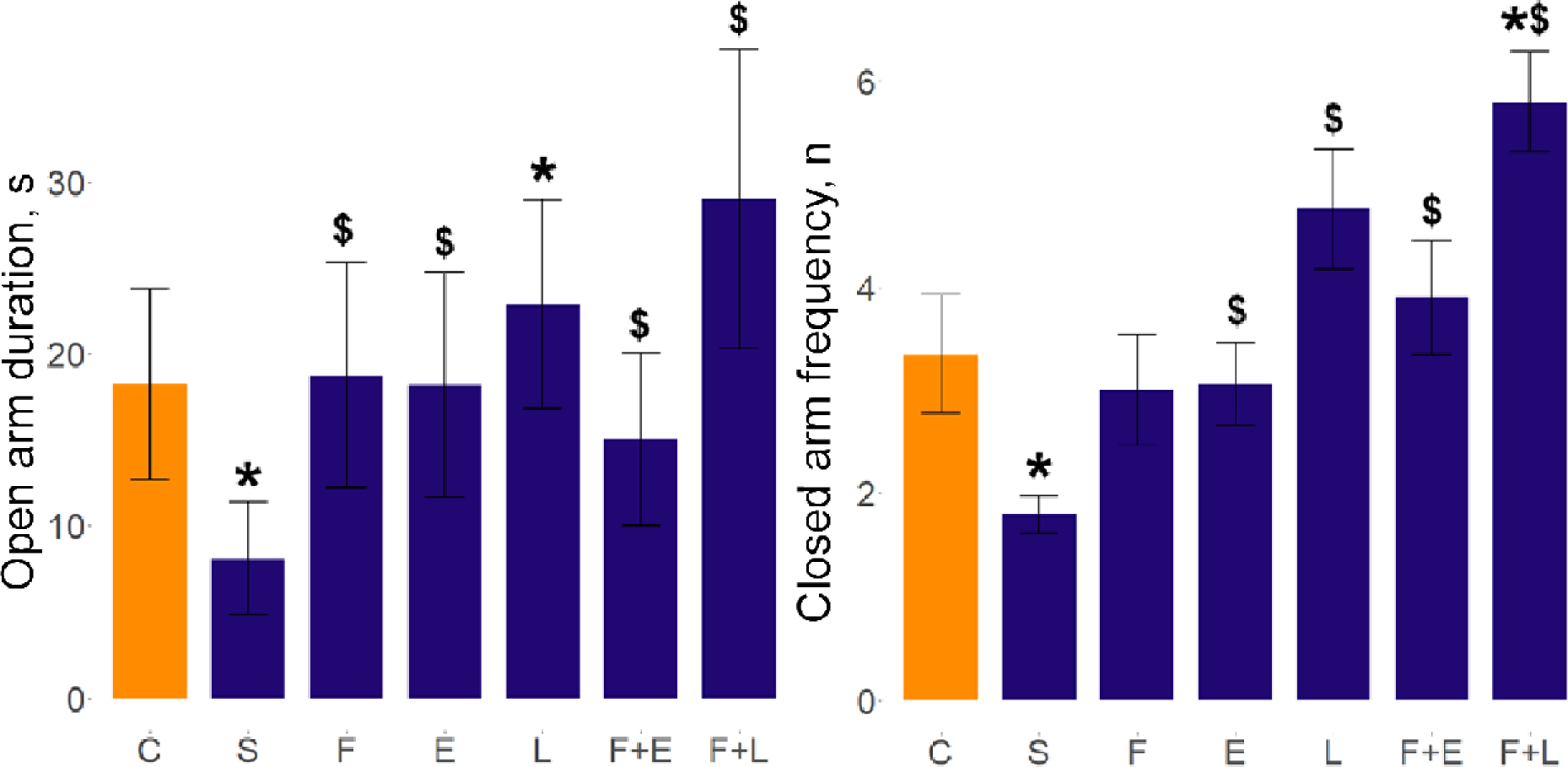
Behavioral effects in the rat elevated plus-maze test induced by prolonged chronic unpredictable stress (PCUS) and fluoxetine, eicosapentaenoic acid (EPA) or lipopolysaccharide (LPS) treatment. Data are presented as mean ± SEM (n=15-20). *p<0.05 vs. control, $p<0.05 vs. PCUS post-hoc Tukey test for pair-wise comparison of significant Wald chi-square (χ²) analysis of variance (ANOVA Type II) for GZLM fits data. Graphs were constructed using the ggplot2 R package^176^ (also see Table 3 and Supplementary Tables S1-S2 for statistical details). C - control, S – PCUS, F – fluoxetine, E – EPA, L - LPS groups.

In the GT, PCUS induced significant effects on rat self-grooming (Fig. 4 and Supplementary Tables S1-S2), increasing caudal grooming in most experimental groups vs. control (p<0.05, Tukey test) and, hence, supporting the value of this stress model. Stress-induced grooming activity was reduced by EPA vs. PCUS group, implying some anti-stress effects of these treatments in the model (p<0.05, Tukey test, Table 3, Fig. 4, Supplementary Tables S1-S2). Furthermore, GT microstructure analyses revealed increased body-, tail- and head self-grooming in all groups (except fluoxetine+LPS) vs. controls (p<0.05, Tukey test, Table 3, Fig. 4, Supplementary Tables S1-S2).

**Figure 4.**
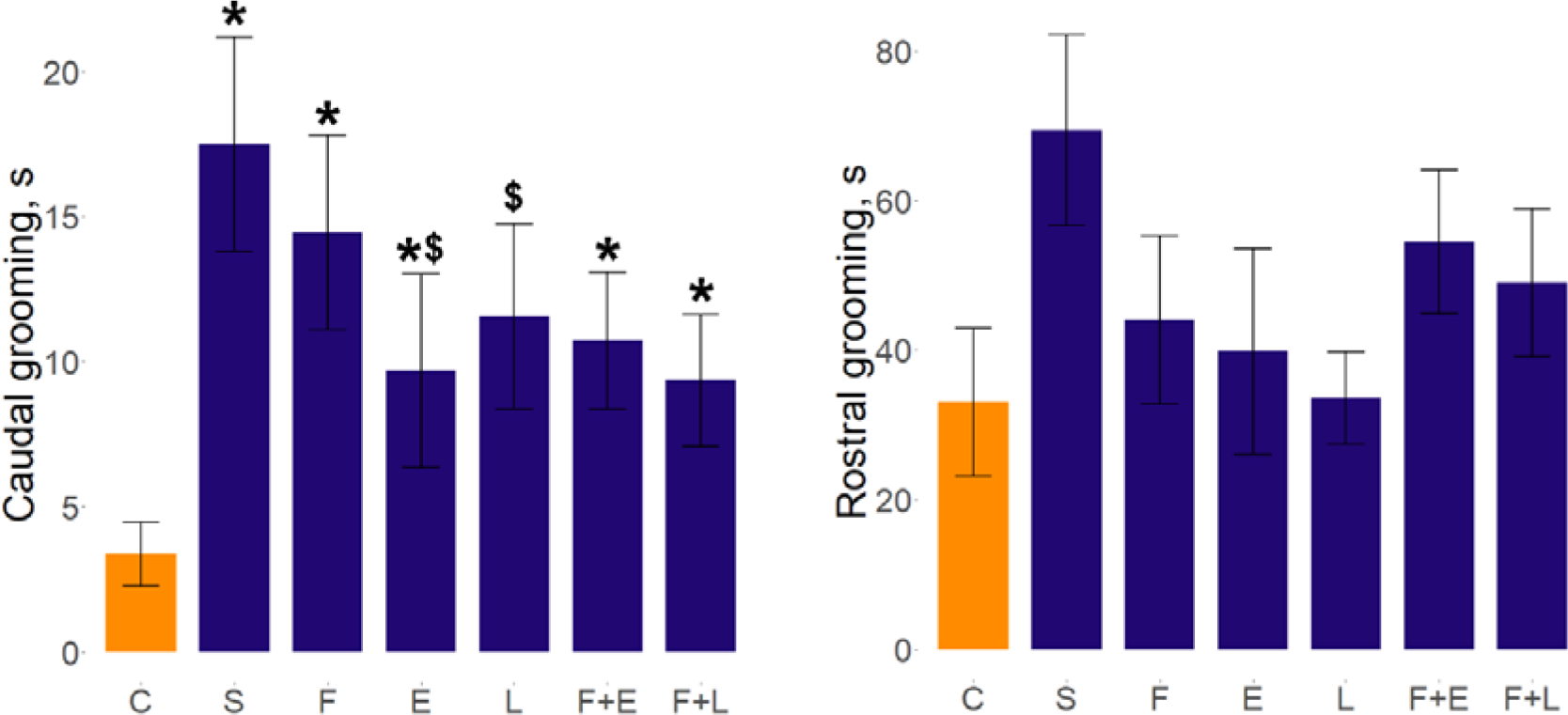
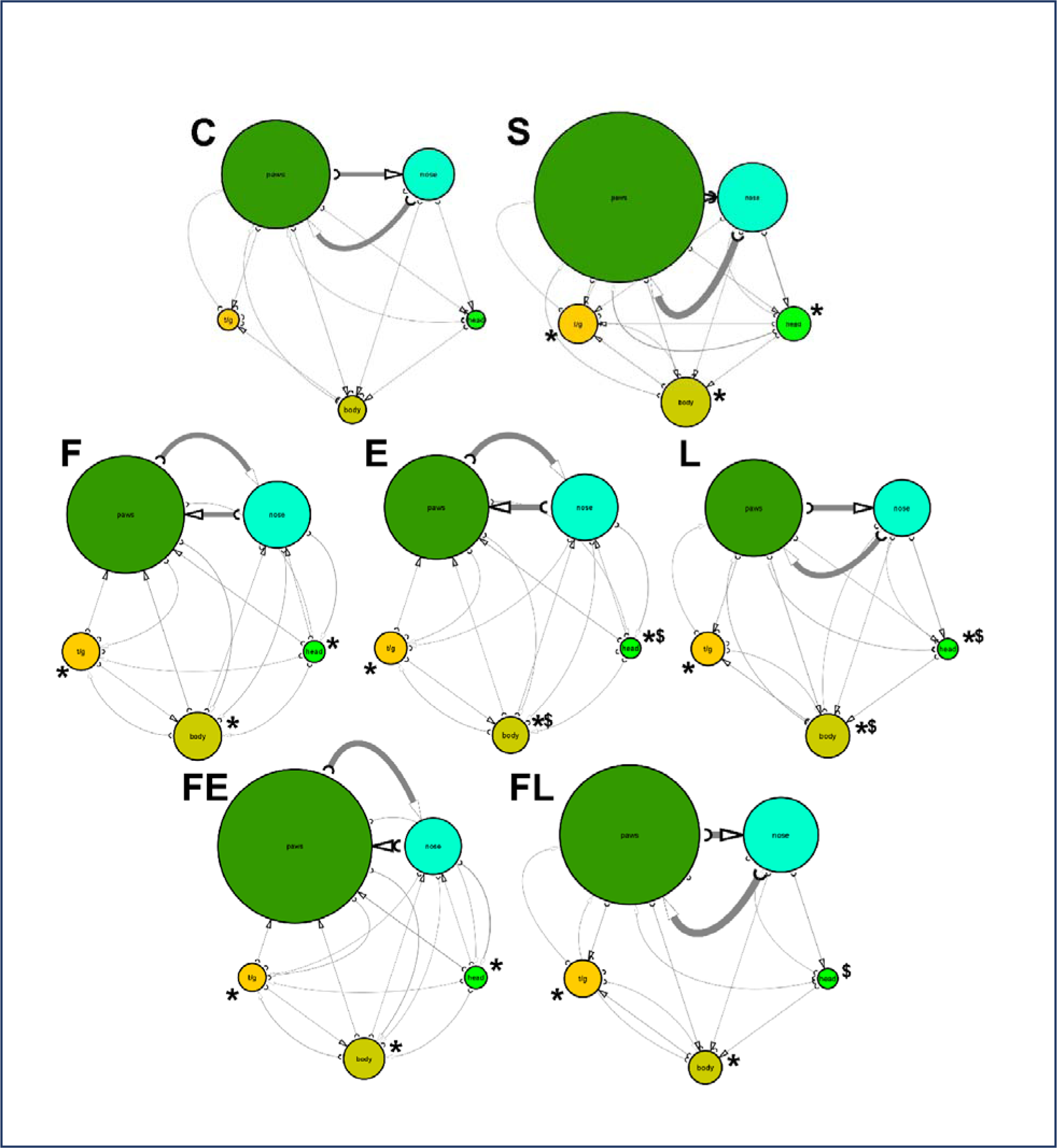
Summary of behavioral effects in the rat grooming test induced by prolonged chronic unpredictable stress (PCUS) and fluoxetine, eicosapentaenoic acid (EPA) or lipopolysaccharide (LPS) treatment (top panel). Data are presented as mean ± SEM (n=15-20). *p<0.05 vs. control, $p<0.05 vs. PCUS post-hoc Tukey test for pair-wise comparison of significant Wald chi-square (χ²) analysis of variance (ANOVA Type II) for GZLM fits data. Graphs were constructed using the ggplot2 R package^176^, also see Table 3 and Supplementary Tables S1-S2 for statistical details). Inset: ethograms-based grooming analyses, showing grooming microstructure of experimental groups in the grooming test. Note overall increase in body, tail and head grooming in PCUS (S) vs. the control (C) group. F – fluoxetine, E – EPA, L - LPS groups, t/g – tail and genital grooming. The relative circle size for each grooming type reflects the duration the observed behavior.

Examining hippocampal transcriptomic data, PCA did not identify overt clusters among brain samples tested (Supplementary Fig. S2), indicating that no group gene expression profiles had pronouncedly distinct expression profiles. However, PCA revealed top 10 up- and down-regulated PC1 and PC2 genes with the most load on these PCs (Fig. 5). For example, *Plcxd2* and *Foxp2* were, respectively, most positively and negatively associated with PC1, whereas *Plagl1* and *Cd3e* were the most positively and negatively PC2-associated genes. Thus, albeit not differentially expressed in pairwise comparisons between the groups, these genes substantially contributed to the study variance observed in general, likely reflecting their potential relation to affective and inflammatory pathogenesis in the PCUS model used here.

**Figure 5.**
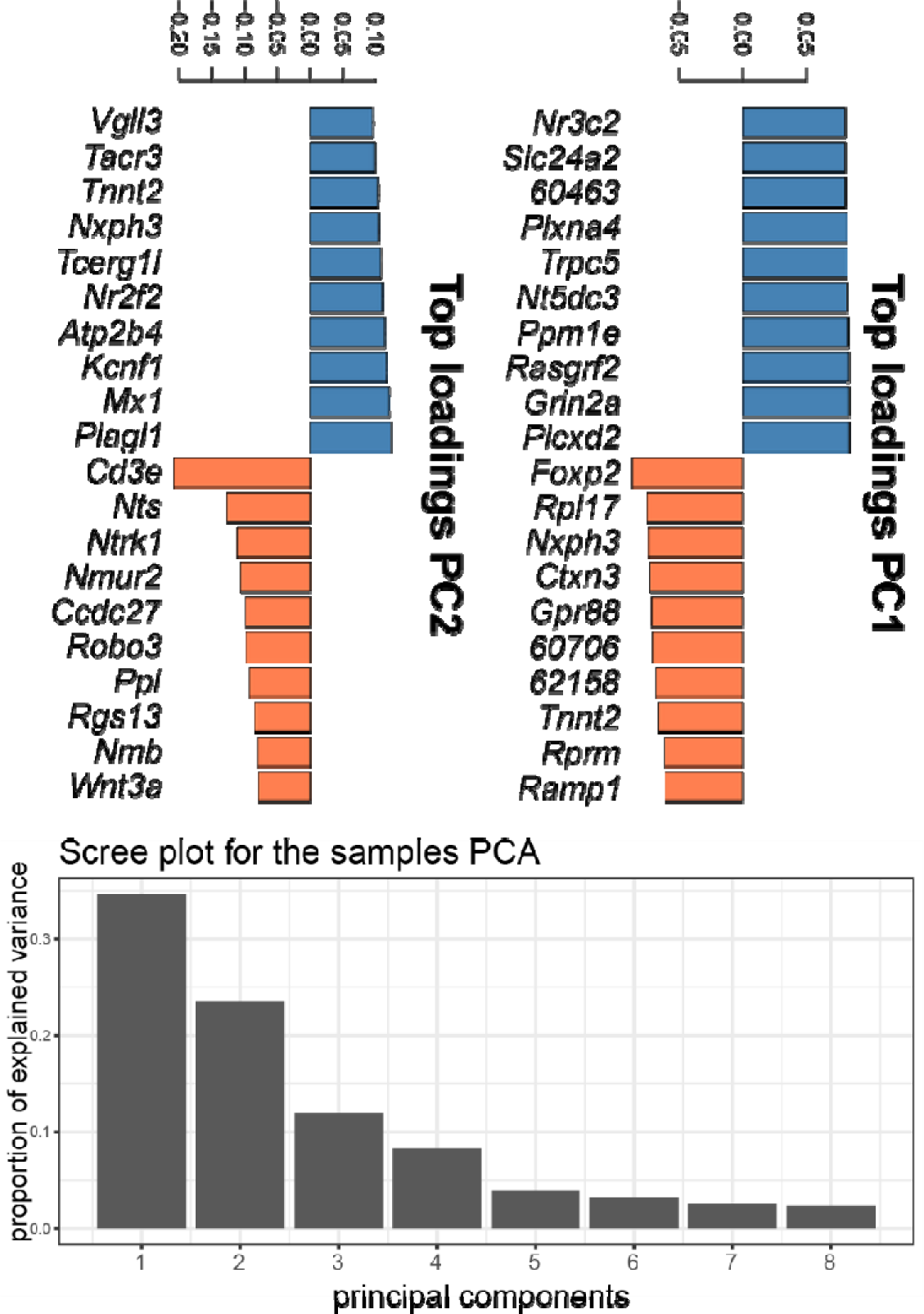
Summary of top 10 up- and down-regulated genes in rat hippocampal samples for Principal Component (PC) 1 and 2 loadings based on Principal Components Analysis (PCA), reflecting most variance in gene expression observed in the study across all rat groups. Inset: proportion of variance observed in the study explained by each PC. PC1 and PC2 together explain 59% of the sample variance observed in the study. Two genes lacking common names are shows as the last 5 digits of their Ensembl ID for *Rattus norvegicus*.

Our hippocampal transcriptomic analyses yielded 361 DE genes in PCUS, 12 in fluoxetine-, 4 in LPS-, 348 in EPA-exposed rats, as well as 1 DE gene (*Cga*) in fluoxetine+LPS and 452 in fluoxetine+EPA groups (q<0.017 vs. controls, Fig. 6 and Supplementary Table S3). The three DE genes (*Gpr6*, *Drd2* and *Adora2a* that encode G protein-coupled receptor 6, dopamine D2 receptor and adenosine A2A receptor, respectively), were downregulated vs. controls in all analyses (except fluoxetine+LPS). There were also 81 DE genes vs. PCUS for fluoxetine, 8 for EPA-, 40 for LPS-, 4 for fluoxetine+LPS- and 299 for fluoxetine+EPA-exposed rats (q<0.02, Fig. 6 and Supplementary Table S3).

**Figure 6.**
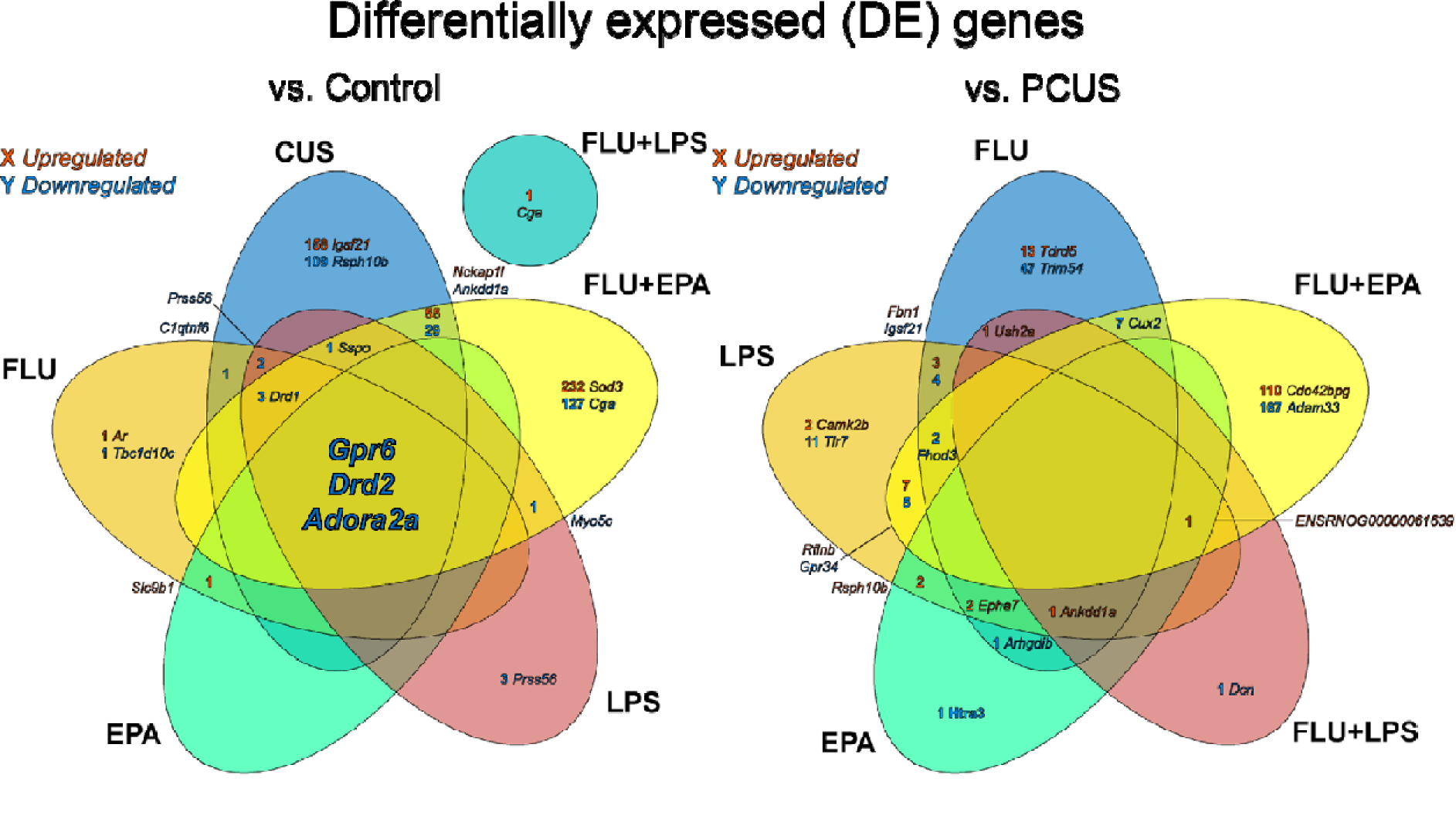
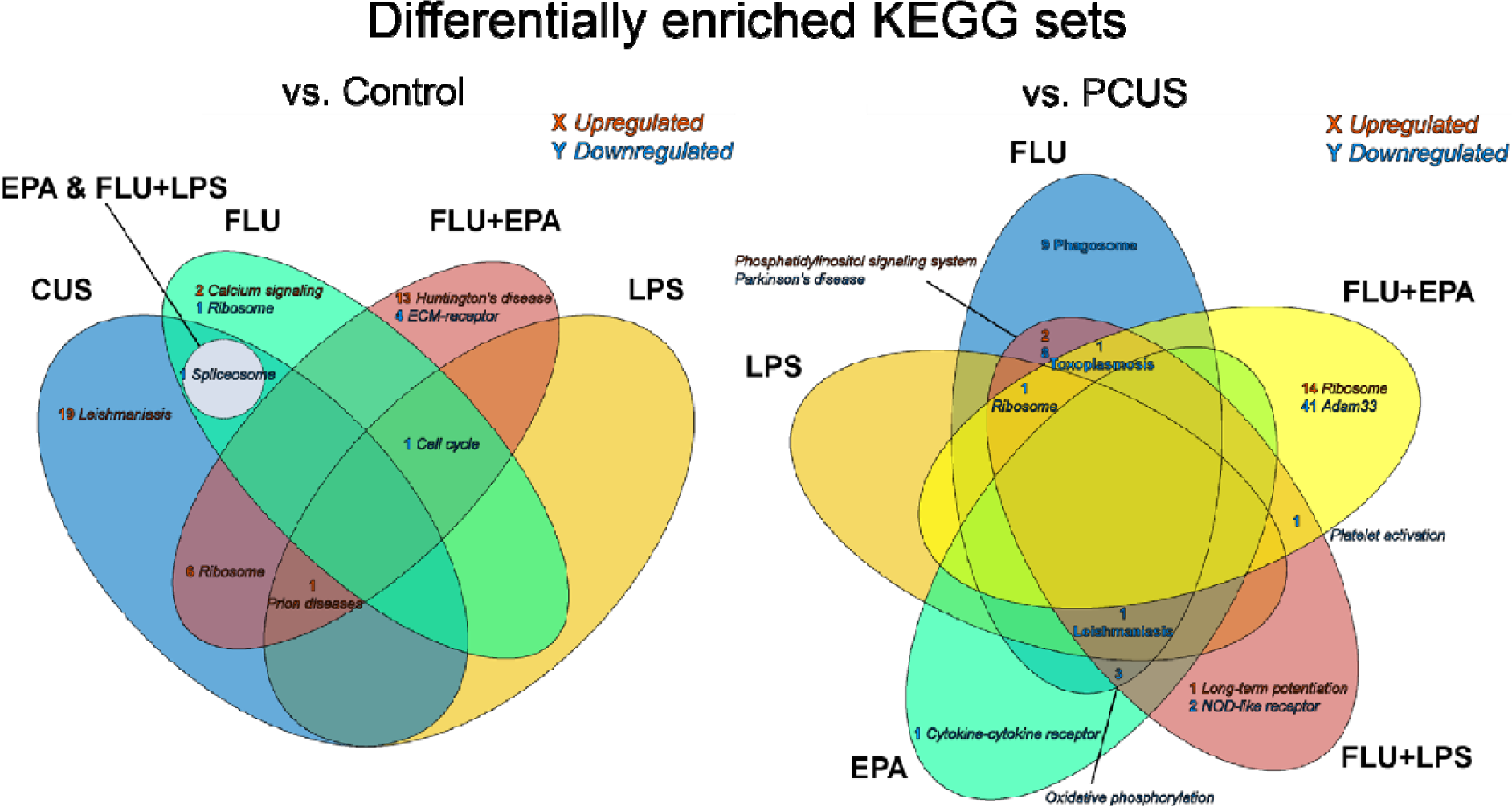
Venn’s diagrams illustrating differentially expressed (DE) genes (top panels) and differentially enriched KEGG gene sets (bottom panels) vs. control (q<0.016) and prolonged chronic unpredictable stress (PCUS) group (q<0.02). FLU – fluoxetine, EPA - eicosapentaenoic acid, LPS – lipopolysaccharide. Numbers correspond to the number of up- or down-regulated genes for the corresponding overlapping groups. Gene names reflect selected representative up- or down-regulated genes with significant changes in expression shared between the corresponding groups. Diagrams were constructed using the VennDiagram R package ^101^. Note that *Gpr6*, *Drd2* and *Adora2a* were downregulated in stressed rats treated with fluoxetine+EPA, suggesting G protein-coupled receptor 6, dopamine D2 receptor and adenosine A2A receptor as potential evolutionarily conserved targets in chronic stress.

GSEA identified 27 enriched sets in PCUS, 5 in fluoxetine-, 2 in LPS- and 26 in fluoxetine+EPA-treated groups, with only one (*rno03040 Spliceosome*) gene downregulated in both the EPA and fluoxetine+LPS groups vs. control (q<0.017). We also found 25 enriched sets in fluoxetine, 2 in LPS, 5 in EPA, 20 in fluoxetine+LPS and 58 in fluoxetine+EPA rats vs. PCUS group (q<0.02, Fig. 7 and Supplementary Table S4).

## Discussion

The present study, for the first time, developed and applied the 12-week PCUS protocol as a clinically relevant chronic stress model in rats, adapting it from our recently validated zebrafish PCUS model^38^, and assessing a wide range of behavioral and hippocampal transcriptomic responses. In general, PCUS recapitulated in rats pronounced and stable behavioral and molecular phenotypes induced by chronic stress clinically, and is also consistent with overt impact on behavior and neurochemistry produced by PCUS in zebrafish^38^. Collectively, this strongly supports the value and importance of cross-species translation between chronic stress models and calls for further studies comparing CUS models between a wide range of model organisms.

The study also combined PCUS with a well-established pro-inflammatory agent LPS (known to promote stress responses in various animal models and clinically), and observed similar LPS-potentiated PCUS effects in rat OFT and EPMT behaviors (Fig. 2-3). We also explored potential pharmacological and alternative (e.g., dietary) therapies in the rat PCUS model, applying a widely used, clinically active conventional antidepressant fluoxetine and a putative novel (EPA) treatment, noting some evidence of their potentially synergistic anti-stress effects.

On the one hand, these findings seem to parallel other rodent chronic stress models^108,109^ and phenotypes observed in clinical patients^110^, hence supporting the translational validity of the present rat PCUS model. Likewise, PCUS increased rat self-grooming, suggesting increased anxiety in stressed rats, similar to ^69,111^. Interestingly, EPA (but not fluoxetine) reduced tail- and head grooming vs. stress group (Fig. 4), suggesting that while classical anxiety-like behaviors (e.g., rearing and freezing) and anxiety-related grooming activity correlate, they may be differentially affected by some treatments, likely reflecting distinct (i.e., exploratory vs. displacement activity) behaviors and, possibly, CNS circuits.

On the other hand, chronic fluoxetine reversed most of anxiogenic-like behavioral effects of PCUS in rats (Fig. 2-3). Generally in line with anxiolytic^112^, antidepressant^113^ and anti-inflammatory^114^ effects observed for fluoxetine and other selective serotonin reuptake inhibitors (SSRIs) clinically. This anti-stress CNS profile also resembles the effects of fluoxetine in other rodent stress studies, including modulating anxiety in various, albeit shorter, chronic stress protocols^51,115–117^. Collectively, these findings further support face validity of the rat PCUS model used here.

Similarly, although EPA did not rescue anxiety-like OFT behavior, it was effective in both EPMT and GT assays. Fluoxetine+EPA was more effective in the OFT, rescuing stress-evoked rearing (but not freezing) and EPMT anxiety-like behaviors. These results are in line with some clinical studies on positive effects of EPA in depression^118,119^, but also suggest their potential synergism, paralleling recent clinical evidence that combining fluoxetine with EPA may be more efficient than monotherapy^120^. This is also consistent with our earlier zebrafish PCUS data on lesser anti-stress efficiency of EPA compared to its combination with fluoxetine^38^. Because EPA is a critical PUFA with multiple physiological functions in vivo, including anti-inflammation beneficial in various psychiatric disorders^121,122^, further studies are needed to better understand potentially synergistic (and, seemingly shared across taxa) effects of fluoxetine and EPA in the PCUS models.

As a key component of gram-negative bacterial membrane, LPS triggers inflammation via immune and non-immune mechanisms^123^, promoting the release of pro-inflammatory cytokines interleukin (IL) IL-1β and tumor necrosis factor-β (TNF-β)^124^. Proinflammatory cytokines are often associated with affective disorders^125^, suggesting that combining stress with LPS by promoting anxiety-like phenotype (as seen for some behavioral endpoints here) can be used for studying stress-neuroimmune interplay in various experimental models.

In addition to behavioral deficits, PCUS caused pronounced transcriptomic changes in 361 rat hippocampal genes and 27 changes in gene set enrichment, that were further corrected by fluoxetine. Altered expression of G protein-coupled receptor (GPCR)-related genes may be particularly interesting here, given their wide use as molecular targets for pharmacological interventions^126^. Hippocampi of PCUS-exposed rats over-expressed *Gpr84* (that controls the levels of inflammatory mediators) ^127,128^ and downregulated *Gpr176* (whose protein inhibits cAMP signaling)^129^ and *Gpr6* (associated with sphingosine-1-phosphate signaling ^130^, learning^131^, neurite outgrowth^132^ and other CNS diosrders^133^). Both animal^134,135^ and human studies^136,137^ implicate dopamine D2 receptor and its gene (*Drd2*) in anxiety and depression, and adenosine A2A receptor gene *(Adora2a*) in the regulation of glutamate and dopamine release, and suggest *Drd2* as a potential therapeutic target for the treatment of various CNS conditions, including insomnia, pain, depression and Parkinson’s disease^138,139^.

Furthermore, PCUS-exposed rats also showed upregulated expression of the immunoglobulin superfamily member 2 (*Igsf2*) and NCK Associated Protein 1 Like (*Nckap1l*) genes that can modulate CNS functions^140,141^, as well as the expression of Kinesin-like protein 17 (*Kif17*) gene involved in microtubule transport of the N-methyl-D-aspartate (NMDA) receptor subunit Nr2b (*Grin2b*) in hippocampal neurons^142,143^. PCUS also increased enrichment of multiple disease (e.g. *rno05152 Tuberculosis*, *rno05323 Rheumatoid arthritis*) and inflammation-related pathways (e.g. *rno04060 Cytokine-cytokine receptor interaction*), supporting the stress-inflammation interplay observed both in animal models of affective pathologies and clinically ^144–146^. PCUS also reduced enrichment of *rno03040 Spliceosome* and increased it for *rno03010 Ribosome,* implying altered protein synthesis in this model. Collectively, this indicates robust neurotranscriptomic responses to PCUS in rats.

Importantly, fluoxetine corrected most behavioral and transcriptome alterations induced by PCUS in rats, also decreasing the expression of the Chemerin Chemokine-Like Receptor 1 (*Cmklr1*) gene (linked to inflammation and depression^147^), hence further linking neuroinflammation and affective pathogenesis in chronic stress. Interestingly, while successfully rescuing most of gene set enrichment profiles, it did not alter protein synthesis genes, with both *rno03040 Spliceosome* and *rno03010 Ribosome* being downregulated. Fluoxetine also increased enrichment of *rno04020 Calcium signaling pathway* and *rno04724 Glutamatergic synapse*, the effect not observed in other groups, and hence likely reflecting specific positive mechanisms of this drug in the PCUS model. For several genes mentioned earlier (*Gpr6*, *Drd2* and *Adora2a*), their expression was reduced in most groups vs. control (Supplementary Table S3). Such shared expression pattern seems to support the common ‘core’ (and, hence, likely evolutionarily conserved), role of genes that remained treatment-resistant here, and therefore merit scrutiny in clinical cohorts with treatment-resistant affective pathologies.

Both EPA and fluoxetine upregulated the expression of Solute Carrier Family 9 Member B1 (*Slc9b1*) gene that regulates DNA methylation^148^ and is linked to various stress-related brain disorders^149–151^. As both drugs treat clinical depression, and their combination is more efficient than monotherapy^152,153^, rats treated with fluoxetine+EPA show upregulated brain Superoxide Dismutase 3 (*Sod3*) gene involved in anti-inflammation and neuroprotection in stress^154^. Both fluoxetine and fluoxetine+EPA reduced hippocampal expression of Cut Like Homeobox 2 (*Cux2)* that regulates the development of dendrites, dendritic spines and synapses of neocortical neurons in mice, and has been linked to clinical affective disorders and schizophrenia^155,156^. Unlike EPA, LPS initiates an inflammatory response via both immune and non-immune factors^123^. Finally, LPS downregulated (vs. PCUS) the expression of Toll-like receptor 7 (*Tlr7*) that modulates neurodevelopment and brain functions^157^, and Calcium/Calmodulin Dependent Protein Kinase II Beta (*Camk2b*; vs. chronic stress), a gene implicated in clinical depression^158^. Overall, these findings suggest that LPS treatment may worsen on neurogenomic level the physiological impact of chronic stress.

Nevertheless, there were also several limitations of the present study. For example, our transcriptomic analyses focused on one brain region (hippocampus), unlike some multi-region neurotranscriptomic studies. Furthermore, like other model organisms, rodent display intra-species variation^159–161^, including behavioral sex differences^41,162–164^, that may play a role in stress mechanisms and effects on pharmacologically evoked phenotypes as well. Although assessing intraspecies variability was outside the scope of the present study, this line of research clearly merits further scrutiny in subsequent follow-up studies.

In summary, the rat PCUS protocol developed here induced overt anxiety-like behavioral effects rescued by fluoxetine and, partially, EPA. Fluoxetine recovered most of PCUS-evoked behavioral and molecular alterations when administered alone or in combination with LPS and especially with EPA. Chronic stress robustly affected rat brain transcriptomic profiles, downregulating hippocampal *Gpr6*, *Drd2* and *Adora2a* genes in most treatment groups, suggesting conserved role of these genes in chronic stress pathogenesis.

## Acknowledgements

This work was financially supported by the Ministry of Science and Higher Education of the Russian Federation (Agreement No. 075-15-2020-901). AVK is the Chair of the International Zebrafish Neuroscience Research Consortium (ZNRC) and President of the International Stress and Behavior Society (ISBS, www.stressandbehavior.com) that coordinated this collaborative multi-laboratory project. The consortium provided a collaborative idea exchange platform for this study. It is not considered as affiliation and did not fund the study. This study utilized equipment of the Core Facilities Centre “Centre for Molecular and Cell Technologies” of St. Petersburg State University. The funders had no role in the design, analyses, and interpretation of the submitted study, or decision to publish.

## Data availability

The datasets generated and/or analyzed during the current study are available from the GEO repository (GSE205325) or from the corresponding authors upon reasonable request.

## Ethical statement

All experimental animal manipulations were approved by the Ethics committee of the Institute of Experimental Medicine at Almazov National Medical Research Center (approval number 20-14PZ#V2). All animals tested were included in final analyses, without removing the outliers. All experiments were performed as planned, and all analyses and endpoints assessed were included without omission. The study experimental design and its description here, as well as data analysis and presenting, adhered to the ARRIVE guidelines for reporting animal research and the PREPARE guidelines for planning animal research and testing, as well as the 3Rs principles of humane animal experimentation.

## Conflict of interest statement

The authors declare no conflict of interest.

## CRediT authorship contribution statement

Konstantin A. Demin (Conceptualization) (Data curation) (Formal analysis) (Funding acquisition) (Investigation) (Methodology) (Project administration) (Resources) (Software) (Validation) (Visualization) (Writing - original draft) (Writing - review and editing), Tatiana O. Kolesnikova (Investigation) (Methodology) (Resources) (Data curation) (Formal analysis) (Writing - original draft) (Writing - review and editing), David S. Galstyan (Investigation) (Resources) (Data curation) (Writing - original draft) (Writing - review and editing), Nataliya A. Krotova (Investigation) (Writing - original draft) (Writing - review and editing), Nikita P. Ilyin (Investigation) (Writing - original draft) (Methodology) (Writing - review and editing), Ksenia A. Derzhavina (Investigation) (Writing - original draft) (Methodology) (Writing - review and editing), Maria Seredinskaya (Investigation) (Writing - original draft) (Methodology) (Writing - review and editing), Yuriy M. Kositsyn (Investigation) (Writing - original draft) (Writing - review and editing), Dmitry V. Sorokin (Investigation) (Writing - original draft) (Writing - review and editing), Maria O. Nerush (Investigation) (Writing - original draft) (Writing - review and editing), Abubakar-Askhab S. Khaybaev (Investigation) (Writing - original draft) (Writing - review and editing), Sofia A. Pushkareva (Investigation) (Writing - original draft) (Writing - review and editing), Elena Petersen (Writing - original draft) (Writing - review and editing), Murilo S. de Abreu (Writing - original draft) (Writing - review and editing), Alexey Masharsky (Investigation) (Writing - original draft) (Writing - review and editing), Allan V. Kalueff (Conceptualization) (Funding acquisition) (Methodology) (Project administration) (Resources) (Supervision) (Validation) (Visualization) (Writing - review and editing)

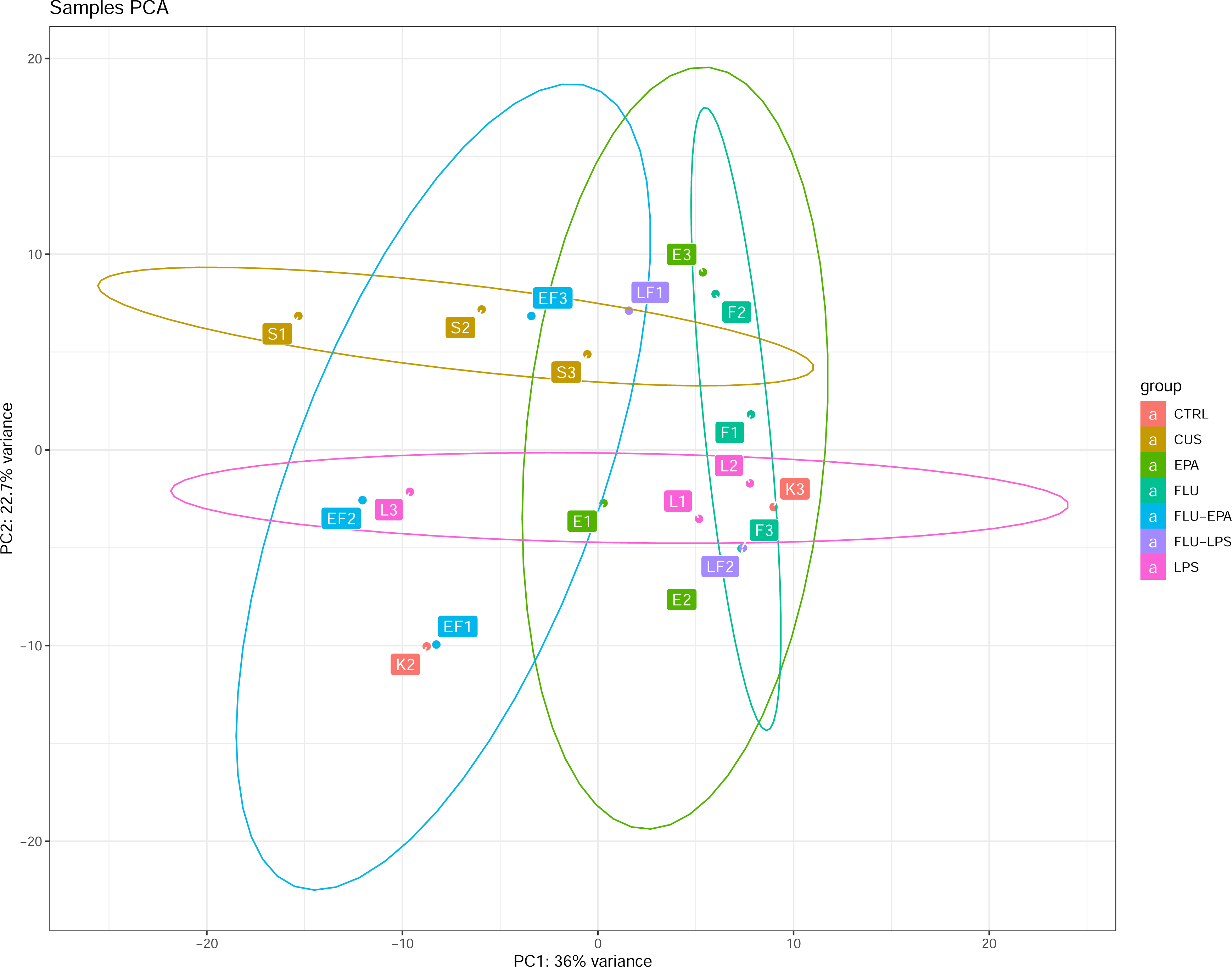

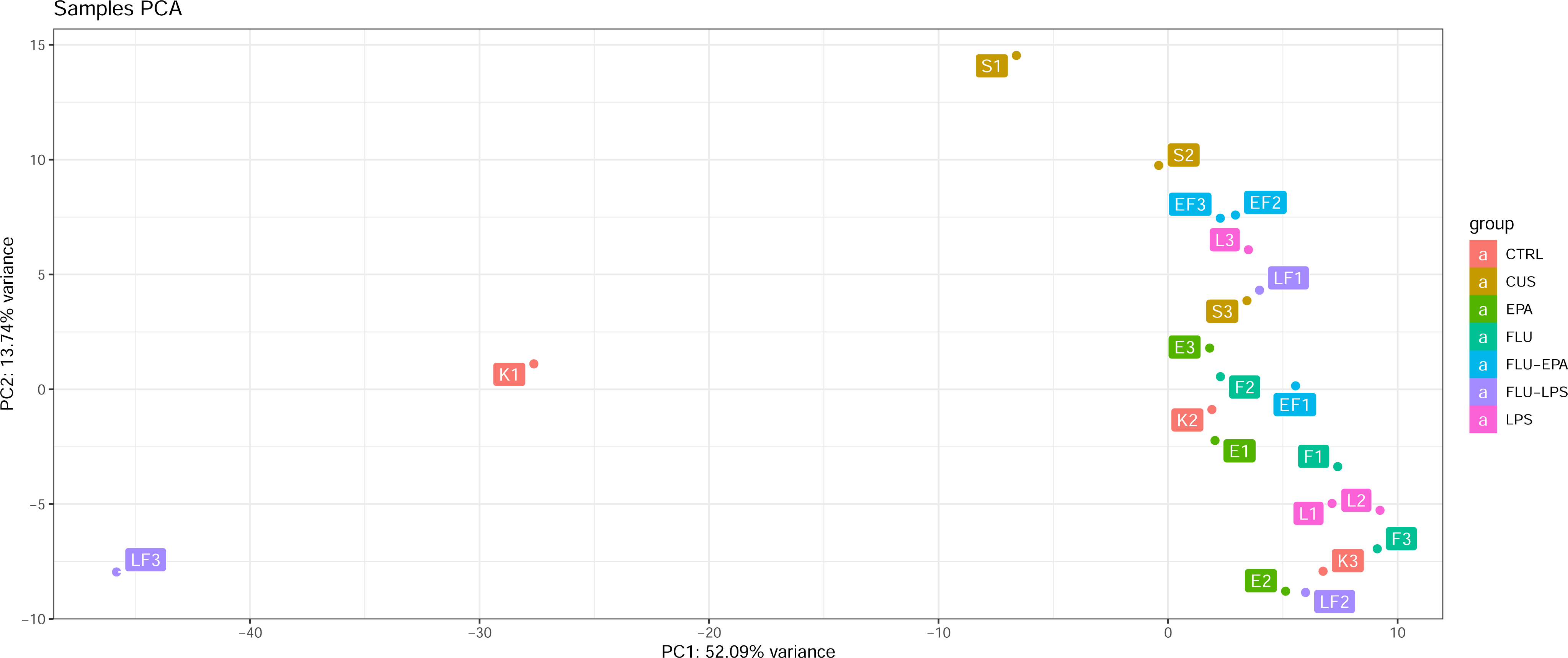

## Notes

https://www.ncbi.nlm.nih.gov/geo/query/acc.cgi?acc=GSE205325

### Competing Interest Statement

The authors have declared no competing interest.

### Summary of Updates

Minor revisions for streamlining the text and improving explanation of specific details.

