## Supplementary Tables for "Effects of chronic exposure to fluoxetine, eicosapentaenoic acid, and lipopolysaccharide on behavior and hippocampal transcriptome in the rat model of prolonged chronic unpredictable stress"

**Supplementary Table S1.** Results of Generalized Linear Model (GZLM) fits using group, testing day and their interaction effects as ‘predictors’, to compare behavior in experimental groups. The corrected Akaike information criterion (AICc) was used to choose the ‘best fit’ model among Gaussian distribution (identity link), Poisson distribution (with log link), Gamma distribution (inverse and log links) and Inverse Gaussian distribution (with inverse and log links) for the Open Field Test, Elevated Plus-Maze Test and Grooming Test. CUS – chronic unpredictable stress, FLU – fluoxetine, EPA - eicosapentaenoic acid, LPS – lipopolysaccharide, FLUE – fluoxetine and eicosapentaenoic acid, FLUL – fluoxetine and lipopolysaccharide.

| **Open Field Test** | | | | |
| --- | --- | --- | --- | --- |
| **Rearing frequency, n** | **Inversed Gauss distribution log link** | | | |
| **predictor** | **estimate** | **Ci** | **statistic** | **p.value** |
| **Intercept** | **1.00** | **0.58; 1.42** | **4.63** | **< .001** |
| Day2 | 0.00 | -0.72; 0.72 | 0.00 | > .999 |
| **GroupStress** | **-0.52** | **-1.02; -0.03** | **-2.06** | **.041** |
| **GroupFlu** | **-0.59** | **-1.09; -0.09** | **-2.30** | **.023** |
| **GroupEPA** | **-0.72** | **-1.22; -0.23** | **-2.88** | **.005** |
| **GroupLPS** | **-0.72** | **-1.20; -0.24** | **-2.96** | **.004** |
| **GroupFLUE** | **-0.94** | **-1.38; -0.50** | **-4.23** | **< .001** |
| **GroupFLUL** | **-0.69** | **-1.21; -0.18** | **-2.63** | **.010** |
| Day2 x GroupStress | -0.37 | -1.18; 0.44 | -0.90 | .372 |
| Day2 x GroupFlu | 0.59 | -0.49; 1.67 | 1.07 | .287 |
| Day2 x GroupEPA | -0.13 | -0.92; 0.67 | -0.31 | .757 |
| Day2 x GroupLPS | 0.72 | -0.35; 1.79 | 1.33 | .188 |
| **Day2 x GroupFLUE** | **0.94** | **0.04; 1.84** | **2.05** | **.043** |
| Day2 x GroupFLUL | -0.27 | -1.05; 0.51 | -0.69 | .492 |
| **Rearing duration, s** | **Inversed Gauss distribution log link** | | | |
| **predictor** | **estimate** | **Ci** | **statistic** | **p.value** |
| **Intercept** | **1.00** | **0.58; 1.42** | **4.63** | **< .001** |
| Day2 | 0.00 | -0.72; 0.72 | 0.00 | > .999 |
| **GroupStress** | **-0.52** | **-1.02; -0.03** | **-2.06** | **.041** |
| **GroupFlu** | **-0.59** | **-1.09; -0.09** | **-2.30** | **.023** |
| **GroupEPA** | **-0.72** | **-1.22; -0.23** | **-2.88** | **.005** |
| **GroupLPS** | **-0.72** | **-1.20; -0.24** | **-2.96** | **.004** |
| **GroupFLUE** | **-0.94** | **-1.38; -0.50** | **-4.23** | **< .001** |
| **GroupFLUL** | **-0.69** | **-1.21; -0.18** | **-2.63** | **.010** |
| Day2 x GroupStress | -0.37 | -1.18; 0.44 | -0.90 | .372 |
| Day2 x GroupFlu | 0.59 | -0.49; 1.67 | 1.07 | .287 |
| Day2 x GroupEPA | -0.13 | -0.92; 0.67 | -0.31 | .757 |
| Day2 x GroupLPS | 0.72 | -0.35; 1.79 | 1.33 | .188 |
| **Day2 x GroupFLUE** | **0.94** | **0.04; 1.84** | **2.05** | **.043** |
| Day2 x GroupFLUL | -0.27 | -1.05; 0.51 | -0.69 | .492 |
| **Climbing frequency, n** | **Inversed Gamma distribution identity link** | | | |
| **predictor** | **estimate** | **ci** | **statistic** | **p.value** |
| **Intercept** | **0.12** | **0.09; 0.16** | **6.62** | **< .001** |
| Day2 | 0.05 | -0.02; 0.14 | 1.35 | .179 |
| GroupStress | -0.02 | -0.07; 0.02 | -1.07 | .287 |
| GroupFlu | -0.03 | -0.08; 0.01 | -1.35 | .178 |
| GroupEPA | -0.02 | -0.07; 0.04 | -0.59 | .558 |
| GroupLPS | -0.03 | -0.08; 0.01 | -1.29 | .199 |
| GroupFLUE | -0.04 | -0.08; 0.01 | -1.68 | .095 |
| GroupFLUL | -0.04 | -0.09; 0.00 | -1.89 | .061 |
| Day2 x GroupStress | -0.01 | -0.13; 0.11 | -0.24 | .809 |
| Day2 x GroupFlu | -0.07 | -0.17; 0.02 | -1.52 | .130 |
| Day2 x GroupEPA | -0.04 | -0.15; 0.05 | -0.86 | .393 |
| Day2 x GroupLPS | 0.02 | -0.10; 0.15 | 0.34 | .735 |
| Day2 x GroupFLUE | -0.02 | -0.12; 0.07 | -0.45 | .656 |
| Day2 x GroupFLUL | -0.02 | -0.12; 0.07 | -0.41 | .682 |
| **Climbing duration, s** | **Gauss distribution identity link** | | | |
| **predictor** | **estimate** | **ci** | **statistic** | **p.value** |
| **Intercept** | **1.00** | **0.58; 1.42** | **4.63** | **< .001** |
| Day2 | 0.00 | -0.72; 0.72 | 0.00 | > .999 |
| **GroupStress** | **-0.52** | **-1.02; -0.03** | **-2.06** | **.041** |
| **GroupFlu** | **-0.59** | **-1.09; -0.09** | **-2.30** | **.023** |
| **GroupEPA** | **-0.72** | **-1.22; -0.23** | **-2.88** | **.005** |
| **GroupLPS** | **-0.72** | **-1.20; -0.24** | **-2.96** | **.004** |
| **GroupFLUE** | **-0.94** | **-1.38; -0.50** | **-4.23** | **< .001** |
| **GroupFLUL** | **-0.69** | **-1.21; -0.18** | **-2.63** | **.010** |
| Day2 x GroupStress | -0.37 | -1.18; 0.44 | -0.90 | .372 |
| Day2 x GroupFlu | 0.59 | -0.49; 1.67 | 1.07 | .287 |
| Day2 x GroupEPA | -0.13 | -0.92; 0.67 | -0.31 | .757 |
| Day2 x GroupLPS | 0.72 | -0.35; 1.79 | 1.33 | .188 |
| **Day2 x GroupFLUE** | **0.94** | **0.04; 1.84** | **2.05** | **.043** |
| Day2 x GroupFLUL | -0.27 | -1.05; 0.51 | -0.69 | .492 |
| **Vertical activity frequency, n** | **Inversed Gamma distribution identity link** | | | |
| **predictor** | **estimate** | **ci** | **statistic** | **p.value** |
| **Intercept** | **0.10** | **0.07; 0.13** | **6.49** | **< .001** |
| Day2 | 0.03 | -0.03; 0.10 | 0.95 | .346 |
| GroupStress | -0.01 | -0.05; 0.03 | -0.51 | .611 |
| GroupFlu | -0.02 | -0.06; 0.02 | -0.86 | .392 |
| GroupEPA | 0.00 | -0.04; 0.05 | 0.09 | .931 |
| GroupLPS | -0.02 | -0.06; 0.01 | -1.16 | .248 |
| GroupFLUE | -0.03 | -0.07; 0.01 | -1.41 | .161 |
| GroupFLUL | -0.03 | -0.07; 0.01 | -1.55 | .125 |
| Day2 x GroupStress | 0.01 | -0.09; 0.13 | 0.20 | .839 |
| Day2 x GroupFlu | -0.06 | -0.14; 0.01 | -1.73 | .087 |
| Day2 x GroupEPA | -0.04 | -0.12; 0.04 | -0.87 | .388 |
| Day2 x GroupLPS | 0.04 | -0.06; 0.15 | 0.78 | .440 |
| Day2 x GroupFLUE | -0.01 | -0.09; 0.06 | -0.36 | .721 |
| Day2 x GroupFLUL | -0.02 | -0.09; 0.05 | -0.46 | .648 |
| **Vertical activity duration, s** | **Gauss distribution identity link** | | | |
| **predictor** | **estimate** | **ci** | **statistic** | **p.value** |
| **Intercept** | **17.03** | **11.16; 22.90** | **5.69** | **< .001** |
| Day2 | -5.38 | -15.30; 4.54 | -1.06 | .290 |
| GroupStress | 1.31 | -6.59; 9.22 | 0.33 | .745 |
| GroupFlu | 0.39 | -7.91; 8.69 | 0.09 | .927 |
| GroupEPA | 0.17 | -8.73; 9.07 | 0.04 | .971 |
| GroupLPS | 5.02 | -3.28;13.32 | 1.19 | .238 |
| GroupFLUE | 8.26 | -0.22; 16.73 | 1.91 | .059 |
| GroupFLUL | 8.70 | -0.81; 18.21 | 1.79 | .076 |
| Day2 x GroupStress | 0.20 | -16.41; 16.80 | 0.02 | .981 |
| **Day2 x GroupFlu** | **20.12** | **4.47; 35.77** | **2.52** | **.013** |
| Day2 x GroupEPA | 6.50 | -7.89; 20.90 | 0.89 | .378 |
| Day2 x GroupLPS | -3.84 | -19.49; 11.81 | -0.48 | .632 |
| Day2 x GroupFLUE | -2.84 | -16.69; 11.01 | -0.40 | .688 |
| Day2 x GroupFLUL | 4.43 | -10.07; 18.94 | 0.60 | .550 |
| **Freezing frequency, n** | **Inversed Gauss distribution identity link** | | | |
| **predictor** | **estimate** | **ci** | **statistic** | **p.value** |
| **Intercept** | **1.00** | **0.72; 1.28** | **7.08** | **< .001** |
| Day2 | 0.00 | -0.47; 0.47 | 0.00 | > .999 |
| GroupStress | -0.27 | -0.62; 0.08 | -1.53 | .129 |
| GroupFlu | -0.35 | -0.71; 0.01 | -1.93 | .056 |
| GroupEPA | -0.23 | -0.62; 0.16 | -1.15 | .251 |
| GroupLPS | -0.19 | -0.56; 0.19 | -0.99 | .326 |
| **GroupFLUE** | **-0.52** | **-0.86; -0.18** | **-2.98** | **.003** |
| GroupFLUL | -0.38 | -0.78; 0.01 | -1.92 | .057 |
| Day2 x GroupStress | -0.39 | -1.01; 0.22 | -1.26 | .210 |
| Day2 x GroupFlu | 0.35 | -0.37; 1.07 | 0.95 | .343 |
| Day2 x GroupEPA | -0.13 | -0.75; 0.49 | -0.42 | .676 |
| Day2 x GroupLPS | 0.19 | -0.54; 0.92 | 0.50 | .615 |
| Day2 x GroupFLUE | 0.52 | -0.10; 1.14 | 1.65 | .103 |
| Day2 x GroupFLUL | -0.25 | -0.84; 0.33 | -0.84 | .400 |
| **Freezing duration, s** | **Inversed Gauss distribution identity link** | | | |
| **predictor** | **estimate** | **ci** | **statistic** | **p.value** |
| **Intercept** | **1.00** | **0.58; 1.42** | **4.63** | **< .001** |
| Day2 | 0.00 | -0.72; 0.72 | 0.00 | > .999 |
| **GroupStress** | **-0.52** | **-1.02; -0.03** | **-2.06** | **.041** |
| **GroupFlu** | **-0.59** | **-1.09; -0.09** | **-2.30** | **.023** |
| **GroupEPA** | **-0.72** | **-1.22; -0.23** | **-2.88** | **.005** |
| **GroupLPS** | **-0.72** | **-1.20; -0.24** | **-2.96** | **.004** |
| **GroupFLUE** | **-0.94** | **-1.38; -0.50** | **-4.23** | **< .001** |
| **GroupFLUL** | **-0.69** | **-1.21; -0.18** | **-2.63** | **.010** |
| Day2 x GroupStress | -0.37 | -1.18; 0.44 | -0.90 | .372 |
| Day2 x GroupFlu | 0.59 | -0.49; 1.67 | 1.07 | .287 |
| Day2 x GroupEPA | -0.13 | -0.92; 0.67 | -0.31 | .757 |
| Day2 x GroupLPS | 0.72 | -0.35;1.79 | 1.33 | .188 |
| **Day2 x GroupFLUE** | **0.94** | **0.04; 1.84** | **2.05** | **.043** |
| Day2 x GroupFLUL | -0.27 | -1.05; 0.51 | -0.69 | .492 |
| **Elevated plus-maze test** | | | | |
| **Closed Arm frequency, n** | **Inversed Gauss distribution log link** | | | |
| **predictor** | **estimate** | **ci** | **statistic** | **p.value** |
| **Intercept** | **1.41** | **1.19; 1.68** | **11.60** | **< .001** |
| Day2 | 0.18 | -0.23; 0.66 | 0.81 | .422 |
| **GroupStress** | **-0.39** | **-0.71; -0.11** | **-2.61** | **.010** |
| GroupFlu | 0.04 | -0.31; 0.39 | 0.21 | .831 |
| GroupEPA | 0.08 | -0.29; 0.47 | 0.41 | .685 |
| **GroupLPS** | **0.45** | **0.10; 0.82** | **2.52** | **.013** |
| GroupFLUE | 0.30 | -0.07; 0.70 | 1.58 | .118 |
| **GroupFLUL** | **0.60** | **0.18; 1.13** | **2.61** | **.010** |
| Day2 x GroupStress | -0.09 | -0.71; 0.59 | -0.28 | .783 |
| Day2 x GroupFlu | -0.44 | -1.07; 0.21 | -1.39 | .169 |
| Day2 x GroupEPA | -0.38 | -1.01; 0.21 | -1.27 | .207 |
| **Day2 x GroupLPS** | **-0.93** | **-1.57; -0.29** | **-2.96** | **.004** |
| Day2 x GroupFLUE | -0.49 | -1.12; 0.10 | -1.63 | .106 |
| Day2 x GroupFLUL | -0.42 | -1.16; 0.32 | -1.16 | .250 |
| **Closed Arm duration, s** | **Gauss distribution identity link** | | | |
| **predictor** | **estimate** | **ci** | **statistic** | **p.value** |
| **Intercept** | **256.87** | **237.31; 276.43** | **25.74** | **< .001** |
| Day2 | 2.10 | -30.96; 35.16 | 0.12 | .901 |
| GroupStress | 5.69 | -20.64; 32.01 | 0.42 | .673 |
| GroupFlu | 8.68 | -18.98; 36.33 | 0.61 | .540 |
| GroupEPA | -4.25 | -33.91; 25.41 | -0.28 | .779 |
| GroupLPS | -19.05 | -45.03; 6.93 | -1.44 | .153 |
| GroupFLUE | 5.47 | -22.76; 33.69 | 0.38 | .705 |
| **GroupFLUL** | **-40.09** | **-70.67; -9.52** | **-2.57** | **.011** |
| Day2 x GroupStress | 16.96 | -38.37; 72.28 | 0.60 | .549 |
| Day2 x GroupFlu | -12.66 | -64.79; 39.48 | -0.48 | .635 |
| Day2 x GroupEPA | 1.02 | -46.94;48.98 | 0.04 | .967 |
| Day2 x GroupLPS | 44.53 | -6.74; 95.79 | 1.70 | .091 |
| Day2 x GroupFLUE | -7.18 | -53.31; 38.96 | -0.30 | .761 |
| **Day2 x GroupFLUL** | **56.99** | **7.25; 106.73** | **2.25** | **.027** |
| **Open Arm frequency, n** | **Inversed Gauss distribution identity link** | | | |
| **predictor** | **estimate** | **ci** | **statistic** | **p.value** |
| **Intercept** | **0.43** | **0.27; 0.60** | **5.12** | **< .001** |
| Day2 | 0.00 | -0.28; 0.29 | 0.03 | .977 |
| GroupStress | 0.14 | -0.10; 0.38 | 1.13 | .259 |
| GroupFlu | 0.03 | -0.21; 0.27 | 0.25 | .800 |
| GroupEPA | 0.07 | -0.20; 0.33 | 0.50 | .619 |
| GroupLPS | 0.04 | -0.19; 0.26 | 0.34 | .735 |
| GroupFLUE | 0.11 | -0.14; 0.37 | 0.86 | .391 |
| GroupFLUL | -0.17 | -0.40; 0.06 | -1.45 | .149 |
| Day2 x GroupStress | 0.42 | -0.20; 1.04 | 1.34 | .182 |
| Day2 x GroupFlu | -0.07 | -0.51; 0.37 | -0.31 | .760 |
| Day2 x GroupEPA | 0.03 | -0.39; 0.46 | 0.16 | .876 |
| Day2 x GroupLPS | 0.52 | -0.03; 1.08 | 1.85 | .067 |
| Day2 x GroupFLUE | 0.07 | -0.36; 0.49 | 0.30 | .762 |
| **Day2 x GroupFLUL** | **0.48** | **0.03; 0.94** | **2.07** | **.040** |
| **Open Arm duration, s** | **Inversed Gauss distribution log link** | | | |
| **predictor** | **estimate** | **ci** | **statistic** | **p.value** |
| **Intercept** | **256.87** | **237.31; 276.43** | **25.74** | **< .001** |
| Day2 | 2.10 | -30.96; 35.16 | 0.12 | .901 |
| GroupStress | 5.69 | -20.64; 32.01 | 0.42 | .673 |
| GroupFlu | 8.68 | -18.98;36.33 | 0.61 | .540 |
| GroupEPA | -4.25 | -33.91; 25.41 | -0.28 | .779 |
| GroupLPS | -19.05 | -45.03; 6.93 | -1.44 | .153 |
| GroupFLUE | 5.47 | -22.76; 33.69 | 0.38 | .705 |
| **GroupFLUL** | **-40.09** | **-70.67; -9.52** | **-2.57** | **.011** |
| Day2 x GroupStress | 16.96 | -38.37; 72.28 | 0.60 | .549 |
| Day2 x GroupFlu | -12.66 | -64.79; 39.48 | -0.48 | .635 |
| Day2 x GroupEPA | 1.02 | -46.94; 48.98 | 0.04 | .967 |
| Day2 x GroupLPS | 44.53 | -6.74; 95.79 | 1.70 | .091 |
| Day2 x GroupFLUE | -7.18 | -53.31; 38.96 | -0.30 | .761 |
| **Day2 x GroupFLUL** | **56.99** | **7.25; 106.73** | **2.25** | **.027** |
| **Freezing frequency, n** | **Inversed Gauss distribution identity link** | | | |
| **predictor** | **estimate** | **ci** | **statistic** | **p.value** |
| Intercept | 0.87 | 0.73; 1.00 | 12.70 | < .001 |
| Day2 | 0.13 | -0.10; 0.37 | 1.10 | .273 |
| GroupStress | 0.02 | -0.16; 0.20 | 0.24 | .810 |
| GroupFlu | 0.06 | -0.13; 0.25 | 0.63 | .530 |
| GroupEPA | 0.13 | -0.08; 0.34 | 1.24 | .219 |
| GroupLPS | 0.03 | -0.15; 0.21 | 0.31 | .759 |
| GroupFLUE | 0.13 | -0.07; 0.33 | 1.30 | .195 |
| GroupFLUL | 0.03 | -0.18; 0.24 | 0.31 | .758 |
| Day2 x GroupStress | -0.02 | -0.42; 0.38 | -0.11 | .914 |
| Day2 x GroupFlu | -0.26 | -0.62; 0.10 | -1.43 | .156 |
| Day2 x GroupEPA | -0.13 | -0.48; 0.22 | -0.75 | .455 |
| Day2 x GroupLPS | -0.69 | -1.00; -0.39 | -4.47 | < .001 |
| Day2 x GroupFLUE | -0.13 | -0.47; 0.20 | -0.78 | .437 |
| Day2 x GroupFLUL | -0.37 | -0.70; -0.03 | -2.14 | .035 |
| **Freezing duration, s** | **Inversed Gauss distribution identity link** | | | |
| **predictor** | **estimate** | **ci** | **statistic** | **p.value** |
| **Intercept** | **0.87** | **0.73; 1.00** | **12.70** | **< .001** |
| Day2 | 0.13 | -0.10; 0.37 | 1.10 | .273 |
| GroupStress | 0.02 | -0.16; 0.20 | 0.24 | .810 |
| GroupFlu | 0.06 | -0.13; 0.25 | 0.63 | .530 |
| GroupEPA | 0.13 | -0.08; 0.34 | 1.24 | .219 |
| GroupLPS | 0.03 | -0.15; 0.21 | 0.31 | .759 |
| GroupFLUE | 0.13 | -0.07; 0.33 | 1.30 | .195 |
| GroupFLUL | 0.03 | -0.18; 0.24 | 0.31 | .758 |
| Day2 x GroupStress | -0.02 | -0.42; 0.38 | -0.11 | .914 |
| Day2 x GroupFlu | -0.26 | -0.62; 0.10 | -1.43 | .156 |
| Day2 x GroupEPA | -0.13 | -0.48; 0.22 | -0.75 | .455 |
| **Day2 x GroupLPS** | **-0.69** | **-1.00; -0.39** | **-4.47** | **< .001** |
| Day2 x GroupFLUE | -0.13 | -0.47; 0.20 | -0.78 | .437 |
| **Day2 x GroupFLUL** | **-0.37** | **-0.70; -0.03** | **-2.14** | **.035** |
| **Freezing latency, s** | **Gauss distribution identity link** | | | |
| **predictor** | **estimate** | **ci** | **statistic** | **p.value** |
| **Intercept** | **293.26** | **263.38; 323.14** | **19.24** | **< .001** |
| Day2 | 7.74 | -42.77; 58.25 | 0.30 | .764 |
| GroupStress | 0.09 | -40.13; 40.32 | 0.00 | .996 |
| GroupFlu | -13.71 | -55.97; 28.54 | -0.64 | .526 |
| GroupEPA | 7.74 | -37.58; 53.06 | 0.33 | .738 |
| GroupLPS | -8.22 | -47.91; 31.48 | -0.41 | .686 |
| GroupFLUE | 7.74 | -35.39; 50.87 | 0.35 | .726 |
| GroupFLUL | -21.78 | -68.49; 24.94 | -0.91 | .363 |
| Day2 x GroupStress | -0.09 | -84.62; 84.44 | 0.00 | .998 |
| Day2 x GroupFlu | -57.20 | -136.86; 22.46 | -1.41 | .162 |
| Day2 x GroupEPA | -7.74 | -81.02; 65.54 | -0.21 | .836 |
| **Day2 x GroupLPS** | **-195.47** | **-273.80; -117.14** | **-4.89** | **< .001** |
| Day2 x GroupFLUE | -7.74 | -78.23; 62.75 | -0.22 | .830 |
| Day2 x GroupFLUL | -36.10 | -112.10; 39.89 | -0.93 | .354 |
| **Grooming test** | | | | |
| **Total grooming duration, s** | **Gamma distribution log link** | | | |
| **predictor** | **estimate** | **ci** | **statistic** | **p.value** |
| **Intercept** | **3.87** | **3.43; 4.39** | **15.81** | **< .001** |
| **Day2** | **-0.99** | **-1.78; -0.13** | **-2.39** | **.019** |
| **GroupStress** | **0.73** | **0.07; 1.38** | **2.22** | **.029** |
| GroupFlu | 0.02 | -0.66; 0.71 | 0.06 | .952 |
| GroupEPA | 0.26 | -0.46; 1.01 | 0.71 | .482 |
| GroupLPS | 0.07 | -0.59; 0.70 | 0.20 | .842 |
| GroupFLUE | 0.36 | -0.34; 1.06 | 1.02 | .309 |
| GroupFLUL | 0.57 | -0.17; 1.35 | 1.49 | .139 |
| Day2 x GroupStress | 0.10 | -1.14; 1.43 | 0.16 | .875 |
| **Day2 x GroupFlu** | **1.63** | **0.38; 2.97** | **2.50** | **.014** |
| Day2 x GroupEPA | 0.28 | -0.94; 1.50 | 0.45 | .650 |
| Day2 x GroupLPS | 0.25 | -0.98; 1.57 | 0.39 | .694 |
| Day2 x GroupFLUE | 0.89 | -0.26; 2.02 | 1.54 | .127 |
| Day2 x GroupFLUL | -0.03 | -1.22; 1.14 | -0.05 | .959 |
| **Rostral grooming duration, s** | **Gamma distribution log link** | | | |
| **predictor** | **estimate** | **ci** | **statistic** | **p.value** |
| **Intercept** | **3.77** | **3.28; 4.35** | **13.88** | **< .001** |
| **Day2** | **-0.92** | **-1.80; 0.03** | **-2.02** | **.046** |
| GroupStress | 0.62 | -0.11; 1.33 | 1.70 | .093 |
| GroupFlu | -0.19 | -0.95; 0.57 | -0.49 | .627 |
| GroupEPA | 0.10 | -0.70; 0.93 | 0.25 | .804 |
| GroupLPS | -0.15 | -0.87; 0.56 | -0.41 | .684 |
| GroupFLUE | 0.27 | -0.50; 1.05 | 0.69 | .495 |
| GroupFLUL | 0.51 | -0.31; 1.37 | 1.20 | .232 |
| Day2 x GroupStress | -0.02 | -1.39;1.46 | -0.03 | .979 |
| **Day2 x GroupFlu** | **1.66** | **0.27; 3.15** | **2.30** | **.023** |
| Day2 x GroupEPA | 0.43 | -0.92; 1.78 | 0.63 | .533 |
| Day2 x GroupLPS | 0.45 | -0.92; 1.92 | 0.63 | .532 |
| Day2 x GroupFLUE | 0.88 | -0.40; 2.14 | 1.37 | .172 |
| Day2 x GroupFLUL | -0.11 | -1.44; 1.18 | -0.17 | .864 |
| **Caudal grooming duration, s** | **Gamma distribution log link** | | | |
| **predictor** | **estimate** | **ci** | **statistic** | **p.value** |
| **Intercept** | **1.76** | **1.30; 2.30** | **6.90** | **< .001** |
| **Day2** | **-1.21** | **-2.03; -0.32** | **-2.81** | **.006** |
| **GroupStress** | **1.26** | **0.57; 1.93** | **3.67** | **< .001** |
| **GroupFlu** | **0.89** | **0.18; 1.61** | **2.48** | **.014** |
| **GroupEPA** | **0.99** | **0.23; 1.76** | **2.55** | **.012** |
| **GroupLPS** | **0.95** | **0.27; 1.61** | **2.80** | **.006** |
| **GroupFLUE** | **0.84** | **0.11; 1.57** | **2.27** | **.025** |
| **GroupFLUL** | **0.89** | **0.11; 1.69** | **2.22** | **.028** |
| Day2 x GroupStress | 0.57 | -0.71; 1.96 | 0.86 | .394 |
| **Day2 x GroupFlu** | **1.53** | **0.23; 2.93** | **2.26** | **.026** |
| Day2 x GroupEPA | -0.56 | -1.82; 0.71 | -0.87 | .384 |
| Day2 x GroupLPS | -0.59 | -1.88; 0.78 | -0.89 | .375 |
| Day2 x GroupFLUE | 0.84 | -0.35; 2.02 | 1.40 | .163 |
| Day2 x GroupFLUL | 0.39 | -0.85; 1.61 | 0.64 | .526 |
| **Paw licking duration, s** | **Inversed Gamma distribution identity link** | | | |
| **predictor** | **estimate** | **ci** | **statistic** | **p.value** |
| **Intercept** | **0.03** | **0.02; 0.05** | **3.58** | **.001** |
| Day2 | 0.04 | -0.01; 0.11 | 1.35 | .181 |
| GroupStress | -0.01 | -0.04; 0.00 | -1.42 | .158 |
| GroupFlu | 0.01 | -0.02; 0.04 | 0.64 | .521 |
| GroupEPA | 0.00 | -0.02; 0.03 | 0.25 | .807 |
| GroupLPS | 0.01 | -0.02; 0.03 | 0.50 | .616 |
| GroupFLUE | -0.01 | -0.03; 0.01 | -0.68 | .496 |
| GroupFLUL | -0.01 | -0.04; 0.01 | -1.18 | .242 |
| Day2 x GroupStress | -0.01 | -0.08; 0.08 | -0.13 | .893 |
| Day2 x GroupFlu | -0.06 | -0.13; 0.00 | -1.82 | .071 |
| Day2 x GroupEPA | -0.02 | -0.10; 0.05 | -0.61 | .542 |
| Day2 x GroupLPS | 0.01 | -0.09; 0.14 | 0.21 | .838 |
| Day2 x GroupFLUE | -0.04 | -0.11; 0.01 | -1.30 | .197 |
| Day2 x GroupFLUL | 0.00 | -0.08; 0.07 | 0.00 | .999 |
| **Nose grooming duration, s** | **Inversed Gamma distribution identity link** | | | |
| **predictor** | **estimate** | **ci** | **statistic** | **p.value** |
| **Intercept** | **0.08** | **0.04; 0.12** | **3.69** | **< .001** |
| Day2 | 0.18 | 0.03; 0.41 | 1.86 | .066 |
| GroupStress | -0.02 | -0.08; 0.02 | -0.95 | .344 |
| GroupFlu | 0.01 | -0.05; 0.07 | 0.21 | .834 |
| GroupEPA | -0.02 | -0.07; 0.04 | -0.61 | .544 |
| GroupLPS | 0.01 | -0.05; 0.07 | 0.33 | .743 |
| GroupFLUE | -0.01 | -0.06; 0.05 | -0.24 | .811 |
| GroupFLUL | -0.04 | -0.09; 0.01 | -1.46 | .146 |
| Day2 x GroupStress | -0.14 | -0.39; 0.04 | -1.36 | .177 |
| **Day2 x GroupFlu** | **-0.22** | **-0.46; -0.06** | **-2.20** | **.030** |
| Day2 x GroupEPA | -0.14 | -0.39; 0.04 | -1.33 | .186 |
| Day2 x GroupLPS | -0.18 | -0.43; 0.01 | -1.71 | .090 |
| Day2 x GroupFLUE | -0.15 | -0.40; 0.02 | -1.49 | .139 |
| Day2 x GroupFLUL | -0.10 | -0.35; 0.07 | -1.00 | .322 |
| **Head grooming duration, s** | **Inversed Gauss distribution identity link** | | | |
| **predictor** | **estimate** | **ci** | **statistic** | **p.value** |
| **Intercept** | **0.89** | **0.65; 1.13** | **7.27** | **< .001** |
| Day2 | 0.11 | -0.32; 0.53 | 0.49 | .622 |
| **GroupStress** | **-0.76** | **-1.02; -0.51** | **-5.89** | **< .001** |
| **GroupFlu** | **-0.34** | **-0.65; -0.03** | **-2.17** | **.032** |
| **GroupEPA** | **-0.39** | **-0.70; -0.08** | **-2.47** | **.015** |
| **GroupLPS** | **-0.37** | **-0.66; -0.08** | **-2.51** | **.014** |
| **GroupFLUE** | **-0.58** | **-0.86; -0.30** | **-4.01** | **< .001** |
| **GroupFLUL** | **-0.44** | **-0.77; -0.12** | **-2.65** | **.009** |
| Day2 x GroupStress | 0.24 | -0.30; 0.77 | 0.87 | .387 |
| Day2 x GroupFlu | -0.41 | -0.92; 0.11 | -1.54 | .127 |
| Day2 x GroupEPA | 0.12 | -0.44; 0.69 | 0.43 | .669 |
| Day2 x GroupLPS | 0.00 | -0.58; 0.58 | -0.01 | .995 |
| Day2 x GroupFLUE | 0.04 | -0.46; 0.54 | 0.15 | .883 |
| Day2 x GroupFLUL | 0.38 | -0.18; 0.94 | 1.33 | .185 |
| **Body grooming duration, s** | **Inversed Gauss distribution log link** | | | |
| **predictor** | **estimate** | **ci** | **statistic** | **p.value** |
| **Intercept** | **0.89** | **0.65; 1.13** | **7.27** | **< .001** |
| Day2 | 0.11 | -0.32; 0.53 | 0.49 | .622 |
| **GroupStress** | **-0.76** | **-1.02; -0.51** | **-5.89** | **< .001** |
| **GroupFlu** | **-0.34** | **-0.65; -0.03** | **-2.17** | **.032** |
| **GroupEPA** | **-0.39** | **-0.70; -0.08** | **-2.47** | **.015** |
| **GroupLPS** | **-0.37** | **-0.66; -0.08** | **-2.51** | **.014** |
| **GroupFLUE** | **-0.58** | **-0.86; -0.30** | **-4.01** | **< .001** |
| **GroupFLUL** | **-0.44** | **-0.77; -0.12** | **-2.65** | **.009** |
| Day2 x GroupStress | 0.24 | -0.30; 0.77 | 0.87 | .387 |
| Day2 x GroupFlu | -0.41 | -0.92; 0.11 | -1.54 | .127 |
| Day2 x GroupEPA | 0.12 | -0.44; 0.69 | 0.43 | .669 |
| Day2 x GroupLPS | 0.00 | -0.58; 0.58 | -0.01 | .995 |
| Day2 x GroupFLUE | 0.04 | -0.46; 0.54 | 0.15 | .883 |
| Day2 x GroupFLUL | 0.38 | -0.18; 0.94 | 1.33 | .185 |
| **Tail grooming duration, s** | **Inversed Gauss distribution log link** | | | |
| **predictor** | **estimate** | **ci** | **statistic** | **p.value** |
| **Intercept** | **0.89** | **0.65; 1.13** | **7.27** | **< .001** |
| Day2 | 0.11 | -0.32; 0.53 | 0.49 | .622 |
| **GroupStress** | **-0.76** | **-1.02; -0.51** | **-5.89** | **< .001** |
| **GroupFlu** | **-0.34** | **-0.65; -0.03** | **-2.17** | **.032** |
| **GroupEPA** | **-0.39** | **-0.70; -0.08** | **-2.47** | **.015** |
| **GroupLPS** | **-0.37** | **-0.66; -0.08** | **-2.51** | **.014** |
| **GroupFLUE** | **-0.58** | **-0.86; -0.30** | **-4.01** | **< .001** |
| **GroupFLUL** | **-0.44** | **-0.77; -0.12** | **-2.65** | **.009** |
| Day2 x GroupStress | 0.24 | -0.30; 0.77 | 0.87 | .387 |
| Day2 x GroupFlu | -0.41 | -0.92; 0.11 | -1.54 | .127 |
| Day2 x GroupEPA | 0.12 | -0.44; 0.69 | 0.43 | .669 |
| Day2 x GroupLPS | 0.00 | -0.58; 0.58 | -0.01 | .995 |
| Day2 x GroupFLUE | 0.04 | -0.46; 0.54 | 0.15 | .883 |
| Day2 x GroupFLUL | 0.38 | -0.18; 0.94 | 1.33 | .185 |
| **Incorrect grooming trasitions, %** | **Inversed Gauss distribution log link** | | | |
| **predictor** | **estimate** | **ci** | **statistic** | **p.value** |
| **Intercept** | **-0.62** | **-0.70; -0.53** | **-13.98** | **< .001** |
| Day2 | 0.06 | -0.15; 0.31 | 0.56 | .578 |
| GroupStress | -0.10 | -0.21; 0.01 | -1.71 | .090 |
| GroupFlu | 0.06 | -0.06; 0.19 | 1.01 | .313 |
| GroupEPA | -0.04 | -0.17; 0.09 | -0.65 | .519 |
| GroupLPS | 0.03 | -0.09; 0.14 | 0.43 | .666 |
| GroupFLUE | -0.07 | -0.19; 0.05 | -1.19 | .237 |
| GroupFLUL | -0.08 | -0.21; 0.04 | -1.31 | .194 |
| Day2 x GroupStress | 0.13 | -0.16; 0.40 | 0.90 | .370 |
| Day2 x GroupFlu | -0.16 | -0.45; 0.12 | -1.11 | .271 |
| Day2 x GroupEPA | -0.06 | -0.34; 0.21 | -0.40 | .687 |
| Day2 x GroupLPS | -0.16 | -0.45; 0.11 | -1.10 | .275 |
| Day2 x GroupFLUE | 0.02 | -0.26; 0.27 | 0.16 | .873 |
| Day2 x GroupFLUL | 0.12 | -0.17; 0.38 | 0.86 | .389 |

**Supplementary Table S2.** Summary of post-hoc Tukey’s test results for significant Wald Chi-square test (ANOVA Type II; Table 3 in the MS) for generalized linear model (GZLM; Supplementary Table S1) group, testing day and their interaction effects as ‘predictors’, to compare behavior in experimental groups. CUS – chronic unpredictable stress, FLU – fluoxetine, EPA - eicosapentaenoic acid, LPS – lipopolysaccharide, FLUE – fluoxetine and eicosapentaenoic acid, FLUL – fluoxetine and lipopolysaccharide.

| **Comparison** | **estimate** | **SE** | **z.ratio** | **p.value** |
| --- | --- | --- | --- | --- |
| **Open Field Test** | | | | |
| **Freezing frequency, n** | | | | |
| **Control - Stress** | **0.46969696969697** | **0.156286914002489** | **3.00535059313724** | **0.00265274885909498** |
| Control - Flu | 0.175 | 0.183670288902804 | 0.952794276338331 | 0.34069431487817 |
| Control - EPA | 0.297202797202797 | 0.158609011677558 | 1.8738077619889 | 0.0609569261817316 |
| Control - LPS | 0.09375 | 0.185865677322188 | 0.504396515541109 | 0.61398276615223 |
| Control - FLUE | 0.26 | 0.1580284950034 | 1.64527289837448 | 0.0999135461635716 |
| **Control - FLUL** | **0.510489510489511** | **0.149019219384859** | **3.4256622239519** | **0.000613302565932391** |
| Stress - Flu | -0.29469696969697 | 0.17213954670849 | -1.71196552641111 | 0.0869030216670451 |
| Stress - EPA | -0.172494172494173 | 0.145100196763121 | -1.18879351194659 | 0.234520931771147 |
| **Stress - LPS** | **-0.37594696969697** | **0.174480080587443** | **-2.15466985360864** | **0.0311876784663105** |
| Stress - FLUE | -0.209696969696969 | 0.144465406753327 | -1.45153759927471 | 0.146630220340125 |
| Stress - FLUL | 0.0407925407925414 | 0.134551017318478 | 0.303175268426153 | 0.761756295529099 |
| **Freezing duration, s** | | | | |
| **Control - Stress** | **0.710176775881035** | **0.207449422216666** | **3.423373120506** | **0.000618491176029704** |
| Control - Flu | 0.295017344686221 | 0.275531116522368 | 1.07072242296913 | 0.284294257454084 |
| **Control - EPA** | **0.787431989319194** | **0.201727594230937** | **3.90344212610668** | **9.48342348944871e-05** |
| Control - LPS | 0.361495844875345 | 0.272706047185205 | 1.32558793105838 | 0.184976226033893 |
| **Control - FLUE** | **0.470604086032041** | **0.230117673124929** | **2.0450584244199** | **0.0408490984916596** |
| **Control - FLUL** | **0.831987664015749** | **0.199106682281922** | **4.1786024179626** | **2.93305909085056e-05** |
| Stress - Flu | -0.415159431194814 | 0.228882705001823 | -1.81385234498827 | 0.0697004611639784 |
| Stress - EPA | 0.0772552134381592 | 0.131011139797691 | 0.589684308963784 | 0.555402316819736 |
| Stress - LPS | -0.34868093100569 | 0.225473911236462 | -1.54643581199076 | 0.121999350841795 |
| Stress - FLUE | -0.239572689848993 | 0.171534369621136 | -1.39664540918611 | 0.162520227930602 |
| Stress - FLUL | 0.121810888134714 | 0.126938439435038 | 0.959606000175007 | 0.337253548087636 |
| **Rearing frequency, n** | | | | |
| **Control - Stress** | **0.620856566154394** | **0.278949384294071** | **2.22569613381861** | **0.0260345419080056** |
| Control - Flu | -0.289909247626469 | 0.417033861813852 | -0.695169563367192 | 0.486949044215772 |
| Control - EPA | 0.388412301864911 | 0.277107526900254 | 1.40166637193049 | 0.161014897021338 |
| Control - LPS | 0.228379201243616 | 0.291397802818517 | 0.783736867727351 | 0.433194524819842 |
| Control - FLUE | -0.0898362757875543 | 0.292856642486416 | -0.306758539006747 | 0.759027168716148 |
| Control - FLUL | -0.151593131440744 | 0.312736940872269 | -0.484730492719947 | 0.627867533448341 |
| **Stress - Flu** | **-0.910765813780864** | **0.41056388375785** | **-2.2183291073845** | **0.0265324009271837** |
| Stress - EPA | -0.232444264289483 | 0.267271476614564 | -0.869693493798046 | 0.384467928765283 |
| Stress - LPS | -0.392477364910778 | 0.282060525835171 | -1.39146505434841 | 0.164084449441379 |
| **Stress - FLUE** | **-0.710692841941949** | **0.283567406087814** | **-2.50625716032351** | **0.0122016840547677** |
| **Stress - FLUL** | **-0.772449697595138** | **0.304055677357124** | **-2.54048766432955** | **0.0110697998576173** |
| **Rearing duration, s** | | | | |
| **Control - Stress** | **0.777476923376969** | **0.326168111552375** | **2.38366932830012** | **0.0171409967251114** |
| Control - Flu | -0.171669608392832 | 0.513583511406314 | -0.334258410911129 | 0.738184570578572 |
| Control - EPA | 0.489904630148168 | 0.338803719973083 | 1.44598362198352 | 0.148181795592086 |
| Control - LPS | 0.173977881346444 | 0.363556421631481 | 0.47854437714429 | 0.632262796891433 |
| Control - FLUE | -0.0297777981588389 | 0.352830926121629 | -0.0843967916479459 | 0.932740958081241 |
| Control - FLUL | -0.155283352285916 | 0.389196616578106 | -0.398984332523746 | 0.689904748639346 |
| Stress - Flu | -0.949146531769801 | 0.495154092831778 | -1.91687102158774 | 0.055254319378093 |
| Stress - EPA | -0.287572293228801 | 0.310157239359685 | -0.927182269943109 | 0.353831906752411 |
| Stress - LPS | -0.603499042030525 | 0.337020510012535 | -1.79068936192661 | 0.0733431577792494 |
| **Stress - FLUE** | **-0.807254721535808** | **0.325421595607095** | **-2.480642749077** | **0.0131145740065355** |
| **Stress - FLUL** | **-0.932760275662885** | **0.364531972281536** | **-2.55878865665724** | **0.0105037576789421** |
| **Elevated Plus-Maze Test** | | | | |
| **Freezing frequency, n** | | | | |
| Control - Stress | -0.0111111111111098 | 0.102204978035183 | -0.10871399147784 | 0.913429344038785 |
| Control - Flu | 0.0690476190476201 | 0.0916268356488814 | 0.753574196452817 | 0.451104943313793 |
| Control - EPA | -0.0666666666666654 | 0.0888635797301192 | -0.750213606846963 | 0.453126065024592 |
| **Control - LPS** | **0.319298245614037** | **0.0776584264144851** | **4.11157243786851** | **3.92973538797608e-05** |
| Control - FLUE | -0.0666666666666654 | 0.0854054458182971 | -0.780590347932859 | 0.435043470450822 |
| Control - FLUL | 0.150000000000001 | 0.0856888493564572 | 1.75051947979854 | 0.0800287158156968 |
| Stress - Flu | 0.0801587301587299 | 0.10736136869045 | 0.746625449511993 | 0.455289681919891 |
| Stress - EPA | -0.0555555555555556 | 0.105012962430887 | -0.529035218791385 | 0.596781018741469 |
| **Stress - LPS** | **0.330409356725146** | **0.095717384362683** | **3.45192630288759** | **0.000556599729966254** |
| Stress - FLUE | -0.0555555555555557 | 0.102103264646541 | -0.544111451753052 | 0.586364780485787 |
| Stress - FLUL | 0.161111111111111 | 0.102340438635435 | 1.57426637269979 | 0.115425880502261 |
| **Freezing duration, s** | | | | |
| Control - Stress | 0.0941891115207032 | 0.304470338390748 | 0.309353981798463 | 0.757052271808696 |
| Control - Flu | 0.238394391703598 | 0.237650843842589 | 1.0031287406726 | 0.315798749220113 |
| Control - EPA | -0.231404958677683 | 0.270929865033374 | -0.854113881646741 | 0.393041893109589 |
| **Control - LPS** | **0.418527715445802** | **0.201375644334903** | **2.07834327149195** | **0.0376777536306161** |
| Control - FLUE | -0.231404958677683 | 0.25974741726456 | -0.890884541277232 | 0.372991114353059 |
| **Control - FLUL** | **0.466001647047475** | **0.211165392001667** | **2.20680880815828** | **0.0273274194396686** |
| Stress - Flu | 0.144205280182895 | 0.293024198578782 | 0.492127547425488 | 0.622629179244374 |
| Stress - EPA | -0.325594070198386 | 0.32060606535224 | -1.01555804891173 | 0.309839890485129 |
| Stress - LPS | 0.324338603925098 | 0.264456815954779 | 1.22643314279545 | 0.220035720279261 |
| Stress - FLUE | -0.325594070198386 | 0.311213717803782 | -1.04620732175974 | 0.295465323212462 |
| Stress - FLUL | 0.371812535526772 | 0.271985441067827 | 1.36703102220111 | 0.171615582006025 |
| **Freezing latency, s** | | | | |
| Control - Stress | -0.0472596153841846 | 21.564251466134 | -0.00219157226293732 | 0.998251379727286 |
| **Control - Flu** | **42.3131730769237** | **20.3215180687904** | **2.08218563857726** | **0.0373255165487241** |
| Control - EPA | -3.87038461538413 | 18.6939317634915 | -0.207039624641341 | 0.835978912886307 |
| **Control - LPS** | **105.953438914027** | **19.982297516162** | **5.30236519741189** | **1.14311872372783e-07** |
| Control - FLUE | -3.8703846153842 | 17.9829025056913 | -0.21522580207279 | 0.829591275134765 |
| **Control - FLUL** | **39.8279487179491** | **19.3866226462352** | **2.05440367023823** | **0.0399366381366158** |
| Stress - Flu | 42.3604326923079 | 23.3655644795146 | 1.81294283429132 | 0.0698406387480441 |
| Stress - EPA | -3.82312499999995 | 21.964712417973 | -0.174057594165067 | 0.861820199367455 |
| **Stress - LPS** | **106.000698529412** | **23.0711447667989** | **4.59451403911064** | **4.33758926103964e-06** |
| Stress - FLUE | -3.82312500000002 | 21.3628249385557 | -0.178961584481274 | 0.857967865812222 |
| Stress - FLUL | 39.8752083333333 | 22.5571860934315 | 1.76773858974124 | 0.0771046154896095 |
| **Closed Arm frequency, n** | | | | |
| **Control - Stress** | **0.437789866547868** | **0.159655377573022** | **2.74209283271799** | **0.00610490852004394** |
| Control - Flu | 0.182377054658061 | 0.158407830142142 | 1.15131338201155 | 0.249603333761835 |
| Control - EPA | 0.1156113583526 | 0.151318293476413 | 0.764027638010744 | 0.444850772573025 |
| Control - LPS | 0.0145230533002927 | 0.158009173405007 | 0.0919127224535719 | 0.926767383195223 |
| Control - FLUE | -0.0526247599674835 | 0.151632247921114 | -0.347055198936716 | 0.728549850098935 |
| **Control - FLUL** | **-0.39263185043689** | **0.1811878391357** | **-2.16698787462678** | **0.0302357786949988** |
| Stress - Flu | -0.255412811889807 | 0.164362983005772 | -1.55395580695221 | 0.120194958472643 |
| **Stress - EPA** | **-0.322178508195267** | **0.157541662657304** | **-2.0450368668261** | **0.0408512236086299** |
| **Stress - LPS** | **-0.423266813247575** | **0.163978804765577** | **-2.58122879876258** | **0.00984493099292707** |
| **Stress - FLUE** | **-0.490414626515351** | **0.157843239138786** | **-3.10697264698267** | **0.00189013886846301** |
| **Stress - FLUL** | **-0.830421716984758** | **0.186416690730413** | **-4.45465324875698** | **8.40288906928029e-06** |
| **Open Arm duration, s** | | | | |
| **Control - Stress** | **1.74607531385394** | **0.380417352622105** | **4.58989397255081** | **4.43471217859724e-06** |
| Control - Flu | -0.116740169278498 | 0.573803275170163 | -0.203449813429311 | 0.838783468358895 |
| Control - EPA | -0.00371515132930269 | 0.495613670544903 | -0.00749606306302662 | 0.994019063027692 |
| **Control - LPS** | **1.2401457558395** | **0.415646578067002** | **2.98365443451235** | **0.00284828256030728** |
| Control - FLUE | 0.13348570587857 | 0.469212615203796 | 0.284488740398833 | 0.776035861590977 |
| Control - FLUL | -0.0461303001777452 | 0.547301729471735 | -0.0842867794740408 | 0.932828423448302 |
| **Stress - Flu** | **-1.86281548313243** | **0.492004531988685** | **-3.78617545574819** | **0.000152983674880332** |
| **Stress - EPA** | **-1.74979046518324** | **0.398059256056671** | **-4.39580398787191** | **1.1036349223484e-05** |
| Stress - LPS | -0.505929558014439 | 0.292541174469864 | -1.72943025518125 | 0.0837321190037845 |
| **Stress - FLUE** | **-1.61258960797536** | **0.364662500358597** | **-4.42214268368587** | **9.77268820239559e-06** |
| **Stress - FLUL** | **-1.79220561403168** | **0.460822573214627** | **-3.88914458232706** | **0.000100598176327873** |
| **Grooming Test** | | | | |
| **Caudal grooming duration, s** | | | | |
| **Control - Stress** | **-1.54613106926408** | **0.335059972491588** | **-4.61449052767083** | **3.94060745675673e-06** |
| **Control - Flu** | **-1.66163036669445** | **0.339569152807356** | **-4.89334897754132** | **9.9134392512862e-07** |
| **Control - EPA** | **-0.70535105076236** | **0.320309705935513** | **-2.20209078180218** | **0.0276588965054381** |
| Control - LPS | -0.649359908988624 | 0.333900834363775 | -1.94476875215342 | 0.0518028165631409 |
| **Control - FLUE** | **-1.25747074733622** | **0.300491279647709** | **-4.18471627133557** | **2.85522499442086e-05** |
| **Control - FLUL** | **-1.08255225678664** | **0.310086637490422** | **-3.49112836834213** | **0.000480985094942897** |
| Stress - Flu | -0.115499297430369 | 0.367237090314576 | -0.314508802287458 | 0.753134632146715 |
| **Stress - EPA** | **0.84078001850172** | **0.34950562038529** | **2.40562660358611** | **0.0161447567975816** |
| **Stress - LPS** | **0.896771160275456** | **0.362002262635521** | **2.47725291479287** | **0.0132398045328731** |
| Stress - FLUE | 0.288660321927864 | 0.331437596100275 | 0.870934152685958 | 0.383790107633956 |
| Stress - FLUL | 0.463578812477445 | 0.340161129046997 | 1.36282124232189 | 0.172938854687677 |
| **Head grooming duration, s** | | | | |
| **Control - Stress** | **0.645956197308847** | **0.136133885623832** | **4.74500668477035** | **2.08499449878462e-06** |
| **Control - Flu** | **0.542043105890946** | **0.132085406935613** | **4.10373195999747** | **4.06538446837759e-05** |
| **Control - EPA** | **0.329178747481603** | **0.144281825899368** | **2.28149834831723** | **0.0225189743361392** |
| **Control - LPS** | **0.371144692965931** | **0.148032757095094** | **2.50717949357326** | **0.0121698894361437** |
| **Control - FLUE** | **0.559848317572587** | **0.127271857114164** | **4.3988382841809** | **1.08831882945227e-05** |
| Control - FLUL | 0.249395202933738 | 0.143035964458085 | 1.74358388730144 | 0.0812316749347003 |
| Stress - Flu | -0.103913091417901 | 0.11295197431169 | -0.919975875155167 | 0.357585366351597 |
| **Stress - EPA** | **-0.316777449827244** | **0.126999366378018** | **-2.49432307311161** | **0.0126197622698591** |
| **Stress - LPS** | **-0.274811504342916** | **0.131245155905798** | **-2.09387921745594** | **0.0362707410391488** |
| Stress - FLUE | -0.0861078797362601 | 0.107283360260809 | -0.802621017154283 | 0.422193816814511 |
| **Stress - FLUL** | **-0.396560994375109** | **0.125582167938035** | **-3.15778108378239** | **0.00158974892557385** |
| **Body grooming duration, s** | | | | |
| **Control - Stress** | **-1.31633690562698** | **0.326561218798783** | **-4.03090394649116** | **5.55627486284399e-05** |
| **Control - Flu** | **-1.49607609016759** | **0.414250468404162** | **-3.61152540377564** | **0.000304401259039456** |
| **Control - EPA** | **-0.475227518270103** | **0.240518030348118** | **-1.975849866982** | **0.048171783618369** |
| **Control - LPS** | **-0.48622930087043** | **0.233009971731898** | **-2.08673172764419** | **0.0369123915846793** |
| **Control - FLUE** | **-1.20193074933191** | **0.281157746125938** | **-4.27493379034819** | **1.91194104287279e-05** |
| **Control - FLUL** | **-0.619233893180414** | **0.255084136303271** | **-2.42756724175197** | **0.0152004688023721** |
| Stress - Flu | -0.17973918454061 | 0.489268393830745 | -0.367363162646447 | 0.713348145490489 |
| **Stress - EPA** | **0.841109387356875** | **0.354441862018565** | **2.37305317878287** | **0.0176417237279929** |
| **Stress - LPS** | **0.830107604756549** | **0.349390551611158** | **2.37587307650033** | **0.0175074838270681** |
| Stress - FLUE | 0.114406156295065 | 0.383183753355054 | 0.298567346066622 | 0.765270183628958 |
| Stress - FLUL | 0.697103012446564 | 0.364483233115372 | 1.91257909585626 | 0.0558019579166375 |
| **Tail grooming duration, s** | | | | |
| **Control - Stress** | **-1.36612322395203** | **0.353089866729667** | **-3.86905247835381** | **0.00010925912611441** |
| **Control - Flu** | **-1.32288316053145** | **0.36883222776737** | **-3.58667996161609** | **0.000334914954425839** |
| **Control - EPA** | **-0.719525657530046** | **0.270504715633943** | **-2.65993757574169** | **0.00781551352961339** |
| **Control - LPS** | **-0.667693510025904** | **0.259121777462688** | **-2.57675567281121** | **0.00997324181833673** |
| **Control - FLUE** | **-0.812370873791946** | **0.257116490046534** | **-3.15954404030997** | **0.00158016216784136** |
| **Control - FLUL** | **-1.28656796882391** | **0.324265461320288** | **-3.96763800740752** | **7.25884785315598e-05** |
| Stress - Flu | 0.0432400634205868 | 0.468329136370533 | 0.0923283649522418 | 0.926437152642374 |
| Stress - EPA | 0.646597566421988 | 0.3955600699185 | 1.63463811338493 | 0.102124932917725 |
| Stress - LPS | 0.69842971392613 | 0.387864748706493 | 1.80070428224105 | 0.0717495023050367 |
| Stress - FLUE | 0.553752350160089 | 0.386527951366493 | 1.43263209866817 | 0.151963003582261 |
| Stress - FLUL | 0.0795552551281279 | 0.434100284656653 | 0.183264692376443 | 0.854590336583979 |

**Supplementary Table S3.** Summary of statistical analyses for all differentially expressed genes in experimental groups vs. control group or in experimental groups vs. stress group using the DESeq function. False discovery rate was set at 0.01(6) for control comparison and 0.02 for CUS comparison. Genes in this table are listed as sorted by their p-adjusted value. CUS – chronic unpredictable stress, FLU – fluoxetine, EPA - eicosapentaenoic acid, LPS – lipopolysaccharide.

| **Lookup** | **entrezgene_accession** | **baseMean** | **log2FoldChange** | **lfcSE** | **stat** | **pvalue** | **padj** |
| --- | --- | --- | --- | --- | --- | --- | --- |
| **CUS vs. Control** | | | | | | | |
| ENSRNOG00000015554 | *Ankdd1a* | 161.2166 | -2.86674 | 0.42303 | -6.77667 | 1.23E-11 | 1.87E-07 |
| ENSRNOG00000007302 | *Fbn1* | 780.2847 | -2.08037 | 0.345227 | -6.02609 | 1.68E-09 | 8.50E-06 |
| ENSRNOG00000010557 | *Smarcd2* | 1205.071 | -0.98074 | 0.161164 | -6.08534 | 1.16E-09 | 8.50E-06 |
| ENSRNOG00000001036 | *Rsph10b* | 880.4996 | -1.99174 | 0.33539 | -5.9386 | 2.87E-09 | 1.07E-05 |
| ENSRNOG00000005882 | *Tle1* | 2075.44 | -1.524 | 0.260537 | -5.84945 | 4.93E-09 | 1.07E-05 |
| ENSRNOG00000013036 | *Epha8* | 832.0532 | -3.09013 | 0.523979 | -5.89744 | 3.69E-09 | 1.07E-05 |
| ENSRNOG00000036829 | *Nckap1l* | 629.4801 | 1.207989 | 0.206261 | 5.856592 | 4.72E-09 | 1.07E-05 |
| ENSRNOG00000049959 | *Igsf21* | 563.1432 | 1.854891 | 0.321243 | 5.774097 | 7.74E-09 | 1.47E-05 |
| ENSRNOG00000033110 | *Svep1* | 494.32 | -2.5808 | 0.454067 | -5.68375 | 1.32E-08 | 2.22E-05 |
| ENSRNOG00000001982 | *Cblb* | 960.7187 | -1.17253 | 0.208769 | -5.61639 | 1.95E-08 | 2.96E-05 |
| ENSRNOG00000039759 | *Gpr34* | 1146.162 | 1.210424 | 0.216547 | 5.589664 | 2.28E-08 | 3.14E-05 |
| ENSRNOG00000021663 | *Vxn* | 14560.56 | 1.148891 | 0.208275 | 5.516228 | 3.46E-08 | 4.05E-05 |
| ENSRNOG00000026974 | *Dbndd1* | 800.9696 | 1.013785 | 0.183424 | 5.527007 | 3.26E-08 | 4.05E-05 |
| ENSRNOG00000005971 | *Gpr176* | 971.3604 | -1.22063 | 0.222851 | -5.47734 | 4.32E-08 | 4.52E-05 |
| ENSRNOG00000007300 | *C1qtnf6* | 97.94694 | -1.90073 | 0.347393 | -5.47142 | 4.46E-08 | 4.52E-05 |
| ENSRNOG00000003694 | *Prox1* | 2091.809 | -2.502 | 0.460396 | -5.43445 | 5.50E-08 | 5.22E-05 |
| ENSRNOG00000007030 | *Epha7* | 3329.05 | -1.82966 | 0.337809 | -5.41626 | 6.09E-08 | 5.44E-05 |
| ENSRNOG00000002215 | *Mylk* | 914.7898 | 0.785136 | 0.146 | 5.377662 | 7.55E-08 | 6.37E-05 |
| ENSRNOG00000002930 | *Ppl* | 528.9597 | -4.21558 | 0.79221 | -5.32129 | 1.03E-07 | 8.24E-05 |
| ENSRNOG00000001427 | *Orai2* | 4943.709 | -0.99669 | 0.190022 | -5.24513 | 1.56E-07 | 0.000119 |
| ENSRNOG00000011589 | *Camk2d* | 1173.796 | 2.120604 | 0.405619 | 5.22807 | 1.71E-07 | 0.000124 |
| ENSRNOG00000014613 | *Ddah1* | 2529.238 | 1.020939 | 0.195642 | 5.218418 | 1.80E-07 | 0.000125 |
| ENSRNOG00000038980 | *Lypd6* | 861.4353 | 1.34777 | 0.259081 | 5.202125 | 1.97E-07 | 0.00013 |
| ENSRNOG00000004152 | *Lrp12* | 2418.609 | -1.13192 | 0.218421 | -5.1823 | 2.19E-07 | 0.000139 |
| ENSRNOG00000004249 | *Tlr7* | 290.8621 | 1.157472 | 0.224126 | 5.164382 | 2.41E-07 | 0.000147 |
| ENSRNOG00000012480 | *Pxylp1* | 910.2834 | -1.1306 | 0.219588 | -5.14872 | 2.62E-07 | 0.000153 |
| ENSRNOG00000018808 | *Vip* | 666.4469 | 1.919861 | 0.375722 | 5.109793 | 3.23E-07 | 0.000181 |
| ENSRNOG00000016366 | *Colec12* | 293.0481 | 2.260612 | 0.443343 | 5.099011 | 3.41E-07 | 0.000185 |
| ENSRNOG00000001757 | *Tm4sf19* | 155.3925 | 2.553519 | 0.502567 | 5.080955 | 3.76E-07 | 0.000197 |
| ENSRNOG00000005565 | *Traf3ip3* | 210.905 | 1.402681 | 0.277533 | 5.054099 | 4.32E-07 | 0.000199 |
| ENSRNOG00000005809 | *Arhgdib* | 525.6785 | 1.073443 | 0.212067 | 5.061818 | 4.15E-07 | 0.000199 |
| ENSRNOG00000011977 | *Sema5a* | 5717.816 | -1.8062 | 0.357139 | -5.05741 | 4.25E-07 | 0.000199 |
| ENSRNOG00000046452 | *Fcgr2b* | 372.4628 | 1.978598 | 0.390843 | 5.06238 | 4.14E-07 | 0.000199 |
| ENSRNOG00000031443 | *Havcr2* | 285.2417 | 1.208469 | 0.240001 | 5.035268 | 4.77E-07 | 0.000213 |
| ENSRNOG00000007027 | *Hgf* | 282.4444 | 1.834304 | 0.366291 | 5.007775 | 5.51E-07 | 0.000232 |
| ENSRNOG00000016573 | *Dgat2* | 4260.065 | -1.41576 | 0.283005 | -5.00259 | 5.66E-07 | 0.000232 |
| ENSRNOG00000054695 | *Calcrl* | 739.4976 | 1.057321 | 0.210956 | 5.012039 | 5.39E-07 | 0.000232 |
| ENSRNOG00000026679 | *Scn4b* | 1530.313 | -2.72345 | 0.547563 | -4.97377 | 6.57E-07 | 0.000262 |
| ENSRNOG00000001528 | *Gap43* | 11009.53 | 1.310238 | 0.264086 | 4.961413 | 7.00E-07 | 0.000272 |
| ENSRNOG00000017659 | *Hs3st2* | 259.2726 | 2.006245 | 0.404758 | 4.956655 | 7.17E-07 | 0.000272 |
| ENSRNOG00000061639 | | 46.05165 | -3.33835 | 0.67524 | -4.94394 | 7.66E-07 | 0.000284 |
| ENSRNOG00000034102 | *Plk5* | 973.7102 | -2.47042 | 0.501796 | -4.92316 | 8.52E-07 | 0.000308 |
| ENSRNOG00000001480 | *Ncf1* | 260.4873 | 0.93155 | 0.190824 | 4.88173 | 1.05E-06 | 0.000343 |
| ENSRNOG00000004554 | *Dcn* | 566.9096 | 1.326128 | 0.271845 | 4.878244 | 1.07E-06 | 0.000343 |
| ENSRNOG00000019009 | *Arrdc2* | 1058.097 | -1.2963 | 0.266077 | -4.87189 | 1.11E-06 | 0.000343 |
| ENSRNOG00000029865 | *Prss56* | 25.78381 | -5.13175 | 1.051769 | -4.87916 | 1.07E-06 | 0.000343 |
| ENSRNOG00000031930 | *Bin2* | 702.7842 | 0.950416 | 0.194312 | 4.891173 | 1.00E-06 | 0.000343 |
| ENSRNOG00000033796 | *Kcnj5* | 154.6946 | 1.456311 | 0.298696 | 4.875565 | 1.08E-06 | 0.000343 |
| ENSRNOG00000036699 | *Faap100* | 608.6026 | -1.366 | 0.278914 | -4.89757 | 9.70E-07 | 0.000343 |
| ENSRNOG00000058568 | *Dhrs9* | 63.20553 | 2.800542 | 0.576102 | 4.861194 | 1.17E-06 | 0.000354 |
| ENSRNOG00000006110 | *Jph1* | 1252.077 | -1.72976 | 0.357881 | -4.83335 | 1.34E-06 | 0.0004 |
| ENSRNOG00000008182 | *Htra3* | 463.5113 | 1.197522 | 0.248376 | 4.821405 | 1.43E-06 | 0.000416 |
| ENSRNOG00000003237 | *Tp53bp2* | 5045.185 | 1.228524 | 0.255523 | 4.807889 | 1.53E-06 | 0.000437 |
| ENSRNOG00000029165 | *Stx1a* | 4386.499 | 1.81807 | 0.378752 | 4.800155 | 1.59E-06 | 0.000446 |
| ENSRNOG00000030016 | *Robo3* | 746.3388 | -3.52895 | 0.73767 | -4.78392 | 1.72E-06 | 0.000475 |
| ENSRNOG00000006674 | *Rflnb* | 178.3245 | -1.18001 | 0.247108 | -4.77529 | 1.79E-06 | 0.000487 |
| ENSRNOG00000001302 | *Adora2a* | 368.615 | -3.99869 | 0.840427 | -4.75793 | 1.96E-06 | 0.000512 |
| ENSRNOG00000025848 | *Sspo* | 39.89744 | -4.60758 | 0.968341 | -4.75823 | 1.95E-06 | 0.000512 |
| ENSRNOG00000001928 | *Il1rap* | 1298.06 | -1.20923 | 0.254707 | -4.74751 | 2.06E-06 | 0.00053 |
| ENSRNOG00000013304 | *Arg1* | 662.2912 | -2.15276 | 0.454832 | -4.73309 | 2.21E-06 | 0.000551 |
| ENSRNOG00000039050 | | 169.035 | -1.2649 | 0.267188 | -4.73411 | 2.20E-06 | 0.000551 |
| ENSRNOG00000043366 | | 342.4103 | -1.62061 | 0.342875 | -4.72652 | 2.28E-06 | 0.00056 |
| ENSRNOG00000002191 | *LOC498368* | 525.2488 | 1.000657 | 0.213095 | 4.695833 | 2.66E-06 | 0.00064 |
| ENSRNOG00000010535 | *Cdh18* | 707.2539 | 1.096551 | 0.233933 | 4.687448 | 2.77E-06 | 0.000646 |
| ENSRNOG00000024312 | *Xkr8* | 397.1491 | -1.3072 | 0.27878 | -4.68901 | 2.75E-06 | 0.000646 |
| ENSRNOG00000054058 | *Osbpl1a* | 5099.777 | 0.993734 | 0.212583 | 4.674576 | 2.95E-06 | 0.000678 |
| ENSRNOG00000010266 | *Cd180* | 258.2559 | 0.987007 | 0.2119 | 4.657897 | 3.19E-06 | 0.000724 |
| ENSRNOG00000036834 | *Gpr84* | 279.5048 | 1.288155 | 0.278105 | 4.631902 | 3.62E-06 | 0.000809 |
| ENSRNOG00000019240 | *Ampd2* | 3536.156 | -0.81848 | 0.177078 | -4.62213 | 3.80E-06 | 0.000836 |
| ENSRNOG00000027456 | *Cdc42bpg* | 579.6898 | -1.56732 | 0.339335 | -4.61881 | 3.86E-06 | 0.000837 |
| ENSRNOG00000006850 | *Ovol2* | 23.81928 | 5.520269 | 1.197512 | 4.609782 | 4.03E-06 | 0.000862 |
| ENSRNOG00000013282 | *Mctp1* | 1395.673 | -1.34115 | 0.29141 | -4.60227 | 4.18E-06 | 0.000882 |
| ENSRNOG00000018851 | *Ppp2r2b* | 5883.072 | 0.714282 | 0.155515 | 4.59301 | 4.37E-06 | 0.000909 |
| ENSRNOG00000006885 | *Dbx2* | 164.5865 | 1.577322 | 0.34419 | 4.582712 | 4.59E-06 | 0.000921 |
| ENSRNOG00000014290 | *Grm1* | 2107.728 | -1.05434 | 0.230114 | -4.58181 | 4.61E-06 | 0.000921 |
| ENSRNOG00000061845 | *Cttnbp2* | 5814.422 | -1.02608 | 0.223912 | -4.5825 | 4.59E-06 | 0.000921 |
| ENSRNOG00000017874 | *Cd53* | 560.9473 | 1.074963 | 0.234756 | 4.579056 | 4.67E-06 | 0.000921 |
| ENSRNOG00000011054 | *Laptm5* | 1693.98 | 0.892394 | 0.195203 | 4.571623 | 4.84E-06 | 0.000942 |
| ENSRNOG00000020991 | *Ms4a6a* | 73.92094 | 1.491165 | 0.327612 | 4.551621 | 5.32E-06 | 0.001024 |
| ENSRNOG00000033893 | *Cacna1h* | 3109.334 | -1.24815 | 0.274481 | -4.5473 | 5.43E-06 | 0.001032 |
| ENSRNOG00000002045 | *Anxa3* | 3860.276 | 1.022825 | 0.225105 | 4.543775 | 5.53E-06 | 0.001036 |
| ENSRNOG00000019716 | *Ntf3* | 536.768 | -2.0437 | 0.453016 | -4.51131 | 6.44E-06 | 0.001184 |
| ENSRNOG00000020485 | *Vav3* | 1224.372 | -2.00002 | 0.443424 | -4.5104 | 6.47E-06 | 0.001184 |
| ENSRNOG00000027230 | *Fhod3* | 481.2656 | 2.2957 | 0.510412 | 4.49774 | 6.87E-06 | 0.001242 |
| ENSRNOG00000004805 | *Stac2* | 1110.094 | 2.359136 | 0.525374 | 4.490391 | 7.11E-06 | 0.001256 |
| ENSRNOG00000005776 | *Bcl11b* | 4156.658 | -1.50656 | 0.335353 | -4.49246 | 7.04E-06 | 0.001256 |
| ENSRNOG00000006649 | *Thrb* | 1754.039 | 1.411608 | 0.314816 | 4.483918 | 7.33E-06 | 0.001279 |
| ENSRNOG00000033734 | *Tnnt2* | 107.2287 | 5.55644 | 1.24012 | 4.480567 | 7.44E-06 | 0.001285 |
| ENSRNOG00000001827 | *Masp1* | 1479.838 | 0.966458 | 0.21641 | 4.46586 | 7.97E-06 | 0.00136 |
| ENSRNOG00000060773 | *Sertad4* | 1056.262 | -1.8787 | 0.420891 | -4.46363 | 8.06E-06 | 0.00136 |
| ENSRNOG00000020652 | *Tgfb1* | 410.5716 | 1.143438 | 0.256462 | 4.458501 | 8.25E-06 | 0.001378 |
| ENSRNOG00000009385 | *Pik3cg* | 318.2916 | 0.982601 | 0.220949 | 4.447185 | 8.70E-06 | 0.001395 |
| ENSRNOG00000021517 | *Tmem231* | 720.4508 | -0.88606 | 0.19927 | -4.44654 | 8.73E-06 | 0.001395 |
| ENSRNOG00000032437 | *Pard3* | 1080.036 | -0.68807 | 0.154661 | -4.44887 | 8.63E-06 | 0.001395 |
| ENSRNOG00000056944 | *Arhgap24* | 188.5457 | 0.893272 | 0.20077 | 4.449223 | 8.62E-06 | 0.001395 |
| ENSRNOG00000046776 | *Msx3* | 26.73332 | -2.53684 | 0.57149 | -4.43899 | 9.04E-06 | 0.00143 |
| ENSRNOG00000006591 | *Fam163b* | 5922.386 | -1.38562 | 0.312534 | -4.4335 | 9.27E-06 | 0.001452 |
| ENSRNOG00000009206 | *Fezf2* | 856.1277 | 0.977197 | 0.221327 | 4.415172 | 1.01E-05 | 0.001524 |
| ENSRNOG00000018951 | *Col4a5* | 469.2719 | 1.251324 | 0.283295 | 4.417028 | 1.00E-05 | 0.001524 |
| ENSRNOG00000025274 | *Hexb* | 4974.31 | 0.53695 | 0.121638 | 4.41433 | 1.01E-05 | 0.001524 |
| ENSRNOG00000026087 | *Igfn1* | 47.056 | 1.86895 | 0.422916 | 4.419197 | 9.91E-06 | 0.001524 |
| ENSRNOG00000002434 | *Tmem100* | 897.3488 | 0.855768 | 0.194562 | 4.398426 | 1.09E-05 | 0.001624 |
| ENSRNOG00000015318 | *Heyl* | 995.3333 | 1.992476 | 0.454229 | 4.3865 | 1.15E-05 | 0.001682 |
| ENSRNOG00000016552 | *Hmgcs1* | 16914.9 | 0.849489 | 0.193626 | 4.387256 | 1.15E-05 | 0.001682 |
| ENSRNOG00000049580 | *Gpr6* | 101.2293 | -3.41395 | 0.779198 | -4.38137 | 1.18E-05 | 0.001706 |
| ENSRNOG00000001115 | *Slc29a4* | 481.3375 | -1.31846 | 0.301886 | -4.36742 | 1.26E-05 | 0.001796 |
| ENSRNOG00000003888 | *Rgs13* | 238.2829 | -2.90682 | 0.665787 | -4.366 | 1.27E-05 | 0.001796 |
| ENSRNOG00000052376 | *Bend6* | 6563.271 | 0.675627 | 0.154939 | 4.360586 | 1.30E-05 | 0.001824 |
| ENSRNOG00000002595 | *Dpp10* | 450.5433 | 1.791934 | 0.412093 | 4.34837 | 1.37E-05 | 0.001877 |
| ENSRNOG00000008758 | *Tspan18* | 594.8324 | -1.59502 | 0.366669 | -4.35002 | 1.36E-05 | 0.001877 |
| ENSRNOG00000058646 | *Zfp36l1* | 1188.739 | 1.025808 | 0.235908 | 4.348341 | 1.37E-05 | 0.001877 |
| ENSRNOG00000008736 | *Slamf8* | 86.6299 | 1.250917 | 0.287899 | 4.344992 | 1.39E-05 | 0.001888 |
| ENSRNOG00000012701 | *Map7* | 3154.81 | -0.91129 | 0.209822 | -4.34318 | 1.40E-05 | 0.001888 |
| ENSRNOG00000009912 | *Fgr* | 69.37473 | 1.493478 | 0.345413 | 4.32375 | 1.53E-05 | 0.002044 |
| ENSRNOG00000009802 | *Fam124a* | 845.6281 | -0.76428 | 0.177015 | -4.31761 | 1.58E-05 | 0.002083 |
| ENSRNOG00000017575 | *Gins2* | 1713.476 | -0.80508 | 0.186681 | -4.31259 | 1.61E-05 | 0.002113 |
| ENSRNOG00000008027 | *Cavin4* | 124.0416 | -1.19518 | 0.277684 | -4.30411 | 1.68E-05 | 0.002177 |
| ENSRNOG00000018198 | *Dapk1* | 3017.618 | -1.14692 | 0.266735 | -4.29982 | 1.71E-05 | 0.002191 |
| ENSRNOG00000056643 | *Cdh8* | 1682.957 | -0.96361 | 0.224155 | -4.29886 | 1.72E-05 | 0.002191 |
| ENSRNOG00000051938 | | 119.6423 | 1.309129 | 0.304709 | 4.296322 | 1.74E-05 | 0.002198 |
| ENSRNOG00000011521 | *Filip1* | 851.5863 | -1.1888 | 0.278223 | -4.27284 | 1.93E-05 | 0.002403 |
| ENSRNOG00000014793 | *Gpr149* | 96.41522 | -2.1768 | 0.509448 | -4.27286 | 1.93E-05 | 0.002403 |
| ENSRNOG00000022116 | *Gjb6* | 2522.749 | 1.007899 | 0.236073 | 4.269439 | 1.96E-05 | 0.00242 |
| ENSRNOG00000011203 | *Farp1* | 1351.408 | 1.055354 | 0.247466 | 4.264651 | 2.00E-05 | 0.002452 |
| ENSRNOG00000019319 | *Fchsd2* | 1265.153 | 0.723565 | 0.169977 | 4.256839 | 2.07E-05 | 0.002505 |
| ENSRNOG00000021938 | *Lrrtm4* | 1270.537 | -1.34116 | 0.315098 | -4.25634 | 2.08E-05 | 0.002505 |
| ENSRNOG00000008142 | *Brpf1* | 1484.889 | -0.90371 | 0.212696 | -4.24885 | 2.15E-05 | 0.00257 |
| ENSRNOG00000015859 | *Chdh* | 82.24286 | 1.466488 | 0.345471 | 4.244898 | 2.19E-05 | 0.002595 |
| ENSRNOG00000014010 | *Gfra2* | 215.3203 | 2.612866 | 0.61619 | 4.240355 | 2.23E-05 | 0.002609 |
| ENSRNOG00000021090 | *Pygm* | 2544.759 | 1.161308 | 0.273879 | 4.240223 | 2.23E-05 | 0.002609 |
| ENSRNOG00000055178 | | 30.13754 | -2.65235 | 0.625879 | -4.2378 | 2.26E-05 | 0.002617 |
| ENSRNOG00000042326 | *Smpdl3b* | 707.491 | -1.38178 | 0.326678 | -4.22979 | 2.34E-05 | 0.002692 |
| ENSRNOG00000018366 | *RGD1310819* | 11854.72 | -1.33223 | 0.315513 | -4.22242 | 2.42E-05 | 0.00276 |
| ENSRNOG00000017976 | *Slco2b1* | 1354.853 | 0.98007 | 0.232553 | 4.214389 | 2.50E-05 | 0.002777 |
| ENSRNOG00000030111 | *Cyp11b3* | 32.37584 | 9.586961 | 2.274802 | 4.214415 | 2.50E-05 | 0.002777 |
| ENSRNOG00000030111 | *Cyp11b2* | 32.37584 | 9.586961 | 2.274802 | 4.214415 | 2.50E-05 | 0.002777 |
| ENSRNOG00000030111 | *LOC100910462* | 32.37584 | 9.586961 | 2.274802 | 4.214415 | 2.50E-05 | 0.002777 |
| ENSRNOG00000030715 | *Cfh* | 1877.615 | 0.917544 | 0.217521 | 4.218184 | 2.46E-05 | 0.002777 |
| ENSRNOG00000052080 | *Camk2b* | 61094.43 | -1.04959 | 0.249014 | -4.21498 | 2.50E-05 | 0.002777 |
| ENSRNOG00000007189 | *Ttc22* | 314.8748 | 1.860858 | 0.442359 | 4.206665 | 2.59E-05 | 0.002847 |
| ENSRNOG00000055305 | *Parvb* | 2378.447 | 1.110856 | 0.264147 | 4.205452 | 2.61E-05 | 0.002847 |
| ENSRNOG00000053200 | *Dock10* | 4304.324 | -1.38689 | 0.330227 | -4.19982 | 2.67E-05 | 0.002898 |
| ENSRNOG00000023688 | *Drd1* | 174.6735 | -2.49519 | 0.59456 | -4.19669 | 2.71E-05 | 0.002918 |
| ENSRNOG00000003330 | *Acsf2* | 1654.198 | 1.082242 | 0.258559 | 4.185664 | 2.84E-05 | 0.003041 |
| ENSRNOG00000008465 | *Tmem176b* | 3258.007 | 0.866569 | 0.207368 | 4.178898 | 2.93E-05 | 0.003111 |
| ENSRNOG00000011231 | *Cntnap4* | 343.2908 | 1.729266 | 0.41482 | 4.168714 | 3.06E-05 | 0.003191 |
| ENSRNOG00000016371 | *Slc18b1* | 846.1953 | -0.63684 | 0.152777 | -4.16842 | 3.07E-05 | 0.003191 |
| ENSRNOG00000018414 | *Csf1r* | 4214.282 | 0.86601 | 0.207606 | 4.171402 | 3.03E-05 | 0.003191 |
| ENSRNOG00000006116 | *Hk2* | 531.5656 | 0.922171 | 0.221331 | 4.166487 | 3.09E-05 | 0.003196 |
| ENSRNOG00000001851 | *Far2* | 1043.441 | 0.590306 | 0.141794 | 4.163112 | 3.14E-05 | 0.003222 |
| ENSRNOG00000029242 | *Plekho2* | 260.3022 | 1.373829 | 0.330138 | 4.16138 | 3.16E-05 | 0.003225 |
| ENSRNOG00000010065 | *Dgkh* | 183.6863 | -2.16272 | 0.519977 | -4.15925 | 3.19E-05 | 0.003233 |
| ENSRNOG00000048187 | *Epop* | 351.898 | 2.158254 | 0.519312 | 4.155991 | 3.24E-05 | 0.003258 |
| ENSRNOG00000014166 | *Smoc2* | 88.56894 | -2.4923 | 0.60365 | -4.12872 | 3.65E-05 | 0.003645 |
| ENSRNOG00000012531 | *Ephb2* | 684.9783 | -1.00111 | 0.24262 | -4.12625 | 3.69E-05 | 0.003661 |
| ENSRNOG00000004180 | *Tafa2* | 3543.349 | -1.48101 | 0.359605 | -4.11844 | 3.81E-05 | 0.003738 |
| ENSRNOG00000013231 | *Ptafr* | 454.5691 | 1.050808 | 0.25508 | 4.119527 | 3.80E-05 | 0.003738 |
| ENSRNOG00000022822 | *Fam169b* | 135.9214 | 1.133691 | 0.275871 | 4.109499 | 3.97E-05 | 0.003861 |
| ENSRNOG00000004409 | *Sash3* | 160.9852 | 1.219945 | 0.297485 | 4.100869 | 4.12E-05 | 0.003867 |
| ENSRNOG00000004863 | *Mpped2* | 2618.136 | -1.1153 | 0.27171 | -4.10475 | 4.05E-05 | 0.003867 |
| ENSRNOG00000005825 | *Lyz2* | 2249.615 | 1.482697 | 0.361187 | 4.105067 | 4.04E-05 | 0.003867 |
| ENSRNOG00000015594 | *Rftn2* | 715.1176 | 0.926327 | 0.225877 | 4.101031 | 4.11E-05 | 0.003867 |
| ENSRNOG00000017444 | *Nrsn1* | 10757.61 | 1.475535 | 0.360224 | 4.096163 | 4.20E-05 | 0.003867 |
| ENSRNOG00000024207 | *Fgfrl1* | 664.7516 | 0.625141 | 0.152601 | 4.096558 | 4.19E-05 | 0.003867 |
| ENSRNOG00000031058 | *Was* | 78.75683 | 1.561485 | 0.380537 | 4.10337 | 4.07E-05 | 0.003867 |
| ENSRNOG00000055451 | | 98.71152 | -3.1837 | 0.776559 | -4.09975 | 4.14E-05 | 0.003867 |
| ENSRNOG00000057044 | *Garnl3* | 828.4141 | 2.334703 | 0.569839 | 4.097128 | 4.18E-05 | 0.003867 |
| ENSRNOG00000004613 | *Gpm6b* | 24664.58 | 0.700677 | 0.171481 | 4.086041 | 4.39E-05 | 0.003991 |
| ENSRNOG00000005342 | *Rassf5* | 438.6055 | 1.767433 | 0.432486 | 4.086687 | 4.38E-05 | 0.003991 |
| ENSRNOG00000002922 | *Adora2b* | 374.3201 | 0.892357 | 0.219005 | 4.074601 | 4.61E-05 | 0.004167 |
| ENSRNOG00000016977 | *Calb2* | 532.4148 | 1.14312 | 0.280859 | 4.070082 | 4.70E-05 | 0.004224 |
| ENSRNOG00000020337 | *Sla2* | 34.142 | -1.76581 | 0.434322 | -4.06568 | 4.79E-05 | 0.004271 |
| ENSRNOG00000042888 | *Cdyl2* | 54.67536 | 1.242698 | 0.305725 | 4.064753 | 4.81E-05 | 0.004271 |
| ENSRNOG00000027869 | *Sox5* | 262.5494 | 1.5736 | 0.387291 | 4.063096 | 4.84E-05 | 0.004276 |
| ENSRNOG00000007324 | *Plxna2* | 3610.994 | -0.95314 | 0.234748 | -4.06028 | 4.90E-05 | 0.004303 |
| ENSRNOG00000014259 | *Mycl* | 1214.568 | -1.30225 | 0.321033 | -4.05643 | 4.98E-05 | 0.004325 |
| ENSRNOG00000015845 | *Niban2* | 2050.464 | 0.825741 | 0.203559 | 4.056522 | 4.98E-05 | 0.004325 |
| ENSRNOG00000001271 | *Card6* | 195.0983 | 1.041619 | 0.256995 | 4.053069 | 5.06E-05 | 0.004363 |
| ENSRNOG00000057578 | *Prodh2* | 61.99851 | 2.180435 | 0.538944 | 4.045753 | 5.22E-05 | 0.00445 |
| ENSRNOG00000060614 | *Pxdn* | 541.0551 | -1.99009 | 0.491794 | -4.0466 | 5.20E-05 | 0.00445 |
| ENSRNOG00000007262 | *Ccdc134* | 488.944 | -1.31233 | 0.325589 | -4.03064 | 5.56E-05 | 0.004568 |
| ENSRNOG00000008620 | *Smad3* | 2993.088 | -1.16127 | 0.287743 | -4.03579 | 5.44E-05 | 0.004568 |
| ENSRNOG00000008759 | *Csf3r* | 164.2852 | 0.979834 | 0.243175 | 4.029345 | 5.59E-05 | 0.004568 |
| ENSRNOG00000010938 | *Slc7a10* | 1199.113 | 1.054411 | 0.261363 | 4.034274 | 5.48E-05 | 0.004568 |
| ENSRNOG00000014786 | *Ccne1* | 536.4418 | -0.81739 | 0.202781 | -4.03092 | 5.56E-05 | 0.004568 |
| ENSRNOG00000019229 | *Fbl* | 912.0663 | -0.89178 | 0.221176 | -4.03198 | 5.53E-05 | 0.004568 |
| ENSRNOG00000028082 | *Tal2* | 16.48052 | -2.94658 | 0.731286 | -4.02932 | 5.59E-05 | 0.004568 |
| ENSRNOG00000034015 | *Capn2* | 4284.632 | 0.618733 | 0.153382 | 4.033934 | 5.49E-05 | 0.004568 |
| ENSRNOG00000011202 | *Chrna4* | 393.1 | 2.558273 | 0.635782 | 4.023822 | 5.73E-05 | 0.004602 |
| ENSRNOG00000013654 | *Cbln2* | 344.9787 | 2.026562 | 0.503638 | 4.023845 | 5.73E-05 | 0.004602 |
| ENSRNOG00000023337 | *Sema3a* | 14.57117 | 4.71426 | 1.171455 | 4.024278 | 5.72E-05 | 0.004602 |
| ENSRNOG00000039471 | *LOC108348771* | 94.84385 | -1.42432 | 0.354143 | -4.02189 | 5.77E-05 | 0.004615 |
| ENSRNOG00000013907 | *Sall1* | 1119.63 | 0.685029 | 0.170518 | 4.017345 | 5.89E-05 | 0.004667 |
| ENSRNOG00000018550 | *Ncbp3* | 710.8705 | -0.6302 | 0.156891 | -4.01679 | 5.90E-05 | 0.004667 |
| ENSRNOG00000018564 | *Nup93* | 442.0259 | -0.89751 | 0.223694 | -4.01223 | 6.01E-05 | 0.004709 |
| ENSRNOG00000020325 | *Calhm2* | 514.2159 | -0.95929 | 0.239082 | -4.01238 | 6.01E-05 | 0.004709 |
| ENSRNOG00000000595 | *Traf3ip2* | 65.86301 | 1.220389 | 0.304327 | 4.010119 | 6.07E-05 | 0.004727 |
| ENSRNOG00000026091 | *Slc10a4* | 34.08048 | -2.46319 | 0.614964 | -4.00542 | 6.19E-05 | 0.004797 |
| ENSRNOG00000011381 | *Acsbg1* | 10101.88 | 0.782598 | 0.195547 | 4.002101 | 6.28E-05 | 0.004841 |
| ENSRNOG00000013886 | *Fyb1* | 136.1368 | 1.072065 | 0.268066 | 3.999264 | 6.35E-05 | 0.004874 |
| ENSRNOG00000037196 | *Spag17* | 32.44612 | -3.08152 | 0.771059 | -3.99648 | 6.43E-05 | 0.004907 |
| ENSRNOG00000021062 | *Fxyd5* | 373.5244 | 1.113312 | 0.278842 | 3.99263 | 6.53E-05 | 0.004963 |
| ENSRNOG00000012580 | *Ccdc141* | 675.541 | 1.139551 | 0.285539 | 3.990869 | 6.58E-05 | 0.004972 |
| ENSRNOG00000025624 | *Arhgap20* | 3206.38 | -0.93866 | 0.235266 | -3.9898 | 6.61E-05 | 0.004972 |
| ENSRNOG00000050869 | *Cebpd* | 543.7884 | -1.43324 | 0.359669 | -3.98488 | 6.75E-05 | 0.005048 |
| ENSRNOG00000059215 | *Slc66a2* | 1509.352 | -0.8771 | 0.220162 | -3.9839 | 6.78E-05 | 0.005048 |
| ENSRNOG00000001065 | *Cyth3* | 1632.714 | -0.84613 | 0.212853 | -3.97519 | 7.03E-05 | 0.005111 |
| ENSRNOG00000009401 | *Lmo2* | 1011.619 | -0.91369 | 0.229779 | -3.97637 | 7.00E-05 | 0.005111 |
| ENSRNOG00000017477 | *Mmp23* | 165.4341 | -1.76077 | 0.44289 | -3.97563 | 7.02E-05 | 0.005111 |
| ENSRNOG00000020300 | *Lsp1* | 257.3937 | 1.808644 | 0.454812 | 3.976685 | 6.99E-05 | 0.005111 |
| ENSRNOG00000022772 | *Prickle1* | 3649.706 | -0.82161 | 0.206502 | -3.97869 | 6.93E-05 | 0.005111 |
| ENSRNOG00000014481 | *Adtrp* | 108.418 | 1.379324 | 0.347382 | 3.97063 | 7.17E-05 | 0.005185 |
| ENSRNOG00000034037 | *Zfp266* | 2388.079 | -0.74677 | 0.188225 | -3.96744 | 7.26E-05 | 0.00523 |
| ENSRNOG00000008113 | *RGD1561149* | 415.1763 | 3.197522 | 0.806504 | 3.964672 | 7.35E-05 | 0.005241 |
| ENSRNOG00000010188 | *Satb2* | 503.0841 | 2.431902 | 0.613224 | 3.965763 | 7.32E-05 | 0.005241 |
| ENSRNOG00000002336 | *Gabra4* | 6194.746 | -0.96328 | 0.243465 | -3.95654 | 7.60E-05 | 0.005395 |
| ENSRNOG00000057996 | *Phactr3* | 9015.757 | 0.619857 | 0.156706 | 3.955531 | 7.64E-05 | 0.005395 |
| ENSRNOG00000021157 | *Ctss* | 11070.5 | 0.789417 | 0.2 | 3.947075 | 7.91E-05 | 0.005563 |
| ENSRNOG00000013160 | *Sash1* | 3120.861 | 0.696998 | 0.176846 | 3.941265 | 8.11E-05 | 0.005647 |
| ENSRNOG00000046984 | *St6galnac6* | 6670.08 | 0.708973 | 0.179838 | 3.942284 | 8.07E-05 | 0.005647 |
| ENSRNOG00000042731 | *Slc25a18* | 3164.674 | 1.512052 | 0.383764 | 3.940051 | 8.15E-05 | 0.00565 |
| ENSRNOG00000016692 | *Hsdl2* | 1999.069 | 0.64245 | 0.163539 | 3.928419 | 8.55E-05 | 0.005903 |
| ENSRNOG00000003800 | *Rgs9* | 500.4486 | -3.48897 | 0.889096 | -3.92418 | 8.70E-05 | 0.005981 |
| ENSRNOG00000019965 | *Tgfb1i1* | 699.5654 | 0.817746 | 0.208533 | 3.92142 | 8.80E-05 | 0.006023 |
| ENSRNOG00000060087 | *Adra1b* | 57.03726 | 2.876857 | 0.73443 | 3.917128 | 8.96E-05 | 0.006104 |
| ENSRNOG00000005166 | *Nhlh1* | 559.0869 | -2.38798 | 0.609969 | -3.91492 | 9.04E-05 | 0.006132 |
| ENSRNOG00000008428 | *Drd2* | 165.0874 | -3.77924 | 0.966958 | -3.90838 | 9.29E-05 | 0.006272 |
| ENSRNOG00000007993 | *Sh3tc1* | 107.6263 | 1.307203 | 0.334753 | 3.904982 | 9.42E-05 | 0.006333 |
| ENSRNOG00000016525 | *Susd3* | 202.3314 | 1.190202 | 0.304926 | 3.903249 | 9.49E-05 | 0.006351 |
| ENSRNOG00000037299 | *Smpd5* | 77.3984 | -1.22914 | 0.315405 | -3.89702 | 9.74E-05 | 0.006487 |
| ENSRNOG00000014232 | *P2ry1* | 108.4497 | -1.37324 | 0.352666 | -3.89389 | 9.87E-05 | 0.006543 |
| ENSRNOG00000001518 | *Itga6* | 1537.953 | 0.744623 | 0.19134 | 3.89162 | 9.96E-05 | 0.006576 |
| ENSRNOG00000011568 | *Rspo3* | 907.4418 | -1.11799 | 0.287457 | -3.88925 | 0.000101 | 0.006612 |
| ENSRNOG00000008941 | *Ets1* | 536.0988 | 0.949369 | 0.244985 | 3.875212 | 0.000107 | 0.006975 |
| ENSRNOG00000012795 | *Cables1* | 1011.54 | 1.206412 | 0.31166 | 3.870929 | 0.000108 | 0.007068 |
| ENSRNOG00000047545 | *Adra2a* | 291.3333 | 2.467092 | 0.637516 | 3.86985 | 0.000109 | 0.007069 |
| ENSRNOG00000019885 | *Magi3* | 1449.847 | 0.700403 | 0.181053 | 3.868492 | 0.00011 | 0.007071 |
| ENSRNOG00000027264 | *Dagla* | 6229.52 | -1.39178 | 0.359941 | -3.86668 | 0.00011 | 0.007071 |
| ENSRNOG00000057556 | *Pdzrn3* | 691.1722 | 1.209601 | 0.312785 | 3.867198 | 0.00011 | 0.007071 |
| ENSRNOG00000028404 | *Ppp1r1b* | 6621.46 | -1.98971 | 0.515189 | -3.8621 | 0.000112 | 0.007174 |
| ENSRNOG00000007477 | *Edn3* | 126.8823 | 1.343937 | 0.3481 | 3.86078 | 0.000113 | 0.007183 |
| ENSRNOG00000062181 | | 2150.938 | -0.95093 | 0.246893 | -3.85159 | 0.000117 | 0.007427 |
| ENSRNOG00000000306 | *Smpd2* | 2113.317 | -1.215 | 0.31565 | -3.84919 | 0.000119 | 0.007469 |
| ENSRNOG00000005299 | *Kif5a* | 50238.91 | 1.287434 | 0.334632 | 3.847318 | 0.000119 | 0.007495 |
| ENSRNOG00000046536 | | 245.2507 | 0.918714 | 0.238877 | 3.845972 | 0.00012 | 0.007505 |
| ENSRNOG00000059834 | *Cib2* | 2031.536 | 1.049295 | 0.273717 | 3.833508 | 0.000126 | 0.007864 |
| ENSRNOG00000000302 | *Sesn1* | 2851.496 | -0.8481 | 0.221753 | -3.82453 | 0.000131 | 0.008123 |
| ENSRNOG00000005686 | *Suclg2* | 3356.465 | 0.883167 | 0.231211 | 3.819747 | 0.000134 | 0.008248 |
| ENSRNOG00000046700 | *Mettl27* | 148.7967 | 0.872489 | 0.228812 | 3.813131 | 0.000137 | 0.008438 |
| ENSRNOG00000036661 | *Rab40b* | 3562.039 | -0.98879 | 0.259914 | -3.80428 | 0.000142 | 0.00871 |
| ENSRNOG00000043186 | *Ppil6* | 280.5399 | -1.18125 | 0.311135 | -3.7966 | 0.000147 | 0.008948 |
| ENSRNOG00000014453 | *Anxa5* | 2677.973 | 0.854566 | 0.225615 | 3.787726 | 0.000152 | 0.009237 |
| ENSRNOG00000010775 | *Arrdc4* | 331.471 | -0.77173 | 0.203947 | -3.78394 | 0.000154 | 0.009304 |
| ENSRNOG00000013321 | *Dock11* | 796.2093 | -0.97141 | 0.256696 | -3.78428 | 0.000154 | 0.009304 |
| ENSRNOG00000001259 | *Cux2* | 547.7272 | 1.800488 | 0.477055 | 3.774173 | 0.000161 | 0.009519 |
| ENSRNOG00000007069 | *Adhfe1* | 2353.83 | 0.883112 | 0.233913 | 3.775385 | 0.00016 | 0.009519 |
| ENSRNOG00000012789 | *Mlnr* | 13.27084 | 7.42462 | 1.968119 | 3.772445 | 0.000162 | 0.009519 |
| ENSRNOG00000016069 | *Cd3e* | 445.6339 | -7.78668 | 2.062835 | -3.77475 | 0.00016 | 0.009519 |
| ENSRNOG00000031939 | | 167.0866 | 1.175306 | 0.311555 | 3.772391 | 0.000162 | 0.009519 |
| ENSRNOG00000051303 | | 3089.854 | -0.65655 | 0.173899 | -3.77549 | 0.00016 | 0.009519 |
| ENSRNOG00000029292 | *LOC500877* | 44.42607 | -2.03794 | 0.540835 | -3.76814 | 0.000164 | 0.009645 |
| ENSRNOG00000019138 | *Clec11a* | 1070.378 | 0.942696 | 0.250377 | 3.765105 | 0.000166 | 0.009726 |
| ENSRNOG00000020702 | *Cyb561a3* | 198.7718 | 0.783539 | 0.208454 | 3.758813 | 0.000171 | 0.009935 |
| ENSRNOG00000013681 | *Kcns1* | 255.9372 | 1.554625 | 0.413918 | 3.755878 | 0.000173 | 0.009938 |
| ENSRNOG00000019077 | *Lipa* | 1971.412 | 0.748847 | 0.199377 | 3.755925 | 0.000173 | 0.009938 |
| ENSRNOG00000057367 | *Glud1* | 19132.03 | 0.804135 | 0.214079 | 3.75626 | 0.000172 | 0.009938 |
| ENSRNOG00000019321 | *Cck* | 7446.509 | 1.647055 | 0.438965 | 3.752135 | 0.000175 | 0.01005 |
| ENSRNOG00000004889 | *Vangl2* | 278.0251 | -1.28666 | 0.34304 | -3.75077 | 0.000176 | 0.010057 |
| ENSRNOG00000023969 | *Herc6* | 279.8593 | 1.124238 | 0.299791 | 3.750069 | 0.000177 | 0.010057 |
| ENSRNOG00000018797 | *Myrip* | 3081.604 | 0.86231 | 0.230083 | 3.74782 | 0.000178 | 0.01011 |
| ENSRNOG00000003738 | *Ush2a* | 62.43995 | -1.8889 | 0.504974 | -3.74058 | 0.000184 | 0.010309 |
| ENSRNOG00000014702 | *Elovl2* | 3303.484 | 0.742366 | 0.198487 | 3.74013 | 0.000184 | 0.010309 |
| ENSRNOG00000042848 | *Jam2* | 2999.729 | 0.789127 | 0.210955 | 3.740736 | 0.000183 | 0.010309 |
| ENSRNOG00000011778 | *Blvra* | 784.0758 | 0.725951 | 0.194196 | 3.738235 | 0.000185 | 0.010348 |
| ENSRNOG00000010440 | *Gnal* | 1131.085 | -1.31598 | 0.35223 | -3.73613 | 0.000187 | 0.010397 |
| ENSRNOG00000049939 | | 665.2732 | 1.073615 | 0.28745 | 3.734957 | 0.000188 | 0.010408 |
| ENSRNOG00000003936 | *Pwwp2a* | 1205.093 | -1.08211 | 0.290037 | -3.73094 | 0.000191 | 0.010537 |
| ENSRNOG00000013578 | *Trem2* | 883.1349 | 1.100066 | 0.295116 | 3.727568 | 0.000193 | 0.01064 |
| ENSRNOG00000008203 | *Synpr* | 4102.058 | -1.13091 | 0.303608 | -3.7249 | 0.000195 | 0.010676 |
| ENSRNOG00000014870 | *Slc13a5* | 452.8112 | 1.322332 | 0.354947 | 3.725438 | 0.000195 | 0.010676 |
| ENSRNOG00000011841 | *Map2* | 60556.83 | 0.903814 | 0.242843 | 3.721797 | 0.000198 | 0.01073 |
| ENSRNOG00000018683 | *Dock1* | 1261.808 | 0.629829 | 0.169199 | 3.722413 | 0.000197 | 0.01073 |
| ENSRNOG00000007089 | *Lgmn* | 3816.424 | 0.472876 | 0.127379 | 3.712368 | 0.000205 | 0.011068 |
| ENSRNOG00000013322 | *Pola1* | 248.797 | -0.97437 | 0.26248 | -3.71216 | 0.000205 | 0.011068 |
| ENSRNOG00000008187 | *Ubash3b* | 1340.148 | -1.12986 | 0.304673 | -3.70845 | 0.000209 | 0.011074 |
| ENSRNOG00000009982 | *Pnp* | 4426.572 | 0.678167 | 0.182904 | 3.707769 | 0.000209 | 0.011074 |
| ENSRNOG00000017208 | *Cspg4* | 815.5998 | 0.921459 | 0.248533 | 3.707589 | 0.000209 | 0.011074 |
| ENSRNOG00000030285 | *Epha3* | 540.8939 | -1.12697 | 0.303911 | -3.70823 | 0.000209 | 0.011074 |
| ENSRNOG00000047080 | *Gng4* | 1090.148 | 1.372867 | 0.369979 | 3.710658 | 0.000207 | 0.011074 |
| ENSRNOG00000046863 | *C1ql2* | 5582.444 | -1.95879 | 0.529234 | -3.70117 | 0.000215 | 0.011318 |
| ENSRNOG00000004414 | *F11r* | 259.1442 | 0.996022 | 0.26945 | 3.696498 | 0.000219 | 0.011489 |
| ENSRNOG00000058589 | | 89.73633 | 2.635503 | 0.713858 | 3.691913 | 0.000223 | 0.011657 |
| ENSRNOG00000008086 | *Dpf3* | 166.7754 | -1.02615 | 0.278371 | -3.68626 | 0.000228 | 0.011838 |
| ENSRNOG00000010301 | *Eif4e3* | 857.3448 | -0.70306 | 0.190684 | -3.68703 | 0.000227 | 0.011838 |
| ENSRNOG00000060693 | *Catsperg* | 84.42276 | 2.474223 | 0.672455 | 3.679387 | 0.000234 | 0.01212 |
| ENSRNOG00000004821 | *Sntb1* | 103.5585 | 1.706233 | 0.463868 | 3.678273 | 0.000235 | 0.012123 |
| ENSRNOG00000009967 | *Otof* | 558.3512 | 1.997921 | 0.54327 | 3.677582 | 0.000235 | 0.012123 |
| ENSRNOG00000002360 | *Gabrg1* | 1671.942 | 1.009406 | 0.274602 | 3.675885 | 0.000237 | 0.012163 |
| ENSRNOG00000058271 | *Chpt1* | 1485.578 | 0.770581 | 0.20971 | 3.674499 | 0.000238 | 0.012188 |
| ENSRNOG00000017097 | *Lhpp* | 2415.235 | 0.796548 | 0.217112 | 3.668841 | 0.000244 | 0.012411 |
| ENSRNOG00000017803 | *Apbb1ip* | 586.3677 | 0.900494 | 0.24549 | 3.668145 | 0.000244 | 0.012411 |
| ENSRNOG00000039300 | *Ahcyl2* | 5846.412 | -0.88813 | 0.242233 | -3.66641 | 0.000246 | 0.012454 |
| ENSRNOG00000018092 | *Cd83* | 1043.858 | 0.891105 | 0.243293 | 3.662681 | 0.00025 | 0.012595 |
| ENSRNOG00000042690 | *Zmat4* | 1285.73 | -0.94233 | 0.257368 | -3.6614 | 0.000251 | 0.012616 |
| ENSRNOG00000004171 | *Dnah9* | 936.2654 | -1.37194 | 0.374966 | -3.65883 | 0.000253 | 0.012701 |
| ENSRNOG00000019570 | *Gng3* | 9033.596 | 0.905291 | 0.247655 | 3.655453 | 0.000257 | 0.012827 |
| ENSRNOG00000008323 | *Pitpnm3* | 1179.428 | 1.919584 | 0.525382 | 3.653695 | 0.000258 | 0.012873 |
| ENSRNOG00000011094 | *Efcab6* | 215.4268 | 1.227926 | 0.336382 | 3.650393 | 0.000262 | 0.012997 |
| ENSRNOG00000005865 | *Itprid2* | 1966.955 | 0.704924 | 0.193541 | 3.642245 | 0.00027 | 0.013242 |
| ENSRNOG00000013248 | *Wwc2* | 1261.921 | 0.437193 | 0.120015 | 3.642809 | 0.00027 | 0.013242 |
| ENSRNOG00000016288 | *Tcea2* | 5491.716 | 0.703474 | 0.193068 | 3.64366 | 0.000269 | 0.013242 |
| ENSRNOG00000019662 | *Tm6sf1* | 825.1123 | 0.629027 | 0.172643 | 3.643513 | 0.000269 | 0.013242 |
| ENSRNOG00000010575 | *Dapp1* | 142.1718 | 1.019437 | 0.279975 | 3.641173 | 0.000271 | 0.013255 |
| ENSRNOG00000060496 | *Kdm5d* | 1366.958 | -0.46622 | 0.128116 | -3.63901 | 0.000274 | 0.013324 |
| ENSRNOG00000017510 | *Mfge8* | 7628.228 | 1.09439 | 0.300884 | 3.637254 | 0.000276 | 0.013372 |
| ENSRNOG00000007234 | *Cyp51* | 10086.82 | 0.684351 | 0.188401 | 3.632417 | 0.000281 | 0.013582 |
| ENSRNOG00000017689 | *Itih3* | 9251.31 | 1.642218 | 0.452622 | 3.628232 | 0.000285 | 0.01376 |
| ENSRNOG00000003005 | *Rasgef1c* | 136.1529 | 3.531893 | 0.973808 | 3.626887 | 0.000287 | 0.013788 |
| ENSRNOG00000002278 | *Tec* | 149.3211 | 1.820182 | 0.502456 | 3.622568 | 0.000292 | 0.013894 |
| ENSRNOG00000003209 | *Pcp4l1* | 1180.635 | -2.06378 | 0.569709 | -3.62252 | 0.000292 | 0.013894 |
| ENSRNOG00000005248 | *Slc1a4* | 1570.664 | 0.749244 | 0.206832 | 3.622469 | 0.000292 | 0.013894 |
| ENSRNOG00000002106 | *Usp46* | 5366.193 | 0.455534 | 0.125822 | 3.620478 | 0.000294 | 0.013958 |
| ENSRNOG00000016595 | *Hhex* | 134.5489 | 1.389056 | 0.384404 | 3.613537 | 0.000302 | 0.014292 |
| ENSRNOG00000000991 | *Arpc1b* | 523.1566 | 1.016119 | 0.281274 | 3.612565 | 0.000303 | 0.014301 |
| ENSRNOG00000004442 | *Dglucy* | 223.5625 | 0.844436 | 0.233816 | 3.611536 | 0.000304 | 0.014306 |
| ENSRNOG00000021682 | *Dcakd* | 1844.506 | -0.60674 | 0.168032 | -3.61088 | 0.000305 | 0.014306 |
| ENSRNOG00000061450 | *Homer2* | 426.8051 | 0.841577 | 0.233175 | 3.609205 | 0.000307 | 0.014354 |
| ENSRNOG00000057347 | *Cebpb* | 866.5396 | -1.39636 | 0.386981 | -3.60835 | 0.000308 | 0.014357 |
| ENSRNOG00000009656 | *Rspo1* | 273.7771 | 3.261632 | 0.904129 | 3.607485 | 0.000309 | 0.014361 |
| ENSRNOG00000011320 | *Igfbpl1* | 65.39115 | -2.07742 | 0.576429 | -3.60395 | 0.000313 | 0.014514 |
| ENSRNOG00000011162 | *Smco4* | 284.5717 | -1.0879 | 0.301954 | -3.60287 | 0.000315 | 0.01453 |
| ENSRNOG00000009946 | *Ldlr* | 1163.003 | -0.83924 | 0.233081 | -3.60066 | 0.000317 | 0.01461 |
| ENSRNOG00000058586 | *Tyro3* | 6444.985 | 0.888427 | 0.246966 | 3.597369 | 0.000321 | 0.014751 |
| ENSRNOG00000032569 | *Lingo3* | 391.6174 | -2.11838 | 0.589321 | -3.59461 | 0.000325 | 0.014863 |
| ENSRNOG00000016737 | *Tcerg1l* | 244.9255 | 3.981315 | 1.107983 | 3.5933 | 0.000327 | 0.014893 |
| ENSRNOG00000001714 | *Atp13a4* | 634.4898 | 1.082479 | 0.301573 | 3.589444 | 0.000331 | 0.014954 |
| ENSRNOG00000005214 | *Plek* | 367.3465 | 1.032096 | 0.287462 | 3.590371 | 0.00033 | 0.014954 |
| ENSRNOG00000005609 | *Neurod1* | 1078.342 | -1.17402 | 0.327104 | -3.58913 | 0.000332 | 0.014954 |
| ENSRNOG00000018250 | *Tnni3* | 38.26121 | -3.35278 | 0.933728 | -3.59074 | 0.00033 | 0.014954 |
| ENSRNOG00000015334 | *Fcho2* | 2615.698 | 0.695745 | 0.193925 | 3.5877 | 0.000334 | 0.014964 |
| ENSRNOG00000016267 | *Chst15* | 382.5638 | -2.04023 | 0.568843 | -3.58664 | 0.000335 | 0.014964 |
| ENSRNOG00000024119 | *Ghsr* | 86.34211 | -2.01938 | 0.562914 | -3.58736 | 0.000334 | 0.014964 |
| ENSRNOG00000029528 | *Cbs* | 3675.089 | 0.973834 | 0.271577 | 3.585855 | 0.000336 | 0.014965 |
| ENSRNOG00000019913 | *Mta2* | 1769.345 | -0.54213 | 0.151353 | -3.58189 | 0.000341 | 0.01515 |
| ENSRNOG00000058609 | | 794.1245 | 0.977307 | 0.272991 | 3.579992 | 0.000344 | 0.015216 |
| ENSRNOG00000015051 | *Golga7b* | 4089.455 | 0.945747 | 0.264309 | 3.578183 | 0.000346 | 0.015277 |
| ENSRNOG00000047102 | *Popdc3* | 67.48913 | 3.067661 | 0.85758 | 3.577112 | 0.000347 | 0.015295 |
| ENSRNOG00000032297 | *Msmo1* | 6727.003 | 0.630257 | 0.176308 | 3.574751 | 0.000351 | 0.015389 |
| ENSRNOG00000017419 | *Map3k4* | 1507.789 | -0.86035 | 0.240875 | -3.57177 | 0.000355 | 0.015478 |
| ENSRNOG00000061484 | *Adamts2* | 59.43224 | 2.698044 | 0.755389 | 3.57173 | 0.000355 | 0.015478 |
| ENSRNOG00000008180 | *Lyn* | 643.9764 | 0.904671 | 0.253624 | 3.566979 | 0.000361 | 0.015672 |
| ENSRNOG00000012185 | *Zfyve21* | 938.4854 | 0.605873 | 0.16985 | 3.567116 | 0.000361 | 0.015672 |
| ENSRNOG00000011452 | *Aldoc* | 51967.39 | 0.812931 | 0.228155 | 3.563068 | 0.000367 | 0.015817 |
| ENSRNOG00000018644 | *Slc6a7* | 1809.534 | 1.311208 | 0.367968 | 3.563378 | 0.000366 | 0.015817 |
| ENSRNOG00000007679 | *Cyth4* | 726.5897 | 0.780219 | 0.219261 | 3.558405 | 0.000373 | 0.016054 |
| ENSRNOG00000014970 | *Kif17* | 610.9578 | -0.81847 | 0.230088 | -3.55719 | 0.000375 | 0.016084 |
| ENSRNOG00000048172 | *Rac3* | 1105.582 | 0.795377 | 0.223705 | 3.555471 | 0.000377 | 0.016143 |
| ENSRNOG00000010624 | *Ap3b1* | 949.5473 | -0.66782 | 0.187882 | -3.55446 | 0.000379 | 0.01616 |
| ENSRNOG00000021068 | *Pip5k1a* | 3349.971 | -0.59789 | 0.168673 | -3.5447 | 0.000393 | 0.016646 |
| ENSRNOG00000033261 | *Fam107a* | 36993.78 | 0.924888 | 0.26094 | 3.544441 | 0.000393 | 0.016646 |
| ENSRNOG00000058805 | *Aida* | 2131.93 | 0.654266 | 0.184518 | 3.545814 | 0.000391 | 0.016646 |
| **FLU vs. Control** | | | | | | | |
| ENSRNOG00000049580 | *Gpr6* | 101.2293 | -4.46222 | 0.787576 | -5.66576 | 1.46E-08 | 0.000295 |
| ENSRNOG00000001302 | *Adora2a* | 368.615 | -4.29658 | 0.84135 | -5.10676 | 3.28E-07 | 0.002817 |
| ENSRNOG00000005639 | *Ar* | 105.8073 | 1.982227 | 0.402379 | 4.926273 | 8.38E-07 | 0.002817 |
| ENSRNOG00000007300 | *C1qtnf6* | 97.94694 | -1.74779 | 0.347829 | -5.02486 | 5.04E-07 | 0.002817 |
| ENSRNOG00000008428 | *Drd2* | 165.0874 | -4.80142 | 0.971471 | -4.94242 | 7.72E-07 | 0.002817 |
| ENSRNOG00000023688 | *Drd1* | 174.6735 | -2.95744 | 0.596668 | -4.95659 | 7.17E-07 | 0.002817 |
| ENSRNOG00000003209 | *Pcp4l1* | 1180.635 | -2.69676 | 0.570083 | -4.73047 | 2.24E-06 | 0.006182 |
| ENSRNOG00000021510 | *Tbc1d10c* | 23.81288 | -4.98867 | 1.058708 | -4.71204 | 2.45E-06 | 0.006182 |
| ENSRNOG00000028404 | *Ppp1r1b* | 6621.46 | -2.40862 | 0.515243 | -4.67473 | 2.94E-06 | 0.006595 |
| ENSRNOG00000029865 | *Prss56* | 25.78381 | -4.73849 | 1.044888 | -4.53493 | 5.76E-06 | 0.010861 |
| ENSRNOG00000042985 | *Slc9b1* | 57.64369 | 3.859984 | 0.852271 | 4.52906 | 5.92E-06 | 0.010861 |
| ENSRNOG00000003800 | *Rgs9* | 500.4486 | -3.9685 | 0.889867 | -4.45966 | 8.21E-06 | 0.013795 |
| **LPS vs. Control** | | | | | | | |
| ENSRNOG00000025848 | *Sspo* | 39.89744 | -5.25586 | 1.000946 | -5.25089 | 1.51E-07 | 0.003532 |
| ENSRNOG00000001302 | *Adora2a* | 368.615 | -4.19886 | 0.84191 | -4.9873 | 6.12E-07 | 0.005524 |
| ENSRNOG00000008428 | *Drd2* | 165.0874 | -4.7714 | 0.973289 | -4.90235 | 9.47E-07 | 0.005524 |
| ENSRNOG00000049580 | *Gpr6* | 101.2293 | -3.85405 | 0.785017 | -4.90951 | 9.13E-07 | 0.005524 |
| ENSRNOG00000025848 | *Sspo* | 39.89744 | -5.25586 | 1.000946 | -5.25089 | 1.51E-07 | 0.003532 |
| ENSRNOG00000001302 | *Adora2a* | 368.615 | -4.19886 | 0.84191 | -4.9873 | 6.12E-07 | 0.005524 |
| **EPA vs. Control** | | | | | | | |
| ENSRNOG00000049580 | *Gpr6* | 101.2293 | -4.29115 | 0.786959 | -5.45283 | 4.96E-08 | 0.001157 |
| ENSRNOG00000008428 | *Drd2* | 165.0874 | -4.77901 | 0.971983 | -4.91677 | 8.80E-07 | 0.010265 |
| ENSRNOG00000001302 | *Adora2a* | 368.615 | -4.05687 | 0.841085 | -4.82337 | 1.41E-06 | 0.010979 |
| **FLU+LPS vs. Control** | | | | | | | |
| ENSRNOG00000009269 | *Cga* | 21.59396 | -23.4346 | 4.450347 | -5.2658 | 1.40E-07 | 0.003257 |
| **FLU+EPA vs. Control** | | | | | | | |
| ENSRNOG00000003869 | *Sod3* | 1289.179632 | 1.22497965 | 0.189335943 | 6.469873754 | 9.81E-11 | 9.89E-07 |
| ENSRNOG00000031930 | *Bin2* | 702.7842329 | 1.253924824 | 0.194972207 | 6.431300348 | 1.27E-10 | 9.89E-07 |
| ENSRNOG00000060496 | *Kdm5d* | 1366.957623 | -0.779457661 | 0.130243355 | -5.984625189 | 2.17E-09 | 1.13E-05 |
| ENSRNOG00000014179 | *Rps2* | 5326.598905 | 1.169184793 | 0.199214637 | 5.868970338 | 4.39E-09 | 1.71E-05 |
| ENSRNOG00000009269 | *Cga* | 21.5939622 | -23.39775059 | 4.019788258 | -5.820642555 | 5.86E-09 | 1.83E-05 |
| ENSRNOG00000049931 | *LOC288913* | 1594.367668 | 0.981813711 | 0.175319642 | 5.600135258 | 2.14E-08 | 5.58E-05 |
| ENSRNOG00000034037 | *Zfp266* | 2388.078907 | -1.025608344 | 0.189051559 | -5.425019224 | 5.79E-08 | 0.000129465 |
| ENSRNOG00000005565 | *Traf3ip3* | 210.904955 | 1.484972861 | 0.279744828 | 5.308312126 | 1.11E-07 | 0.000208032 |
| ENSRNOG00000029773 | *Atm* | 895.2636029 | -1.044241265 | 0.197252792 | -5.293923866 | 1.20E-07 | 0.000208032 |
| ENSRNOG00000009521 |  | 191.6699903 | 1.506148411 | 0.286077518 | 5.264826188 | 1.40E-07 | 0.000212982 |
| ENSRNOG00000012722 | *Ppdpf* | 2035.27742 | 1.012727043 | 0.193408199 | 5.236215686 | 1.64E-07 | 0.000212982 |
| ENSRNOG00000022116 | *Gjb6* | 2522.749129 | 1.233709049 | 0.236254182 | 5.221956455 | 1.77E-07 | 0.000212982 |
| ENSRNOG00000023688 | *Drd1* | 174.6734668 | -3.143238298 | 0.60082627 | -5.231526074 | 1.68E-07 | 0.000212982 |
| ENSRNOG00000007324 | *Plxna2* | 3610.994136 | -1.22141323 | 0.235210252 | -5.192857102 | 2.07E-07 | 0.000231336 |
| ENSRNOG00000011054 | *Laptm5* | 1693.980184 | 1.010012674 | 0.195588861 | 5.163958061 | 2.42E-07 | 0.000252082 |
| ENSRNOG00000016190 | *Coq9* | 1905.96011 | 0.461071773 | 0.089782617 | 5.135423643 | 2.82E-07 | 0.000270069 |
| ENSRNOG00000021887 | *Cyb561d2* | 953.4422668 | 0.862387142 | 0.168187705 | 5.127527843 | 2.94E-07 | 0.000270069 |
| ENSRNOG00000008356 | *Myo5c* | 55.50094865 | -4.763524911 | 0.933130586 | -5.104885618 | 3.31E-07 | 0.000275116 |
| ENSRNOG00000017905 | *Map1lc3b* | 6064.726661 | 0.663205754 | 0.129962886 | 5.103039614 | 3.34E-07 | 0.000275116 |
| ENSRNOG00000014613 | *Ddah1* | 2529.238405 | 0.997880391 | 0.195987683 | 5.091546455 | 3.55E-07 | 0.000277713 |
| ENSRNOG00000008182 | *Htra3* | 463.5113397 | 1.267856218 | 0.249494459 | 5.081700909 | 3.74E-07 | 0.000278575 |
| ENSRNOG00000008190 | *Pnpla7* | 889.4383502 | 1.094323384 | 0.217804078 | 5.024347539 | 5.05E-07 | 0.000359091 |
| ENSRNOG00000001480 | *Ncf1* | 260.4873217 | 0.961436436 | 0.193800848 | 4.960950629 | 7.01E-07 | 0.00044109 |
| ENSRNOG00000002191 | *LOC498368* | 525.248805 | 1.061736465 | 0.214391477 | 4.952325899 | 7.33E-07 | 0.00044109 |
| ENSRNOG00000020029 | *Mcrip2* | 417.816656 | 1.147884332 | 0.231491369 | 4.958648507 | 7.10E-07 | 0.00044109 |
| ENSRNOG00000052025 | *Tle5* | 16621.24775 | 0.916226281 | 0.184742271 | 4.959483692 | 7.07E-07 | 0.00044109 |
| ENSRNOG00000019294 | *Stk16* | 2121.001712 | 0.700645349 | 0.141908228 | 4.937313067 | 7.92E-07 | 0.00045878 |
| ENSRNOG00000008428 | *Drd2* | 165.0873702 | -4.782097724 | 0.975386881 | -4.902770188 | 9.45E-07 | 0.000515281 |
| ENSRNOG00000016288 | *Tcea2* | 5491.71593 | 0.945915109 | 0.193172473 | 4.896738613 | 9.74E-07 | 0.000515281 |
| ENSRNOG00000038864 |  | 35.20345958 | 3.180745962 | 0.650717191 | 4.88806198 | 1.02E-06 | 0.000515281 |
| ENSRNOG00000059061 | *Uqcr10* | 5654.889784 | 1.153234241 | 0.235957321 | 4.887469631 | 1.02E-06 | 0.000515281 |
| ENSRNOG00000036829 | *Nckap1l* | 629.4801048 | 1.010949679 | 0.207815433 | 4.864651604 | 1.15E-06 | 0.000560358 |
| ENSRNOG00000049580 | *Gpr6* | 101.2292809 | -3.818474086 | 0.787126287 | -4.851158132 | 1.23E-06 | 0.000581688 |
| ENSRNOG00000015385 | *Pink1* | 11991.96809 | 0.866911326 | 0.178975909 | 4.843731929 | 1.27E-06 | 0.000586108 |
| ENSRNOG00000036834 | *Gpr84* | 279.5047689 | 1.348423467 | 0.279835268 | 4.81863304 | 1.45E-06 | 0.000630761 |
| ENSRNOG00000059834 | *Cib2* | 2031.535578 | 1.319609433 | 0.273906618 | 4.817734753 | 1.45E-06 | 0.000630761 |
| ENSRNOG00000031313 |  | 437.9514261 | 1.611639955 | 0.336002839 | 4.796506958 | 1.61E-06 | 0.000682437 |
| ENSRNOG00000020377 | *Cideb* | 38.76888861 | 2.637969077 | 0.55130766 | 4.784930933 | 1.71E-06 | 0.000685895 |
| ENSRNOG00000038239 | *Spata33* | 433.7908837 | 1.102807307 | 0.230464605 | 4.785148278 | 1.71E-06 | 0.000685895 |
| ENSRNOG00000055777 |  | 142.041954 | 1.441549655 | 0.301741221 | 4.777436945 | 1.78E-06 | 0.000694151 |
| ENSRNOG00000021528 | *Tusc2* | 4036.412721 | 0.680366836 | 0.143097376 | 4.75457242 | 1.99E-06 | 0.000758556 |
| ENSRNOG00000002963 | *C1ql1* | 762.9153952 | 1.273609068 | 0.2686891 | 4.740084616 | 2.14E-06 | 0.000778311 |
| ENSRNOG00000026974 | *Dbndd1* | 800.9696251 | 0.875270195 | 0.184666537 | 4.73973363 | 2.14E-06 | 0.000778311 |
| ENSRNOG00000046109 | *LOC689574* | 1115.829589 | 0.891796169 | 0.190114163 | 4.690845515 | 2.72E-06 | 0.000967053 |
| ENSRNOG00000002080 | *Urb1* | 711.9160142 | -0.90210442 | 0.193072863 | -4.67235222 | 2.98E-06 | 0.000986191 |
| ENSRNOG00000009982 | *Pnp* | 4426.572486 | 0.855546082 | 0.183068828 | 4.673357523 | 2.96E-06 | 0.000986191 |
| ENSRNOG00000013118 | *Atox1* | 1531.49806 | 1.108613727 | 0.237538567 | 4.667089384 | 3.05E-06 | 0.000986191 |
| ENSRNOG00000020918 | *Ccnd1* | 1091.851846 | 1.015790937 | 0.217079599 | 4.679347758 | 2.88E-06 | 0.000986191 |
| ENSRNOG00000039528 | *Fam32a* | 5303.959077 | 0.637195947 | 0.136598093 | 4.66474995 | 3.09E-06 | 0.000986191 |
| ENSRNOG00000005195 | *Cst3* | 86656.51604 | 1.055842754 | 0.226766489 | 4.656079302 | 3.22E-06 | 0.000988286 |
| ENSRNOG00000061088 | *Atf1* | 680.3092896 | -0.623668571 | 0.133885434 | -4.658225699 | 3.19E-06 | 0.000988286 |
| ENSRNOG00000005335 | *Galnt13* | 178.0265497 | -1.232025098 | 0.266178372 | -4.628569506 | 3.68E-06 | 0.001107362 |
| ENSRNOG00000001302 | *Adora2a* | 368.6149806 | -3.884631331 | 0.841805095 | -4.614644596 | 3.94E-06 | 0.00115202 |
| ENSRNOG00000033110 | *Svep1* | 494.3200413 | -2.098159479 | 0.454881769 | -4.612538077 | 3.98E-06 | 0.00115202 |
| ENSRNOG00000052968 | *Hnrnpa3* | 585.6208212 | -0.889752077 | 0.193176432 | -4.605903887 | 4.11E-06 | 0.001167744 |
| ENSRNOG00000013036 | *Epha8* | 832.0532305 | -2.410804737 | 0.524190641 | -4.599099159 | 4.24E-06 | 0.001184994 |
| ENSRNOG00000013572 | *Lxn* | 1599.994476 | 1.325034454 | 0.288524156 | 4.592455878 | 4.38E-06 | 0.00118756 |
| ENSRNOG00000021091 | *Trank1* | 2307.26779 | -2.319208528 | 0.505127684 | -4.591331259 | 4.40E-06 | 0.00118756 |
| ENSRNOG00000019178 | *Taf10* | 2251.979599 | 0.886070901 | 0.19328619 | 4.584243204 | 4.56E-06 | 0.001187609 |
| ENSRNOG00000031834 | *Nkain4* | 2029.287786 | 1.206383356 | 0.263102751 | 4.585217572 | 4.54E-06 | 0.001187609 |
| ENSRNOG00000011452 | *Aldoc* | 51967.3884 | 1.041673341 | 0.228164075 | 4.565457305 | 4.98E-06 | 0.0012778 |
| ENSRNOG00000019990 | *Rex1bd* | 1573.488188 | 1.116997079 | 0.245344084 | 4.552777722 | 5.29E-06 | 0.001335424 |
| ENSRNOG00000026679 | *Scn4b* | 1530.313028 | -2.488188365 | 0.547907466 | -4.541256539 | 5.59E-06 | 0.001388146 |
| ENSRNOG00000023541 | *Ahctf1* | 1546.638858 | -0.747920774 | 0.164832252 | -4.537466213 | 5.69E-06 | 0.001391239 |
| ENSRNOG00000006116 | *Hk2* | 531.5655883 | 1.007339659 | 0.222471675 | 4.527945673 | 5.96E-06 | 0.001433011 |
| ENSRNOG00000007769 | *Marchf2* | 4106.34316 | 0.823424197 | 0.182128222 | 4.521123581 | 6.15E-06 | 0.001457561 |
| ENSRNOG00000016249 | *Cep85* | 643.0309591 | -0.928915176 | 0.20578226 | -4.514068306 | 6.36E-06 | 0.001484438 |
| ENSRNOG00000008142 | *Brpf1* | 1484.889377 | -0.962922912 | 0.213610624 | -4.507841861 | 6.55E-06 | 0.001506016 |
| ENSRNOG00000051938 |  | 119.6422616 | 1.389181731 | 0.308380448 | 4.504765909 | 6.64E-06 | 0.001506016 |
| ENSRNOG00000016525 | *Susd3* | 202.3313588 | 1.378411276 | 0.306709293 | 4.494194693 | 6.98E-06 | 0.001560185 |
| ENSRNOG00000006019 | *G0s2* | 471.7713105 | 0.973071253 | 0.217010101 | 4.483990589 | 7.33E-06 | 0.001613678 |
| ENSRNOG00000011381 | *Acsbg1* | 10101.88162 | 0.876002142 | 0.195622194 | 4.478030443 | 7.53E-06 | 0.001625716 |
| ENSRNOG00000011719 | *Ngb* | 317.3334436 | 1.371274105 | 0.306328981 | 4.476475262 | 7.59E-06 | 0.001625716 |
| ENSRNOG00000025604 | *Atad2* | 146.3063049 | -1.363427888 | 0.304849497 | -4.472462325 | 7.73E-06 | 0.001634149 |
| ENSRNOG00000005668 | *Ndufa8* | 7570.905786 | 1.103889269 | 0.247110546 | 4.467188004 | 7.93E-06 | 0.001652614 |
| ENSRNOG00000003833 | *Nenf* | 1349.621523 | 1.364374039 | 0.305973744 | 4.459121302 | 8.23E-06 | 0.001693464 |
| ENSRNOG00000003237 | *Tp53bp2* | 5045.184551 | 1.136263351 | 0.255657968 | 4.444466799 | 8.81E-06 | 0.001754848 |
| ENSRNOG00000015213 | *Fxn* | 734.4111535 | 1.172284323 | 0.26386392 | 4.442760957 | 8.88E-06 | 0.001754848 |
| ENSRNOG00000017144 | *Mea1* | 3115.523049 | 0.854339604 | 0.192399 | 4.440457618 | 8.98E-06 | 0.001754848 |
| ENSRNOG00000022939 | *Gpkow* | 1439.979914 | -0.431491956 | 0.097046424 | -4.4462427 | 8.74E-06 | 0.001754848 |
| ENSRNOG00000001336 | *Orai1* | 492.6948751 | 0.847876364 | 0.191243515 | 4.433490805 | 9.27E-06 | 0.001763191 |
| ENSRNOG00000015936 | *LOC108349548* | 543.1007652 | 0.941306164 | 0.21229386 | 4.433977328 | 9.25E-06 | 0.001763191 |
| ENSRNOG00000015936 | *Gng5* | 543.1007652 | 0.941306164 | 0.21229386 | 4.433977328 | 9.25E-06 | 0.001763191 |
| ENSRNOG00000016134 | *Msh6* | 951.5396391 | -0.61748618 | 0.139380135 | -4.430230907 | 9.41E-06 | 0.001763191 |
| ENSRNOG00000021839 | *Cep126* | 487.5018947 | -1.083109549 | 0.244617302 | -4.427771616 | 9.52E-06 | 0.001763191 |
| ENSRNOG00000024207 | *Fgfrl1* | 664.7515808 | 0.682296581 | 0.154143566 | 4.426370819 | 9.58E-06 | 0.001763191 |
| ENSRNOG00000004048 | *Lrrk2* | 1192.823215 | -0.876717099 | 0.198780136 | -4.410486458 | 1.03E-05 | 0.001875564 |
| ENSRNOG00000012172 | *Spi1* | 260.8175557 | 1.232689649 | 0.280369785 | 4.396656545 | 1.10E-05 | 0.001976106 |
| ENSRNOG00000004554 | *Dcn* | 566.9095907 | 1.193028685 | 0.273068207 | 4.368976895 | 1.25E-05 | 0.002218427 |
| ENSRNOG00000033796 | *Kcnj5* | 154.6945616 | 1.319890185 | 0.302546884 | 4.362597191 | 1.29E-05 | 0.002258473 |
| ENSRNOG00000018250 | *Tnni3* | 38.26121498 | -4.195626293 | 0.963018562 | -4.356744987 | 1.32E-05 | 0.00229391 |
| ENSRNOG00000020005 | *Map2k2* | 3585.61689 | 0.664634556 | 0.152667924 | 4.353465599 | 1.34E-05 | 0.002302922 |
| ENSRNOG00000010389 | *Ndrg2* | 76694.11376 | 1.202793452 | 0.276436607 | 4.351064297 | 1.35E-05 | 0.002302984 |
| ENSRNOG00000002708 | *Phf8* | 893.4363505 | -0.974581688 | 0.224271496 | -4.345544151 | 1.39E-05 | 0.002336279 |
| ENSRNOG00000004459 | *Sdr9c7* | 338.9090222 | 1.441817334 | 0.332090214 | 4.341643546 | 1.41E-05 | 0.002352563 |
| ENSRNOG00000016062 | *Snta1* | 2476.263698 | 0.683499209 | 0.157741896 | 4.333022662 | 1.47E-05 | 0.002352563 |
| ENSRNOG00000016790 | *Kmt5b* | 1673.926963 | -0.614286474 | 0.141858618 | -4.330272511 | 1.49E-05 | 0.002352563 |
| ENSRNOG00000026236 | *Morc3* | 1278.469544 | -0.734036523 | 0.169297567 | -4.335777149 | 1.45E-05 | 0.002352563 |
| ENSRNOG00000042421 | *RGD1559909* | 3514.310239 | 1.067399213 | 0.246396348 | 4.332041528 | 1.48E-05 | 0.002352563 |
| ENSRNOG00000057806 | *Trpm7* | 818.5702 | -1.313481856 | 0.303259221 | -4.331218195 | 1.48E-05 | 0.002352563 |
| ENSRNOG00000005861 | *Hsd11b1* | 4730.853416 | 0.924498254 | 0.215086183 | 4.298268921 | 1.72E-05 | 0.002692054 |
| ENSRNOG00000010653 | *Elmsan1* | 891.2522241 | -0.853001455 | 0.198916095 | -4.288247533 | 1.80E-05 | 0.002788516 |
| ENSRNOG00000002896 | *Prdx6* | 13858.2719 | 0.971423191 | 0.227138401 | 4.276789774 | 1.90E-05 | 0.002907132 |
| ENSRNOG00000019702 | *Med29* | 780.4678686 | 0.678186918 | 0.15883817 | 4.269672188 | 1.96E-05 | 0.002972328 |
| ENSRNOG00000007023 | *Galm* | 1019.04786 | 0.776355387 | 0.182120415 | 4.262868532 | 2.02E-05 | 0.003022667 |
| ENSRNOG00000047928 |  | 572.1710924 | 1.161235243 | 0.272566621 | 4.260372158 | 2.04E-05 | 0.003022667 |
| ENSRNOG00000048172 | *Rac3* | 1105.582307 | 0.954966932 | 0.224196351 | 4.259511481 | 2.05E-05 | 0.003022667 |
| ENSRNOG00000007678 | *Zfp263* | 349.1421686 | -0.776799031 | 0.182643081 | -4.253098598 | 2.11E-05 | 0.003052974 |
| ENSRNOG00000010551 | *Lhx2* | 3257.286277 | 0.641697576 | 0.150855132 | 4.253733819 | 2.10E-05 | 0.003052974 |
| ENSRNOG00000058593 | *Zbtb1* | 278.3050181 | -1.293628665 | 0.304578678 | -4.247272572 | 2.16E-05 | 0.003104681 |
| ENSRNOG00000009536 | *Pgp* | 3038.889794 | 0.700033693 | 0.165034115 | 4.241751436 | 2.22E-05 | 0.003144433 |
| ENSRNOG00000023985 | *Fan1* | 312.9580249 | -1.067413759 | 0.251728288 | -4.240340912 | 2.23E-05 | 0.003144433 |
| ENSRNOG00000016186 | *Zfp709* | 378.56559 | -0.743164486 | 0.17560835 | -4.231942768 | 2.32E-05 | 0.003235058 |
| ENSRNOG00000024825 | *Fam163a* | 1671.529377 | 1.014845987 | 0.240550285 | 4.218851741 | 2.46E-05 | 0.003368554 |
| ENSRNOG00000054724 | *Pycr3* | 908.5696073 | 0.791484861 | 0.187528302 | 4.220615509 | 2.44E-05 | 0.003368554 |
| ENSRNOG00000043300 | *Enho* | 4439.854813 | 1.165236475 | 0.276615566 | 4.212476149 | 2.53E-05 | 0.003434955 |
| ENSRNOG00000009263 | *Ifi27* | 4759.652779 | 1.346537329 | 0.320437785 | 4.202180243 | 2.64E-05 | 0.003564027 |
| ENSRNOG00000006523 | *Atl2* | 1212.458173 | -0.613669609 | 0.146381572 | -4.192259994 | 2.76E-05 | 0.003691729 |
| ENSRNOG00000027869 | *Sox5* | 262.5493941 | 1.628252436 | 0.388718387 | 4.188771328 | 2.80E-05 | 0.003717163 |
| ENSRNOG00000057352 | *LOC100909912* | 503.7215969 | 1.01338948 | 0.242107201 | 4.185705656 | 2.84E-05 | 0.003736033 |
| ENSRNOG00000011984 | *Cxcl14* | 5542.069781 | 0.95809454 | 0.229002417 | 4.183774799 | 2.87E-05 | 0.003736524 |
| ENSRNOG00000004206 | *Glrx5* | 2088.911693 | 0.74259635 | 0.178322838 | 4.16433677 | 3.12E-05 | 0.004035882 |
| ENSRNOG00000016660 | *Cox5b* | 7919.573588 | 0.957251085 | 0.230148271 | 4.15927994 | 3.19E-05 | 0.00409245 |
| ENSRNOG00000006749 | *Tmtc3* | 511.9650278 | -0.792497492 | 0.190973215 | -4.149783479 | 3.33E-05 | 0.004163601 |
| ENSRNOG00000009889 | *Pgm1* | 3218.939066 | 0.556024027 | 0.133924059 | 4.151785942 | 3.30E-05 | 0.004163601 |
| ENSRNOG00000047988 | *Bola2* | 3088.639565 | 1.234031373 | 0.297327878 | 4.150405885 | 3.32E-05 | 0.004163601 |
| ENSRNOG00000012550 | *Uqcrh* | 7258.410297 | 0.884202844 | 0.213258741 | 4.146150537 | 3.38E-05 | 0.004196609 |
| ENSRNOG00000020178 | *Cope* | 2999.736915 | 0.680287487 | 0.164242281 | 4.14197539 | 3.44E-05 | 0.004240106 |
| ENSRNOG00000016545 | *Ift140* | 1042.482961 | -1.032990211 | 0.249810525 | -4.13509483 | 3.55E-05 | 0.004335034 |
| ENSRNOG00000000595 | *Traf3ip2* | 65.86300538 | 1.286313436 | 0.311289853 | 4.132204829 | 3.59E-05 | 0.004355886 |
| ENSRNOG00000020202 | *Asrgl1* | 7584.936328 | 0.751577245 | 0.182046585 | 4.128488572 | 3.65E-05 | 0.004392822 |
| ENSRNOG00000043094 | *Oxct1* | 10583.53817 | 0.720831437 | 0.174742882 | 4.125097566 | 3.71E-05 | 0.004424018 |
| ENSRNOG00000014319 | *Smchd1* | 1129.908943 | -0.818988372 | 0.198797265 | -4.119716494 | 3.79E-05 | 0.004427224 |
| ENSRNOG00000029805 | *Zfp780b* | 216.0477146 | -1.199552306 | 0.290991072 | -4.122299351 | 3.75E-05 | 0.004427224 |
| ENSRNOG00000054809 |  | 74.75496388 | -3.214989263 | 0.780173534 | -4.120864297 | 3.77E-05 | 0.004427224 |
| ENSRNOG00000012329 | *Saraf* | 10271.36995 | 0.606513205 | 0.14731989 | 4.116981113 | 3.84E-05 | 0.004430521 |
| ENSRNOG00000019974 | *Uba52* | 14268.01034 | 1.181508217 | 0.287043476 | 4.116129834 | 3.85E-05 | 0.004430521 |
| ENSRNOG00000018092 | *Cd83* | 1043.858054 | 1.002840282 | 0.243855554 | 4.112435687 | 3.92E-05 | 0.004469176 |
| ENSRNOG00000001348 | *Erp29* | 2628.482766 | 0.762151836 | 0.185441979 | 4.109920746 | 3.96E-05 | 0.004485389 |
| ENSRNOG00000010557 | *Smarcd2* | 1205.070742 | -0.663095707 | 0.161895738 | -4.095819416 | 4.21E-05 | 0.004733079 |
| ENSRNOG00000024577 | *Gamt* | 2029.739421 | 0.909850602 | 0.222234235 | 4.094106385 | 4.24E-05 | 0.004734145 |
| ENSRNOG00000004538 | *Brd1* | 1752.04271 | -0.598322609 | 0.146329396 | -4.088875003 | 4.33E-05 | 0.004785293 |
| ENSRNOG00000021177 | *Cox8a* | 12014.39738 | 0.875925677 | 0.214250472 | 4.088325545 | 4.34E-05 | 0.004785293 |
| ENSRNOG00000027049 | *Atp5mf* | 8750.874751 | 1.053446658 | 0.25785302 | 4.085454023 | 4.40E-05 | 0.004810984 |
| ENSRNOG00000050490 | *Mir3064* | 69.98049847 | -2.089657544 | 0.511875686 | -4.082353585 | 4.46E-05 | 0.00484178 |
| ENSRNOG00000024312 | *Xkr8* | 397.1490985 | -1.144683188 | 0.280652334 | -4.078651946 | 4.53E-05 | 0.004852117 |
| ENSRNOG00000042740 | *Mrpl42* | 1511.214211 | 1.015611748 | 0.248922622 | 4.080029932 | 4.50E-05 | 0.004852117 |
| ENSRNOG00000008455 | *Tmem87a* | 495.2699448 | -0.988396194 | 0.242529019 | -4.075372913 | 4.59E-05 | 0.004854501 |
| ENSRNOG00000038761 | *Hacd2* | 597.6878086 | 0.641883043 | 0.157460126 | 4.076479919 | 4.57E-05 | 0.004854501 |
| ENSRNOG00000059715 | *Cc2d2a* | 982.262528 | -1.251387616 | 0.307354135 | -4.071484567 | 4.67E-05 | 0.004903137 |
| ENSRNOG00000004637 | *Fbxo7* | 2199.671841 | 0.436647611 | 0.107575906 | 4.058972188 | 4.93E-05 | 0.004946309 |
| ENSRNOG00000006500 | *Snf8* | 4636.769991 | 1.013726608 | 0.249270947 | 4.066765974 | 4.77E-05 | 0.004946309 |
| ENSRNOG00000014352 | *Smim12* | 810.4569273 | 0.668047081 | 0.164498598 | 4.061111094 | 4.88E-05 | 0.004946309 |
| ENSRNOG00000016267 | *Chst15* | 382.5637677 | -2.321202623 | 0.572113689 | -4.057240138 | 4.97E-05 | 0.004946309 |
| ENSRNOG00000022822 | *Fam169b* | 135.9214138 | 1.135416541 | 0.279707601 | 4.059298119 | 4.92E-05 | 0.004946309 |
| ENSRNOG00000023020 | *Fdx2* | 2696.192291 | 1.059899609 | 0.261213869 | 4.05759316 | 4.96E-05 | 0.004946309 |
| ENSRNOG00000034015 | *Capn2* | 4284.631879 | 0.623589583 | 0.153642313 | 4.058709937 | 4.93E-05 | 0.004946309 |
| ENSRNOG00000054866 |  | 80.99396354 | -2.223836914 | 0.548016384 | -4.057975234 | 4.95E-05 | 0.004946309 |
| ENSRNOG00000004196 | *LOC108352650* | 2666.193505 | 1.202668892 | 0.296874979 | 4.0510955 | 5.10E-05 | 0.005045893 |
| ENSRNOG00000004196 | *Rps29* | 2666.193505 | 1.202668892 | 0.296874979 | 4.0510955 | 5.10E-05 | 0.005045893 |
| ENSRNOG00000013681 | *Kcns1* | 255.9372009 | 1.680238505 | 0.414955709 | 4.049199631 | 5.14E-05 | 0.005054947 |
| ENSRNOG00000020464 | *Mrpl54* | 2370.553354 | 1.127773217 | 0.278933091 | 4.043167535 | 5.27E-05 | 0.005059544 |
| ENSRNOG00000029465 | *Slc26a10* | 89.96782729 | -1.724943348 | 0.42637347 | -4.045616041 | 5.22E-05 | 0.005059544 |
| ENSRNOG00000037402 | *Cops9* | 4206.393424 | 0.950787763 | 0.235078669 | 4.044551419 | 5.24E-05 | 0.005059544 |
| ENSRNOG00000049110 | *Dpm2* | 1202.255105 | 0.764704983 | 0.189078809 | 4.0443717 | 5.25E-05 | 0.005059544 |
| ENSRNOG00000014530 | *Nav2* | 841.2574316 | -0.931737616 | 0.230604599 | -4.040412122 | 5.34E-05 | 0.005088144 |
| ENSRNOG00000033261 | *Fam107a* | 36993.78153 | 1.051890387 | 0.260954423 | 4.030935265 | 5.56E-05 | 0.005233916 |
| ENSRNOG00000050016 | *Gtf3a* | 688.4597108 | 0.78139884 | 0.193840695 | 4.031139287 | 5.55E-05 | 0.005233916 |
| ENSRNOG00000005840 | *Cacfd1* | 500.7613665 | 0.604409485 | 0.149997801 | 4.02945565 | 5.59E-05 | 0.005235428 |
| ENSRNOG00000047102 | *Popdc3* | 67.48912969 | 3.459492275 | 0.858996682 | 4.027363955 | 5.64E-05 | 0.005250765 |
| ENSRNOG00000007830 | *Apold1* | 176.3329612 | -1.68570501 | 0.418929372 | -4.023840581 | 5.73E-05 | 0.005271909 |
| ENSRNOG00000019022 | *Fam89a* | 250.8755067 | 1.200154152 | 0.298276195 | 4.023633703 | 5.73E-05 | 0.005271909 |
| ENSRNOG00000011535 | *Gcsh* | 3037.234623 | 0.930246143 | 0.231396408 | 4.020140808 | 5.82E-05 | 0.005319397 |
| ENSRNOG00000007302 | *Fbn1* | 780.284739 | -1.385611998 | 0.345393737 | -4.011688255 | 6.03E-05 | 0.005481476 |
| ENSRNOG00000025848 | *Sspo* | 39.89744408 | -3.88488709 | 0.969525761 | -4.006997281 | 6.15E-05 | 0.005559129 |
| ENSRNOG00000021232 | *Ddrgk1* | 4348.216482 | 0.687429809 | 0.171654238 | 4.004735434 | 6.21E-05 | 0.00558033 |
| ENSRNOG00000010065 | *Dgkh* | 183.6862716 | -2.103771759 | 0.525545892 | -4.003021988 | 6.25E-05 | 0.005588797 |
| ENSRNOG00000033498 | *Cib1* | 741.0182144 | 1.067286879 | 0.266991192 | 3.997461009 | 6.40E-05 | 0.005689178 |
| ENSRNOG00000003510 | *Fmo2* | 41.24472203 | -1.938272225 | 0.485639954 | -3.991171259 | 6.57E-05 | 0.00577657 |
| ENSRNOG00000053047 | *Top2a* | 88.20685539 | -2.762991508 | 0.692080555 | -3.992297555 | 6.54E-05 | 0.00577657 |
| ENSRNOG00000021355 | *Ca6* | 1098.439534 | 1.051743079 | 0.26377308 | 3.987302571 | 6.68E-05 | 0.005838744 |
| ENSRNOG00000059894 | *Hmmr* | 100.1195488 | -1.113248083 | 0.279616677 | -3.98133651 | 6.85E-05 | 0.00592112 |
| ENSRNOG00000061845 | *Cttnbp2* | 5814.421824 | -0.892200576 | 0.224094265 | -3.981362818 | 6.85E-05 | 0.00592112 |
| ENSRNOG00000024114 | *Zdbf2* | 428.0677919 | -1.311719103 | 0.329587888 | -3.979876537 | 6.90E-05 | 0.005924867 |
| ENSRNOG00000005814 | *NEWGENE_1582771* | 167.4757011 | -1.074951631 | 0.27090881 | -3.967946382 | 7.25E-05 | 0.006128344 |
| ENSRNOG00000005814 | *Katnbl1* | 167.4757011 | -1.074951631 | 0.27090881 | -3.967946382 | 7.25E-05 | 0.006128344 |
| ENSRNOG00000008530 | *Stoml1* | 2034.251416 | 0.52346798 | 0.131915846 | 3.968196359 | 7.24E-05 | 0.006128344 |
| ENSRNOG00000038980 | *Lypd6* | 861.4353157 | 1.032719103 | 0.260126966 | 3.970057854 | 7.19E-05 | 0.006128344 |
| ENSRNOG00000007100 | *Ccdc136* | 1963.319471 | 0.62468057 | 0.157546179 | 3.965063283 | 7.34E-05 | 0.006136549 |
| ENSRNOG00000020845 | *Tyrobp* | 521.2546326 | 1.185979159 | 0.299077246 | 3.965461019 | 7.33E-05 | 0.006136549 |
| ENSRNOG00000046996 | *Pea15* | 34253.6194 | 0.616290061 | 0.15557601 | 3.961343795 | 7.45E-05 | 0.006199793 |
| ENSRNOG00000017071 | *Babam1* | 3275.295627 | 0.579155492 | 0.146285923 | 3.959065094 | 7.52E-05 | 0.00622612 |
| ENSRNOG00000037898 | *LOC100912028* | 12.10227794 | -3.609051789 | 0.912356992 | -3.955745196 | 7.63E-05 | 0.006280001 |
| ENSRNOG00000002922 | *Adora2b* | 374.3201029 | 0.872794911 | 0.22088189 | 3.951410006 | 7.77E-05 | 0.006328263 |
| ENSRNOG00000049105 |  | 2720.836218 | 0.872763652 | 0.220836468 | 3.952081185 | 7.75E-05 | 0.006328263 |
| ENSRNOG00000009837 | *Tmem131l* | 397.2997072 | -1.147519818 | 0.290655702 | -3.948038215 | 7.88E-05 | 0.006384782 |
| ENSRNOG00000008099 | *Galnt12* | 23.39107241 | -2.266386137 | 0.574591498 | -3.944343318 | 8.00E-05 | 0.0064506 |
| ENSRNOG00000012274 | *Ddi2* | 2484.715065 | -1.013996546 | 0.257242475 | -3.941792834 | 8.09E-05 | 0.00648616 |
| ENSRNOG00000014169 |  | 36.37954479 | 1.961080452 | 0.499111687 | 3.929141519 | 8.52E-05 | 0.006600096 |
| ENSRNOG00000016828 | *Cmtm5* | 2575.57298 | 0.886945934 | 0.225452148 | 3.934076223 | 8.35E-05 | 0.006600096 |
| ENSRNOG00000019570 | *Gng3* | 9033.596173 | 0.974508548 | 0.247724287 | 3.933843378 | 8.36E-05 | 0.006600096 |
| ENSRNOG00000021903 | *Atad5* | 159.0269055 | -1.11602345 | 0.28370715 | -3.933716341 | 8.36E-05 | 0.006600096 |
| ENSRNOG00000052004 | *Txnl4a* | 1314.933669 | 1.04689858 | 0.266314129 | 3.931066615 | 8.46E-05 | 0.006600096 |
| ENSRNOG00000055451 |  | 98.7115238 | -3.085349701 | 0.785104602 | -3.929858126 | 8.50E-05 | 0.006600096 |
| ENSRNOG00000058663 |  | 679.5570156 | -1.060376828 | 0.269725974 | -3.931311515 | 8.45E-05 | 0.006600096 |
| ENSRNOG00000010775 | *Arrdc4* | 331.470954 | -0.815687372 | 0.207926 | -3.922969571 | 8.75E-05 | 0.006720931 |
| ENSRNOG00000023340 | *Tmem72* | 15.70225043 | -5.256085414 | 1.340016424 | -3.922403725 | 8.77E-05 | 0.006720931 |
| ENSRNOG00000018564 | *Nup93* | 442.0259405 | -0.886487232 | 0.226100413 | -3.920767857 | 8.83E-05 | 0.006733716 |
| ENSRNOG00000008553 | *Mthfr* | 139.7131588 | -1.522626987 | 0.388713766 | -3.917090463 | 8.96E-05 | 0.006771167 |
| ENSRNOG00000023130 | *Lyplal1* | 554.0732262 | 0.737592988 | 0.188248794 | 3.918181742 | 8.92E-05 | 0.006771167 |
| ENSRNOG00000026636 | *Urm1* | 1610.529454 | 0.617860558 | 0.157818561 | 3.915005653 | 9.04E-05 | 0.006797107 |
| ENSRNOG00000046002 | *Micos13* | 4657.24682 | 0.921937037 | 0.235617947 | 3.912847248 | 9.12E-05 | 0.006825356 |
| ENSRNOG00000019120 | *Hmgcs2* | 514.2386585 | 1.597071403 | 0.408403041 | 3.910527692 | 9.21E-05 | 0.006858424 |
| ENSRNOG00000014625 | *Atp5f1d* | 8559.925227 | 0.84355011 | 0.21584439 | 3.908140078 | 9.30E-05 | 0.006861197 |
| ENSRNOG00000048174 | *Uqcrq* | 4658.100678 | 0.945558743 | 0.241938479 | 3.908261088 | 9.30E-05 | 0.006861197 |
| ENSRNOG00000011879 | *Nfat5* | 2787.119665 | -0.735804028 | 0.188499762 | -3.903474574 | 9.48E-05 | 0.006897273 |
| ENSRNOG00000013282 | *Mctp1* | 1395.672867 | -1.140188283 | 0.291965791 | -3.905211904 | 9.41E-05 | 0.006897273 |
| ENSRNOG00000019077 | *Lipa* | 1971.412106 | 0.780168793 | 0.199822601 | 3.904307061 | 9.45E-05 | 0.006897273 |
| ENSRNOG00000057194 | *Tars2* | 1686.509175 | 0.473339301 | 0.121365401 | 3.90011729 | 9.61E-05 | 0.006961245 |
| ENSRNOG00000001982 | *Cblb* | 960.7186985 | -0.816172709 | 0.20951661 | -3.895503596 | 9.80E-05 | 0.007033765 |
| ENSRNOG00000029594 |  | 1555.863507 | 1.005226406 | 0.258056443 | 3.895374189 | 9.80E-05 | 0.007033765 |
| ENSRNOG00000009888 | *Timm8b* | 38.82806078 | 1.623794714 | 0.417375862 | 3.890485437 | 0.000100044 | 0.00714423 |
| ENSRNOG00000007069 | *Adhfe1* | 2353.829605 | 0.910344752 | 0.234187877 | 3.887241149 | 0.00010139 | 0.007201717 |
| ENSRNOG00000016692 | *Hsdl2* | 1999.068712 | 0.637699298 | 0.164087664 | 3.886332972 | 0.00010177 | 0.007201717 |
| ENSRNOG00000006559 | *Akap8* | 1236.07643 | -0.783638641 | 0.201799959 | -3.883244807 | 0.000103072 | 0.00724434 |
| ENSRNOG00000019670 | *Wdr3* | 767.6086412 | -0.577241019 | 0.148669613 | -3.882710175 | 0.000103299 | 0.00724434 |
| ENSRNOG00000001514 | *Cdca7* | 146.8718 | -2.605981717 | 0.671725122 | -3.879535887 | 0.000104656 | 0.007306761 |
| ENSRNOG00000024239 | *Fam89b* | 2271.406939 | 0.750975545 | 0.193704468 | 3.876913906 | 0.00010579 | 0.007314856 |
| ENSRNOG00000028883 |  | 189.8238161 | 1.147185572 | 0.295890604 | 3.877059819 | 0.000105726 | 0.007314856 |
| ENSRNOG00000054891 | *Cnot6* | 999.2240457 | -0.746251246 | 0.192529847 | -3.876028866 | 0.000106175 | 0.007314856 |
| ENSRNOG00000024056 | *Zfp17* | 471.4060115 | -0.71318222 | 0.184197198 | -3.871840753 | 0.000108017 | 0.007388931 |
| ENSRNOG00000028841 | *Mt1m* | 1351.425359 | 1.237258238 | 0.319586229 | 3.871437896 | 0.000108195 | 0.007388931 |
| ENSRNOG00000007955 | *Timp4* | 1125.520654 | 1.412540289 | 0.365011595 | 3.869850465 | 0.000108902 | 0.007404871 |
| ENSRNOG00000000917 | *Nupr2* | 220.2836069 | 1.047971474 | 0.271300708 | 3.862767195 | 0.00011211 | 0.007524832 |
| ENSRNOG00000007764 | *Frmd4b* | 755.3775348 | -0.90431346 | 0.23409735 | -3.86298033 | 0.000112012 | 0.007524832 |
| ENSRNOG00000029938 | *Pik3c2b* | 720.4820727 | -1.217943389 | 0.315260078 | -3.863297237 | 0.000111867 | 0.007524832 |
| ENSRNOG00000008921 | *Dynll2* | 22065.83538 | 0.523579139 | 0.135718443 | 3.857833387 | 0.000114397 | 0.007548726 |
| ENSRNOG00000014284 | *Elof1* | 1434.764462 | 0.766161504 | 0.198442746 | 3.860869287 | 0.000112984 | 0.007548726 |
| ENSRNOG00000020533 | *Htra1* | 7532.949919 | 0.791499453 | 0.205096694 | 3.859152655 | 0.000113781 | 0.007548726 |
| ENSRNOG00000030364 |  | 710.1350257 | 1.261998679 | 0.327108956 | 3.858037689 | 0.000114301 | 0.007548726 |
| ENSRNOG00000047796 | *Pcbd2* | 215.1767969 | 1.393484685 | 0.361421701 | 3.855564515 | 0.000115463 | 0.007587075 |
| ENSRNOG00000061450 | *Homer2* | 426.8051457 | 0.904808268 | 0.234855957 | 3.852609399 | 0.000116866 | 0.007647127 |
| ENSRNOG00000002928 | *Guk1* | 12325.14614 | 0.879840932 | 0.228668954 | 3.847662375 | 0.00011925 | 0.0077384 |
| ENSRNOG00000031273 |  | 2611.231054 | 0.995691324 | 0.258773609 | 3.84773134 | 0.000119217 | 0.0077384 |
| ENSRNOG00000018550 | *Ncbp3* | 710.8704964 | -0.61014832 | 0.158960274 | -3.838369825 | 0.000123854 | 0.008003927 |
| ENSRNOG00000000990 | *Pdap1* | 6332.527572 | 0.732615316 | 0.19094938 | 3.836699101 | 0.000124699 | 0.008025388 |
| ENSRNOG00000061883 | *Aqp9* | 495.020711 | 1.29373904 | 0.337464756 | 3.833701197 | 0.000126229 | 0.008090584 |
| ENSRNOG00000014482 | *Slf2* | 1194.126303 | -1.032975951 | 0.269635836 | -3.831003948 | 0.000127621 | 0.008146416 |
| ENSRNOG00000003120 | *Prelp* | 1747.124867 | 0.789615461 | 0.206209631 | 3.829188084 | 0.000128567 | 0.008173394 |
| ENSRNOG00000012260 | *Ddx25* | 2358.946942 | 0.697732013 | 0.182404745 | 3.825185639 | 0.000130674 | 0.008207246 |
| ENSRNOG00000021153 | *Fkbp2* | 4900.172818 | 1.100514546 | 0.287622381 | 3.82624795 | 0.000130111 | 0.008207246 |
| ENSRNOG00000030680 | *Ddx5* | 16476.89435 | -0.653117142 | 0.170725964 | -3.825529085 | 0.000130492 | 0.008207246 |
| ENSRNOG00000019629 | *Lamp1* | 18430.72242 | 0.512332898 | 0.133986589 | 3.823762531 | 0.000131431 | 0.008221767 |
| ENSRNOG00000008176 | *Nppa* | 435.6213362 | 2.071047968 | 0.541791757 | 3.8225904 | 0.000132057 | 0.008228048 |
| ENSRNOG00000060436 | *Vti1b* | 2884.66644 | 0.635959485 | 0.166572458 | 3.817914989 | 0.000134584 | 0.008352236 |
| ENSRNOG00000037673 |  | 3319.661587 | 1.42917787 | 0.374480916 | 3.816423771 | 0.0001354 | 0.008369639 |
| ENSRNOG00000015158 | *Pikfyve* | 2771.176639 | -0.577107259 | 0.151296945 | -3.814401263 | 0.000136514 | 0.008392157 |
| ENSRNOG00000015980 | *Dclre1c* | 163.0877164 | -1.150718239 | 0.30172359 | -3.813815948 | 0.000136837 | 0.008392157 |
| ENSRNOG00000012099 | *Tent2* | 536.1729191 | -0.536779835 | 0.140918192 | -3.80915925 | 0.00013944 | 0.008518376 |
| ENSRNOG00000000583 | *Cdk19* | 3694.561929 | -0.688222343 | 0.180858619 | -3.805305751 | 0.000141629 | 0.008612682 |
| ENSRNOG00000043098 | *Mt2A* | 642.6043846 | 1.340991584 | 0.352474211 | 3.804509778 | 0.000142085 | 0.008612682 |
| ENSRNOG00000019624 | *Morc2* | 1244.032839 | -0.521978763 | 0.137286445 | -3.802114359 | 0.000143466 | 0.008662827 |
| ENSRNOG00000012988 | *Lix1* | 10986.06525 | 0.911142767 | 0.239780132 | 3.799909352 | 0.000144749 | 0.008706654 |
| ENSRNOG00000018568 | *Rab5c* | 7333.827519 | 0.498442875 | 0.13128805 | 3.79655936 | 0.000146718 | 0.00879129 |
| ENSRNOG00000000701 | *Iscu* | 3970.669261 | 0.710153066 | 0.187098502 | 3.795610639 | 0.00014728 | 0.008791296 |
| ENSRNOG00000001170 | *Cox6a1* | 16592.37475 | 0.88675168 | 0.233895374 | 3.79123223 | 0.000149902 | 0.008846463 |
| ENSRNOG00000017895 | *Eno1* | 9186.523555 | 0.837958529 | 0.220988366 | 3.791867163 | 0.000149519 | 0.008846463 |
| ENSRNOG00000018239 | *Dhrs4* | 381.1395345 | 1.086268139 | 0.286453689 | 3.792124808 | 0.000149364 | 0.008846463 |
| ENSRNOG00000016456 | *Il33* | 2210.478521 | 0.786980304 | 0.207778781 | 3.787587453 | 0.000152117 | 0.008943459 |
| ENSRNOG00000001136 | *Pebp1* | 21842.69466 | 1.008843535 | 0.266730981 | 3.782251055 | 0.000155416 | 0.009035534 |
| ENSRNOG00000004697 | *Baalc* | 6934.919827 | 0.622071587 | 0.164418049 | 3.783475049 | 0.000154654 | 0.009035534 |
| ENSRNOG00000019811 | *Timm23* | 873.5609962 | 0.633913021 | 0.167600576 | 3.782284263 | 0.000155396 | 0.009035534 |
| ENSRNOG00000019811 | *LOC100362432* | 873.5609962 | 0.633913021 | 0.167600576 | 3.782284263 | 0.000155396 | 0.009035534 |
| ENSRNOG00000015869 | *Pccb* | 1254.432103 | 0.569605753 | 0.150781907 | 3.777679723 | 0.000158296 | 0.009168872 |
| ENSRNOG00000058094 |  | 871.0644247 | -2.317372011 | 0.613876993 | -3.774977782 | 0.000160022 | 0.009234623 |
| ENSRNOG00000016497 | *Bloc1s5* | 1504.719043 | 0.838391809 | 0.222247752 | 3.772329765 | 0.00016173 | 0.009298898 |
| ENSRNOG00000018170 | *Slc27a1* | 3803.430814 | 0.679129994 | 0.180137993 | 3.77005419 | 0.000163212 | 0.009346888 |
| ENSRNOG00000019682 | *Timm13* | 3198.178358 | 0.937121349 | 0.248654865 | 3.768763382 | 0.000164058 | 0.009346888 |
| ENSRNOG00000045698 | *Lin7c* | 632.5968572 | -1.405727038 | 0.373039337 | -3.76830779 | 0.000164358 | 0.009346888 |
| ENSRNOG00000005618 | *Fmc1* | 489.7853533 | 0.956329906 | 0.253852397 | 3.767267584 | 0.000165044 | 0.009351899 |
| ENSRNOG00000057558 |  | 533.8515052 | 1.013255306 | 0.269086536 | 3.765536999 | 0.000166192 | 0.009382921 |
| ENSRNOG00000026407 | *Fam184a* | 340.68985 | -0.956883272 | 0.254261278 | -3.763385755 | 0.000167628 | 0.009429998 |
| ENSRNOG00000018569 | *Ahcyl1* | 16453.06854 | 0.691865704 | 0.184085573 | 3.758391781 | 0.000171009 | 0.009541256 |
| ENSRNOG00000021255 | *Smox* | 1741.657368 | 0.737868094 | 0.196358123 | 3.757767093 | 0.000171436 | 0.009541256 |
| ENSRNOG00000026488 | *Gltpd2* | 108.0878658 | 1.776633046 | 0.47262214 | 3.759098221 | 0.000170527 | 0.009541256 |
| ENSRNOG00000014202 | *Snx20* | 114.3342602 | 1.109933982 | 0.295447283 | 3.756791981 | 0.000172105 | 0.009544528 |
| ENSRNOG00000000568 | *Slc29a3* | 1392.089711 | 0.948338343 | 0.252662388 | 3.753381536 | 0.000174465 | 0.009641195 |
| ENSRNOG00000056608 |  | 36.03740417 | 2.124235858 | 0.566234501 | 3.751512589 | 0.000175771 | 0.00967916 |
| ENSRNOG00000009656 | *Rspo1* | 273.7771304 | 3.391198565 | 0.904594659 | 3.748859816 | 0.00017764 | 0.00974778 |
| ENSRNOG00000001211 | *Gatd3a* | 3650.891196 | 0.575870201 | 0.153650532 | 3.747921956 | 0.000178306 | 0.009750081 |
| ENSRNOG00000015329 | *Kpna2* | 232.3467072 | -0.920951793 | 0.246027209 | -3.743292443 | 0.000181625 | 0.009896961 |
| ENSRNOG00000001431 | *Rasa4* | 144.9507535 | 1.114283501 | 0.297779843 | 3.741970877 | 0.000182583 | 0.009908644 |
| ENSRNOG00000046799 | *Phb* | 2500.295952 | 0.561998961 | 0.150216838 | 3.741251426 | 0.000183106 | 0.009908644 |
| ENSRNOG00000016977 | *Calb2* | 532.4148012 | 1.053786636 | 0.281969026 | 3.737242534 | 0.000186049 | 0.009953404 |
| ENSRNOG00000027791 |  | 6501.738952 | 1.156721341 | 0.309520437 | 3.737140439 | 0.000186125 | 0.009953404 |
| ENSRNOG00000028238 | *Sh3bgr* | 457.5398204 | 1.261851575 | 0.337694845 | 3.736662243 | 0.000186479 | 0.009953404 |
| ENSRNOG00000034161 |  | 5063.173301 | 0.816678367 | 0.21852061 | 3.737305913 | 0.000186003 | 0.009953404 |
| ENSRNOG00000050200 | *Kdm3b* | 1707.571974 | -0.503930947 | 0.134897014 | -3.735671618 | 0.000187215 | 0.009958684 |
| ENSRNOG00000039668 | *Col8a1* | 13.74005859 | -5.559107485 | 1.488580298 | -3.734502931 | 0.000188086 | 0.009971124 |
| ENSRNOG00000039417 | *Dda1* | 3023.884767 | 0.557799927 | 0.149589402 | 3.728873288 | 0.000192338 | 0.010162065 |
| ENSRNOG00000007325 | *Usp28* | 825.1676261 | -0.576802218 | 0.154789104 | -3.726374824 | 0.000194253 | 0.010228721 |
| ENSRNOG00000037446 | *Pxmp2* | 579.1943635 | 1.055569258 | 0.283439745 | 3.724139877 | 0.000195982 | 0.010285122 |
| ENSRNOG00000007679 | *Cyth4* | 726.5896835 | 0.81753519 | 0.220190562 | 3.712853005 | 0.000204936 | 0.010712184 |
| ENSRNOG00000045884 |  | 535.5674793 | -1.829868046 | 0.492937556 | -3.712170081 | 0.00020549 | 0.010712184 |
| ENSRNOG00000056069 | *Kif11* | 43.65193945 | -2.252529084 | 0.607506362 | -3.707827974 | 0.000209045 | 0.01086129 |
| ENSRNOG00000007887 | *Elk4* | 1106.437784 | -0.621717041 | 0.167794719 | -3.705224133 | 0.000211204 | 0.010866017 |
| ENSRNOG00000012266 | *Zcchc17* | 2699.585395 | 0.614665178 | 0.165892361 | 3.705204833 | 0.00021122 | 0.010866017 |
| ENSRNOG00000046883 | *Mydgf* | 1053.056038 | 0.446143046 | 0.120358455 | 3.706786075 | 0.000209906 | 0.010866017 |
| ENSRNOG00000019692 | *Metrn* | 4162.285084 | 1.145323103 | 0.309240341 | 3.703666536 | 0.000212506 | 0.01089631 |
| ENSRNOG00000001065 | *Cyth3* | 1632.714221 | -0.78980076 | 0.213588679 | -3.697765083 | 0.000217506 | 0.011043101 |
| ENSRNOG00000003209 | *Pcp4l1* | 1180.634992 | -2.108445891 | 0.570092204 | -3.698429616 | 0.000216937 | 0.011043101 |
| ENSRNOG00000011568 | *Rspo3* | 907.4417829 | -1.066177665 | 0.288392727 | -3.69696447 | 0.000218193 | 0.011043101 |
| ENSRNOG00000017571 | *Ndufa2* | 3154.54222 | 1.050554017 | 0.284008316 | 3.699025551 | 0.000216429 | 0.011043101 |
| ENSRNOG00000010038 | *Psmc5* | 10279.0384 | 0.797116691 | 0.215751238 | 3.69461005 | 0.000220224 | 0.011109971 |
| ENSRNOG00000018958 | *Mt3* | 27665.93835 | 1.033753419 | 0.279903048 | 3.693255312 | 0.000221402 | 0.011133436 |
| ENSRNOG00000014890 | *Mrpl43* | 1906.60951 | 0.686189192 | 0.185883854 | 3.691494325 | 0.00022294 | 0.01113447 |
| ENSRNOG00000019270 | *P2ry6* | 187.9284638 | 1.081213061 | 0.293013103 | 3.689981945 | 0.00022427 | 0.01113447 |
| ENSRNOG00000020083 | *Scly* | 375.6761093 | 0.722950468 | 0.195903348 | 3.690342593 | 0.000223952 | 0.01113447 |
| ENSRNOG00000026091 | *Slc10a4* | 34.08047577 | -2.317386242 | 0.627628232 | -3.692291269 | 0.000222243 | 0.01113447 |
| ENSRNOG00000013169 | *Traf4* | 564.387569 | 0.759454678 | 0.205990359 | 3.686845742 | 0.000227051 | 0.011202717 |
| ENSRNOG00000020526 | *Mau2* | 2621.095305 | -0.693865811 | 0.188201901 | -3.686816174 | 0.000227077 | 0.011202717 |
| ENSRNOG00000047853 | *Ss18l2* | 1407.760142 | 0.726427719 | 0.197088233 | 3.685799551 | 0.000227986 | 0.011212167 |
| ENSRNOG00000007338 | *Fbln2* | 690.011873 | 1.190165336 | 0.32322307 | 3.682179424 | 0.000231249 | 0.011231355 |
| ENSRNOG00000037567 | *RGD1309730* | 1084.865628 | 0.970784907 | 0.263569147 | 3.683226647 | 0.0002303 | 0.011231355 |
| ENSRNOG00000058522 | *Fam214a* | 1080.958848 | -0.754166341 | 0.20481085 | -3.682257761 | 0.000231177 | 0.011231355 |
| ENSRNOG00000061881 |  | 12.4954085 | -5.225000941 | 1.41839159 | -3.68375065 | 0.000229827 | 0.011231355 |
| ENSRNOG00000011491 | *Dnajc13* | 1140.816641 | -0.971304427 | 0.264005725 | -3.679103653 | 0.000234055 | 0.011293199 |
| ENSRNOG00000023107 | *Yeats2* | 957.9620006 | -0.819981806 | 0.222830955 | -3.679837955 | 0.000233382 | 0.011293199 |
| ENSRNOG00000060293 | *Ndufa3* | 1745.00001 | 0.864251052 | 0.234952051 | 3.678414585 | 0.000234688 | 0.011293199 |
| ENSRNOG00000032765 |  | 440.0324799 | 0.950500108 | 0.258515673 | 3.676760092 | 0.000236215 | 0.011331799 |
| ENSRNOG00000001177 | *Acads* | 994.5361863 | 0.744642481 | 0.20265356 | 3.674460398 | 0.000238353 | 0.011364623 |
| ENSRNOG00000020308 | *Ech1* | 2312.864251 | 0.520730455 | 0.141691213 | 3.675107595 | 0.000237749 | 0.011364623 |
| ENSRNOG00000015865 | *Ap2s1* | 6447.367527 | 0.800645734 | 0.217951611 | 3.67350225 | 0.000239249 | 0.01137267 |
| ENSRNOG00000010184 | *Brk1* | 10452.39973 | 0.71712899 | 0.195265253 | 3.672588844 | 0.000240106 | 0.011378824 |
| ENSRNOG00000017241 | *Dpcd* | 4451.846119 | 0.833871726 | 0.227138376 | 3.671205816 | 0.000241409 | 0.011406021 |
| ENSRNOG00000057578 | *Prodh2* | 61.99851442 | 1.99364118 | 0.543353125 | 3.669144591 | 0.000243363 | 0.011463736 |
| ENSRNOG00000028166 | *Asmtl* | 1515.285939 | 1.061572799 | 0.289497122 | 3.666954589 | 0.000245456 | 0.011527604 |
| ENSRNOG00000023492 | *Greb1l* | 58.13940815 | -1.810005519 | 0.494126117 | -3.663043616 | 0.000249236 | 0.011670069 |
| ENSRNOG00000015334 | *Fcho2* | 2615.697692 | 0.710460367 | 0.194237006 | 3.657698292 | 0.00025449 | 0.011810014 |
| ENSRNOG00000024651 | *Greb1* | 72.26991506 | -1.977332 | 0.54039852 | -3.659025564 | 0.000253176 | 0.011810014 |
| ENSRNOG00000045743 | *Etnppl* | 1053.640168 | 1.242793394 | 0.339712984 | 3.658362945 | 0.000253831 | 0.011810014 |
| ENSRNOG00000043498 | *Sik2* | 1626.632451 | -0.732847248 | 0.200481708 | -3.655431985 | 0.000256749 | 0.011879591 |
| ENSRNOG00000009409 | *Fbxo2* | 4792.668195 | 0.777297182 | 0.212767428 | 3.653271506 | 0.00025892 | 0.011944699 |
| ENSRNOG00000042233 | *Rpl41* | 14055.44164 | 1.196626927 | 0.327765078 | 3.65086767 | 0.000261356 | 0.012021602 |
| ENSRNOG00000014999 | *Tnpo1* | 2059.568723 | -0.464962601 | 0.127457148 | -3.647991558 | 0.000264298 | 0.012121295 |
| ENSRNOG00000009431 | *Tbc1d4* | 189.2582868 | -1.576607685 | 0.432334499 | -3.646731154 | 0.000265598 | 0.012145264 |
| ENSRNOG00000016313 | *Ctnnbip1* | 1370.091413 | 0.892567715 | 0.245149658 | 3.64090949 | 0.000271677 | 0.012361343 |
| ENSRNOG00000047230 | *Zfp551* | 409.808068 | -0.882181095 | 0.242311217 | -3.640694424 | 0.000271904 | 0.012361343 |
| ENSRNOG00000018053 | *Fech* | 1567.8895 | 0.451705308 | 0.124168079 | 3.637853706 | 0.000274919 | 0.012367331 |
| ENSRNOG00000018186 | *Lamtor5* | 1912.413057 | 0.748191024 | 0.205667313 | 3.637870375 | 0.000274902 | 0.012367331 |
| ENSRNOG00000032575 | *Ocel1* | 983.7460504 | 0.890317341 | 0.244691578 | 3.63852875 | 0.0002742 | 0.012367331 |
| ENSRNOG00000033024 |  | 1322.330832 | 1.052102388 | 0.289230433 | 3.637592274 | 0.000275199 | 0.012367331 |
| ENSRNOG00000000632 | *Cdk1* | 25.14269894 | -3.05031185 | 0.838843234 | -3.63633123 | 0.000276549 | 0.012392392 |
| ENSRNOG00000014793 | *Gpr149* | 96.41522016 | -1.863272796 | 0.512903108 | -3.632796853 | 0.000280366 | 0.012456363 |
| ENSRNOG00000020970 | *Rundc3a* | 14683.81182 | 0.545931835 | 0.15021812 | 3.634260869 | 0.000278779 | 0.012456363 |
| ENSRNOG00000060365 |  | 9.446888096 | 3.798869252 | 1.045691156 | 3.632878818 | 0.000280277 | 0.012456363 |
| ENSRNOG00000003738 | *Ush2a* | 62.43995164 | -1.907786831 | 0.525353117 | -3.631437161 | 0.000281847 | 0.012486713 |
| ENSRNOG00000005281 | *Stx16* | 2672.919187 | -0.561258401 | 0.154722138 | -3.62752485 | 0.000286151 | 0.01264158 |
| ENSRNOG00000001295 | *S100b* | 86806.03911 | 0.926963322 | 0.255758519 | 3.624369284 | 0.000289667 | 0.012703564 |
| ENSRNOG00000001314 | *Fam20c* | 1992.020485 | 0.532964905 | 0.147072916 | 3.623814095 | 0.00029029 | 0.012703564 |
| ENSRNOG00000017976 | *Slco2b1* | 1354.852968 | 0.844747959 | 0.233139559 | 3.623357455 | 0.000290803 | 0.012703564 |
| ENSRNOG00000018494 | *Ppp1r3c* | 3550.796419 | 0.7338878 | 0.202430055 | 3.62538952 | 0.000288526 | 0.012703564 |
| ENSRNOG00000009267 | *B3gnt2* | 644.7037277 | -1.048790439 | 0.289672098 | -3.620612572 | 0.000293906 | 0.012767781 |
| ENSRNOG00000014048 | *Cyld* | 1293.571946 | -0.980810206 | 0.270856435 | -3.621144194 | 0.000293303 | 0.012767781 |
| ENSRNOG00000016855 | *B3galnt2* | 524.638661 | -0.784297087 | 0.216768522 | -3.618131818 | 0.000296737 | 0.012840253 |
| ENSRNOG00000022044 | *Cabp4* | 82.25444211 | -1.917815055 | 0.530210831 | -3.61708012 | 0.000297945 | 0.012840253 |
| ENSRNOG00000030319 | *Morn2* | 93.06807046 | 1.145988023 | 0.316833879 | 3.616999634 | 0.000298038 | 0.012840253 |
| ENSRNOG00000017097 | *Lhpp* | 2415.235351 | 0.785602496 | 0.217417503 | 3.613336026 | 0.000302283 | 0.012953283 |
| ENSRNOG00000017575 | *Gins2* | 1713.476391 | -0.676716322 | 0.187284543 | -3.613305782 | 0.000302318 | 0.012953283 |
| ENSRNOG00000005713 | *Ccdc82* | 934.4932948 | -0.984937661 | 0.27268151 | -3.612044174 | 0.000303793 | 0.012980917 |
| ENSRNOG00000015304 | *Tmem160* | 2136.97014 | 0.888286412 | 0.246073632 | 3.609839894 | 0.000306386 | 0.013020575 |
| ENSRNOG00000027891 | *Dhrs11* | 233.2387635 | 1.149303465 | 0.318369711 | 3.609964845 | 0.000306239 | 0.013020575 |
| ENSRNOG00000013231 | *Ptafr* | 454.5690637 | 0.926020553 | 0.256649304 | 3.608116361 | 0.000308428 | 0.013071838 |
| ENSRNOG00000013774 | *Lmnb1* | 148.6436913 | -1.342743584 | 0.372298023 | -3.606636356 | 0.000310192 | 0.013111057 |
| ENSRNOG00000055222 |  | 1611.142138 | 0.762915009 | 0.211633127 | 3.604894094 | 0.00031228 | 0.013163749 |
| ENSRNOG00000043225 | *Zfp771* | 1754.854449 | 0.882041656 | 0.244826533 | 3.602720856 | 0.000314904 | 0.013238651 |
| ENSRNOG00000001391 | *Sdsl* | 417.5419772 | 1.037409533 | 0.288171264 | 3.599975655 | 0.000318247 | 0.013343336 |
| ENSRNOG00000001827 | *Masp1* | 1479.837648 | 0.780464033 | 0.217119984 | 3.594620908 | 0.000324864 | 0.013402972 |
| ENSRNOG00000014874 | *Zfyve28* | 999.5562483 | -0.698075223 | 0.194132785 | -3.595864669 | 0.000323316 | 0.013402972 |
| ENSRNOG00000017397 | *Upf3a* | 2907.17547 | 0.857845343 | 0.238735173 | 3.593292659 | 0.000326525 | 0.013402972 |
| ENSRNOG00000018481 | *Zpr1* | 1807.496142 | 0.440853901 | 0.122597187 | 3.595954463 | 0.000323204 | 0.013402972 |
| ENSRNOG00000042283 | *Zfp267* | 410.9503777 | -0.589468829 | 0.163851299 | -3.597584115 | 0.000321187 | 0.013402972 |
| ENSRNOG00000043141 | *Ap3s2* | 2155.501604 | 0.594694227 | 0.16533362 | 3.596934646 | 0.000321989 | 0.013402972 |
| ENSRNOG00000045779 |  | 676.6356408 | -1.122572301 | 0.31225024 | -3.595104689 | 0.000324261 | 0.013402972 |
| ENSRNOG00000048320 | *Ndufa11* | 4073.725665 | 0.919703025 | 0.255920212 | 3.593710001 | 0.000326003 | 0.013402972 |
| ENSRNOG00000057159 | *Rsl1* | 74.1247242 | -1.447568437 | 0.402971982 | -3.592230974 | 0.000327859 | 0.013422481 |
| ENSRNOG00000020602 | *Ndufa13* | 3190.169564 | 0.911969853 | 0.253974115 | 3.590798431 | 0.000329667 | 0.01346124 |
| ENSRNOG00000021068 | *Pip5k1a* | 3349.971324 | -0.607058003 | 0.169116991 | -3.589574285 | 0.000331218 | 0.013489388 |
| ENSRNOG00000028137 | *Mki67* | 145.7158606 | -1.676129188 | 0.467085843 | -3.588482106 | 0.000332609 | 0.013510829 |
| ENSRNOG00000001092 | *Kl* | 43.43836091 | -4.323581705 | 1.205981013 | -3.585115899 | 0.000336929 | 0.013582488 |
| ENSRNOG00000002270 | *Hnrnpdl* | 4440.841857 | -0.601104622 | 0.167723349 | -3.583905425 | 0.000338495 | 0.013582488 |
| ENSRNOG00000014857 | *RGD1560065* | 1666.38499 | 0.58266632 | 0.162586319 | 3.583735243 | 0.000338715 | 0.013582488 |
| ENSRNOG00000015554 | *Ankdd1a* | 161.2166148 | -1.498748746 | 0.418192874 | -3.583869642 | 0.000338541 | 0.013582488 |
| ENSRNOG00000016272 | *Edf1* | 6574.464796 | 0.879552176 | 0.245404378 | 3.584093252 | 0.000338251 | 0.013582488 |
| ENSRNOG00000053210 | *Zc3h11a* | 1608.522727 | -1.049579143 | 0.292979959 | -3.582426413 | 0.000340418 | 0.01361583 |
| ENSRNOG00000038205 | *LOC108349548* | 260.6157199 | 1.035495451 | 0.289149843 | 3.581172454 | 0.000342056 | 0.013646455 |
| ENSRNOG00000052062 | *Tut4* | 488.9941668 | -0.828936401 | 0.231630733 | -3.578697833 | 0.00034531 | 0.013741246 |
| ENSRNOG00000007350 | *Rac2* | 164.0967443 | 1.077894223 | 0.30148528 | 3.575279776 | 0.000349854 | 0.013851544 |
| ENSRNOG00000024533 | *Eogt* | 668.5788405 | -0.817945038 | 0.228744341 | -3.575804473 | 0.000349153 | 0.013851544 |
| ENSRNOG00000057241 |  | 40.8970968 | -1.309306628 | 0.366309831 | -3.574314738 | 0.000351146 | 0.01386762 |
| ENSRNOG00000014272 | *Rpl35* | 13055.63185 | 1.141948072 | 0.319617585 | 3.572857455 | 0.000353107 | 0.013909925 |
| ENSRNOG00000037632 | *LOC688553* | 28.1854028 | -2.817245351 | 0.789448132 | -3.568626283 | 0.000358858 | 0.014100951 |
| ENSRNOG00000005686 | *Suclg2* | 3356.464863 | 0.824922197 | 0.231438823 | 3.564320736 | 0.0003648 | 0.014193152 |
| ENSRNOG00000007398 | *Zfp691* | 286.1265179 | 0.632156852 | 0.17735814 | 3.56429567 | 0.000364835 | 0.014193152 |
| ENSRNOG00000021156 | *Vegfb* | 1287.60445 | 1.030999184 | 0.289105655 | 3.566167476 | 0.00036224 | 0.014193152 |
| ENSRNOG00000061231 | *Selenom* | 7488.596213 | 1.040418211 | 0.2918111 | 3.565382567 | 0.000363326 | 0.014193152 |
| ENSRNOG00000005877 | *Lrpprc* | 3607.770491 | -0.499850731 | 0.140308611 | -3.562509295 | 0.000367327 | 0.014200367 |
| ENSRNOG00000009206 | *Fezf2* | 856.1277088 | 0.792652607 | 0.222517001 | 3.562211441 | 0.000367744 | 0.014200367 |
| ENSRNOG00000025787 | *Spag6* | 187.319372 | -1.310783379 | 0.367930777 | -3.562581499 | 0.000367226 | 0.014200367 |
| ENSRNOG00000001173 | *Cabp1* | 4219.01378 | 0.926735617 | 0.26056964 | 3.556575582 | 0.00037572 | 0.014273279 |
| ENSRNOG00000001714 | *Atp13a4* | 634.4897782 | 1.075345966 | 0.302434051 | 3.555637874 | 0.000377063 | 0.014273279 |
| ENSRNOG00000005868 | *Ttc21b* | 261.1105751 | -0.957276518 | 0.268931437 | -3.559556027 | 0.000371482 | 0.014273279 |
| ENSRNOG00000007784 | *Bloc1s1* | 2105.175664 | 0.896511132 | 0.252176603 | 3.555092424 | 0.000377846 | 0.014273279 |
| ENSRNOG00000008193 | *Cr1l* | 649.569387 | -0.62769716 | 0.176420438 | -3.55796169 | 0.000373744 | 0.014273279 |
| ENSRNOG00000017401 | *Tmco4* | 202.7899788 | 0.772050497 | 0.216965499 | 3.558402139 | 0.000373118 | 0.014273279 |
| ENSRNOG00000028087 | *Sdhaf4* | 1000.120639 | 0.7396881 | 0.207789494 | 3.559795477 | 0.000371144 | 0.014273279 |
| ENSRNOG00000051487 | *Kremen1* | 329.2758504 | -1.674955897 | 0.470914418 | -3.55681592 | 0.000375377 | 0.014273279 |
| ENSRNOG00000058588 | *Nfxl1* | 485.2123762 | -0.920154257 | 0.258803135 | -3.555421609 | 0.000377373 | 0.014273279 |
| ENSRNOG00000057996 | *Phactr3* | 9015.757099 | 0.55735809 | 0.15683767 | 3.553725893 | 0.000379815 | 0.014313071 |
| ENSRNOG00000049326 | *Tmem234* | 258.5155446 | 0.73831283 | 0.20779692 | 3.553049921 | 0.000380792 | 0.014315407 |
| ENSRNOG00000021817 | *Irak2* | 360.5366801 | -0.645262859 | 0.181684617 | -3.551554727 | 0.000382962 | 0.014362465 |
| ENSRNOG00000057699 |  | 11.94108428 | -2.836469885 | 0.799427838 | -3.548124983 | 0.000387984 | 0.014515988 |
| ENSRNOG00000013436 | *Pde7b* | 182.5725937 | -1.71127612 | 0.482654765 | -3.545548998 | 0.000391796 | 0.014623627 |
| ENSRNOG00000011422 |  | 1060.821541 | -0.552016347 | 0.155785587 | -3.543436584 | 0.000394948 | 0.014698052 |
| ENSRNOG00000024027 | *Lemd3* | 694.4772704 | -0.611898741 | 0.17270857 | -3.542955283 | 0.00039567 | 0.014698052 |
| ENSRNOG00000033663 | *P4ha2* | 789.0546277 | -0.727969843 | 0.205547652 | -3.541611082 | 0.000397691 | 0.014738141 |
| ENSRNOG00000022943 | *Dgka* | 1558.85406 | -0.733314294 | 0.207342542 | -3.536728585 | 0.000405116 | 0.01497779 |
| ENSRNOG00000018477 | *Otud4* | 414.9963278 | -1.634587481 | 0.462268488 | -3.536013208 | 0.000406214 | 0.014982988 |
| ENSRNOG00000028404 | *Ppp1r1b* | 6621.460229 | -1.821260486 | 0.515244652 | -3.534748938 | 0.000408163 | 0.015019429 |
| ENSRNOG00000015501 | *Ddhd2* | 1097.662058 | -1.043434033 | 0.295246344 | -3.534113303 | 0.000409146 | 0.015020255 |
| ENSRNOG00000017803 | *Apbb1ip* | 586.3677474 | 0.871188996 | 0.246637312 | 3.53226764 | 0.000412012 | 0.015033814 |
| ENSRNOG00000038228 | *Serp2* | 2936.550647 | 0.813533494 | 0.230330972 | 3.532019544 | 0.000412399 | 0.015033814 |
| ENSRNOG00000048237 | *Tcta* | 2239.514255 | 0.591228936 | 0.167334852 | 3.533208596 | 0.000410548 | 0.015033814 |
| ENSRNOG00000018198 | *Dapk1* | 3017.618279 | -0.942726243 | 0.266983295 | -3.531030821 | 0.000413944 | 0.015055029 |
| ENSRNOG00000007407 | *Ndufa12* | 403.2485161 | 0.736767423 | 0.208915042 | 3.526636543 | 0.000420874 | 0.015271577 |
| ENSRNOG00000061424 | *Map3k3* | 1013.453198 | -0.752341828 | 0.213562678 | -3.522815108 | 0.000426989 | 0.015457599 |
| ENSRNOG00000046700 | *Mettl27* | 148.7966945 | 0.824234115 | 0.234123279 | 3.520513293 | 0.000430712 | 0.015556377 |
| ENSRNOG00000018795 | *Rpl18a* | 10354.62826 | 0.954826253 | 0.271320712 | 3.51917937 | 0.000432884 | 0.015598783 |
| ENSRNOG00000020557 | *Ryr1* | 457.3863924 | -1.983952157 | 0.563945721 | -3.517984235 | 0.000434838 | 0.015633183 |
| ENSRNOG00000007369 | *Shisa4* | 4778.255902 | 0.566624809 | 0.161213906 | 3.514739032 | 0.000440186 | 0.015778024 |
| ENSRNOG00000007387 | *Per1* | 1208.9068 | -0.841248074 | 0.239377337 | -3.514317965 | 0.000440885 | 0.015778024 |
| ENSRNOG00000029726 | *Gstm1* | 8182.711631 | 0.727929427 | 0.207307653 | 3.511348542 | 0.000445839 | 0.015909805 |
| ENSRNOG00000051952 | *Tes* | 719.246745 | 0.796809445 | 0.226953405 | 3.510894422 | 0.000446602 | 0.015909805 |
| ENSRNOG00000062253 |  | 227.3818872 | 1.068589602 | 0.304531685 | 3.508960325 | 0.000449862 | 0.015989527 |
| ENSRNOG00000050315 | *Dcxr* | 329.2637101 | 0.942416132 | 0.268628165 | 3.508255112 | 0.000451056 | 0.015995622 |
| ENSRNOG00000013578 | *Trem2* | 883.1348563 | 1.037180386 | 0.295715241 | 3.507361952 | 0.000452573 | 0.016013101 |
| ENSRNOG00000021179 | *Naa40* | 354.4021287 | -0.983430754 | 0.280529471 | -3.505623677 | 0.000455539 | 0.016081651 |
| ENSRNOG00000004351 | *Slc25a29* | 789.8664224 | 0.450822739 | 0.1286762 | 3.503544081 | 0.000459111 | 0.016171242 |
| ENSRNOG00000034071 | *Chmp4bl1* | 4765.545391 | 0.599326368 | 0.171189259 | 3.500957771 | 0.000463589 | 0.016292299 |
| ENSRNOG00000014064 | *Ctsh* | 2428.168809 | 0.745267451 | 0.213125046 | 3.49685532 | 0.000470777 | 0.016507809 |
| ENSRNOG00000004937 | *Csmd3* | 685.8306389 | -0.710532703 | 0.203257654 | -3.495724223 | 0.000472777 | 0.016540851 |
| ENSRNOG00000018932 | *Ccdc124* | 3704.862445 | 0.785251109 | 0.224744259 | 3.493976269 | 0.000475883 | 0.016612364 |
| **FLU vs. CUS** | | | | | | | |
| ENSRNOG00000003738 | *Ush2a* | 62.43995 | 2.928242 | 0.450383 | 6.50167 | 7.94E-11 | 1.24E-06 |
| ENSRNOG00000006146 | *Trim54* | 73.29952 | -4.00397 | 0.735814 | -5.44155 | 5.28E-08 | 0.00038 |
| ENSRNOG00000008736 | *Slamf8* | 86.6299 | -1.37561 | 0.255497 | -5.38408 | 7.28E-08 | 0.00038 |
| ENSRNOG00000006885 | *Dbx2* | 164.5865 | -1.56549 | 0.305427 | -5.12559 | 2.97E-07 | 0.000994 |
| ENSRNOG00000018680 | *Rpl17* | 3337.816 | -2.16724 | 0.423911 | -5.11249 | 3.18E-07 | 0.000994 |
| ENSRNOG00000015554 | *Ankdd1a* | 161.2166 | 1.915356 | 0.383291 | 4.997129 | 5.82E-07 | 0.001399 |
| ENSRNOG00000060414 |  | 45.78751 | -3.97205 | 0.797124 | -4.98297 | 6.26E-07 | 0.001399 |
| ENSRNOG00000007030 | *Epha7* | 3329.05 | 1.479424 | 0.30228 | 4.894224 | 9.87E-07 | 0.001883 |
| ENSRNOG00000013036 | *Epha8* | 832.0532 | 2.288839 | 0.469424 | 4.875849 | 1.08E-06 | 0.001883 |
| ENSRNOG00000049959 | *Igsf21* | 563.1432 | -1.37445 | 0.285455 | -4.81495 | 1.47E-06 | 0.002303 |
| ENSRNOG00000027230 | *Fhod3* | 481.2656 | -2.17653 | 0.455715 | -4.77608 | 1.79E-06 | 0.002541 |
| ENSRNOG00000002434 | *Tmem100* | 897.3488 | -0.82241 | 0.173522 | -4.73954 | 2.14E-06 | 0.002792 |
| ENSRNOG00000002361 | *Prkg2* | 238.9475 | -2.49573 | 0.532386 | -4.68783 | 2.76E-06 | 0.003228 |
| ENSRNOG00000015896 | *Rbpms2* | 199.1909 | -1.27062 | 0.272412 | -4.66433 | 3.1E-06 | 0.003228 |
| ENSRNOG00000029330 | *Ca5b* | 210.1655 | -1.07825 | 0.231129 | -4.66516 | 3.08E-06 | 0.003228 |
| ENSRNOG00000001259 | *Cux2* | 547.7272 | -1.96759 | 0.426506 | -4.61327 | 3.96E-06 | 0.003647 |
| ENSRNOG00000048187 | *Epop* | 351.898 | -2.14223 | 0.463709 | -4.61977 | 3.84E-06 | 0.003647 |
| ENSRNOG00000051612 |  | 38.68564 | -3.12099 | 0.68 | -4.58969 | 4.44E-06 | 0.003857 |
| ENSRNOG00000026087 | *Igfn1* | 47.056 | -1.66467 | 0.366125 | -4.54671 | 5.45E-06 | 0.004485 |
| ENSRNOG00000007118 | *Eva1a* | 459.7839 | -0.98346 | 0.220463 | -4.4609 | 8.16E-06 | 0.006078 |
| ENSRNOG00000026974 | *Dbndd1* | 800.9696 | -0.72697 | 0.162787 | -4.46574 | 7.98E-06 | 0.006078 |
| ENSRNOG00000001258 | *Snx8* | 621.7581 | -0.95311 | 0.216506 | -4.40224 | 1.07E-05 | 0.007616 |
| ENSRNOG00000005809 | *Arhgdib* | 525.6785 | -0.82186 | 0.188126 | -4.36864 | 1.25E-05 | 0.008501 |
| ENSRNOG00000011589 | *Camk2d* | 1173.796 | -1.56586 | 0.361983 | -4.32577 | 1.52E-05 | 0.009143 |
| ENSRNOG00000013169 | *Traf4* | 564.3876 | -0.79401 | 0.183333 | -4.33099 | 1.48E-05 | 0.009143 |
| ENSRNOG00000048651 | *Nrtn* | 54.85551 | -1.76144 | 0.406315 | -4.33517 | 1.46E-05 | 0.009143 |
| ENSRNOG00000012795 | *Cables1* | 1011.54 | -1.1977 | 0.27846 | -4.30117 | 1.7E-05 | 0.009841 |
| ENSRNOG00000000704 | *Cmklr1* | 160.2616 | -1.03151 | 0.241926 | -4.26372 | 2.01E-05 | 0.010143 |
| ENSRNOG00000004018 | *Tdrd5* | 99.51492 | 1.806681 | 0.423462 | 4.26645 | 1.99E-05 | 0.010143 |
| ENSRNOG00000010065 | *Dgkh* | 183.6863 | 1.998079 | 0.466649 | 4.281758 | 1.85E-05 | 0.010143 |
| ENSRNOG00000016366 | *Colec12* | 293.0481 | -1.68053 | 0.393269 | -4.27323 | 1.93E-05 | 0.010143 |
| ENSRNOG00000001658 | *Kcnj6* | 829.3245 | 1.538986 | 0.36186 | 4.252991 | 2.11E-05 | 0.010309 |
| ENSRNOG00000004171 | *Dnah9* | 936.2654 | 1.410073 | 0.335498 | 4.202928 | 2.63E-05 | 0.01212 |
| ENSRNOG00000005286 | *Coch* | 412.9342 | -3.62343 | 0.861187 | -4.20748 | 2.58E-05 | 0.01212 |
| ENSRNOG00000000605 | *Hs3st5* | 32.91022 | -2.70036 | 0.644327 | -4.19098 | 2.78E-05 | 0.012411 |
| ENSRNOG00000006019 | *G0s2* | 471.7713 | -0.80978 | 0.193557 | -4.18368 | 2.87E-05 | 0.01246 |
| ENSRNOG00000053288 | *Ank3* | 7861.873 | 0.541 | 0.129739 | 4.169902 | 3.05E-05 | 0.01288 |
| ENSRNOG00000018808 | *Vip* | 666.4469 | -1.3893 | 0.334509 | -4.15324 | 3.28E-05 | 0.013491 |
| ENSRNOG00000017444 | *Nrsn1* | 10757.61 | -1.33492 | 0.322152 | -4.14376 | 3.42E-05 | 0.013701 |
| ENSRNOG00000007302 | *Fbn1* | 780.2847 | 1.281235 | 0.309655 | 4.137623 | 3.51E-05 | 0.01372 |
| ENSRNOG00000028082 | *Tal2* | 16.48052 | 2.773875 | 0.671752 | 4.129316 | 3.64E-05 | 0.013878 |
| ENSRNOG00000001476 | *Cldn4* | 16.11613 | -5.61642 | 1.365529 | -4.113 | 3.91E-05 | 0.013881 |
| ENSRNOG00000046984 | *St6galnac6* | 6670.08 | -0.66159 | 0.160784 | -4.11477 | 3.88E-05 | 0.013881 |
| ENSRNOG00000058646 | *Zfp36l1* | 1188.739 | -0.86766 | 0.210521 | -4.12147 | 3.76E-05 | 0.013881 |
| ENSRNOG00000037196 | *Spag17* | 32.44612 | 2.882231 | 0.702001 | 4.105736 | 4.03E-05 | 0.014007 |
| ENSRNOG00000003005 | *Rasgef1c* | 136.1529 | -3.55617 | 0.868986 | -4.09232 | 4.27E-05 | 0.01452 |
| ENSRNOG00000000036 | *Klhdc8a* | 162.6838 | -2.6208 | 0.644617 | -4.06567 | 4.79E-05 | 0.015937 |
| ENSRNOG00000014010 | *Gfra2* | 215.3203 | -2.21706 | 0.548149 | -4.04463 | 5.24E-05 | 0.017074 |
| ENSRNOG00000006645 | *Ryr3* | 1045.524 | 1.230368 | 0.304817 | 4.03642 | 5.43E-05 | 0.017322 |
| ENSRNOG00000001528 | *Gap43* | 11009.53 | -0.94444 | 0.236127 | -3.99971 | 6.34E-05 | 0.017457 |
| ENSRNOG00000001963 | *Mx2* | 890.7056 | -1.76273 | 0.438833 | -4.01687 | 5.9E-05 | 0.017457 |
| ENSRNOG00000004026 | *Atp2b1* | 14683.62 | 0.844457 | 0.20962 | 4.028505 | 5.61E-05 | 0.017457 |
| ENSRNOG00000008323 | *Pitpnm3* | 1179.428 | -1.88505 | 0.469707 | -4.01324 | 5.99E-05 | 0.017457 |
| ENSRNOG00000008465 | *Tmem176b* | 3258.007 | -0.74389 | 0.185309 | -4.0143 | 5.96E-05 | 0.017457 |
| ENSRNOG00000013668 | *Capg* | 207.2334 | -1.26465 | 0.316247 | -3.99894 | 6.36E-05 | 0.017457 |
| ENSRNOG00000014936 | *Ifitm2* | 176.3765 | -1.82512 | 0.456117 | -4.00144 | 6.3E-05 | 0.017457 |
| ENSRNOG00000057044 | *Garnl3* | 828.4141 | -2.0368 | 0.509049 | -4.00118 | 6.3E-05 | 0.017457 |
| ENSRNOG00000006850 | *Ovol2* | 23.81928 | -3.61052 | 0.905663 | -3.9866 | 6.7E-05 | 0.017508 |
| ENSRNOG00000011826 | *Lzts1* | 2804.429 | -1.46791 | 0.368259 | -3.98609 | 6.72E-05 | 0.017508 |
| ENSRNOG00000016346 | *Prkcd* | 371.8734 | -1.17979 | 0.295435 | -3.99341 | 6.51E-05 | 0.017508 |
| ENSRNOG00000001316 | *Anapc5* | 8341.837 | -0.3809 | 0.096314 | -3.95477 | 7.66E-05 | 0.018442 |
| ENSRNOG00000005359 | *Csrnp3* | 145.1902 | 0.942069 | 0.237791 | 3.961743 | 7.44E-05 | 0.018442 |
| ENSRNOG00000016587 | *Ninj1* | 756.9352 | -0.90247 | 0.227352 | -3.96948 | 7.2E-05 | 0.018442 |
| ENSRNOG00000016831 | *Serpinh1* | 1868.544 | -0.71465 | 0.180762 | -3.95352 | 7.7E-05 | 0.018442 |
| ENSRNOG00000030763 | *Dpp4* | 34.12239 | -3.82852 | 0.966874 | -3.95968 | 7.5E-05 | 0.018442 |
| ENSRNOG00000033942 | *Grin2a* | 865.3669 | 2.157938 | 0.546177 | 3.950985 | 7.78E-05 | 0.018442 |
| ENSRNOG00000007069 | *Adhfe1* | 2353.83 | -0.82138 | 0.209025 | -3.92956 | 8.51E-05 | 0.018631 |
| ENSRNOG00000011977 | *Sema5a* | 5717.816 | 1.25498 | 0.319523 | 3.927663 | 8.58E-05 | 0.018631 |
| ENSRNOG00000019317 | *Cplx3* | 167.9889 | -2.54271 | 0.645423 | -3.9396 | 8.16E-05 | 0.018631 |
| ENSRNOG00000031834 | *Nkain4* | 2029.288 | -0.92362 | 0.23512 | -3.9283 | 8.55E-05 | 0.018631 |
| ENSRNOG00000032063 | *Gfral* | 19.53566 | 5.517247 | 1.401527 | 3.936597 | 8.26E-05 | 0.018631 |
| ENSRNOG00000042731 | *Slc25a18* | 3164.674 | -1.34956 | 0.343094 | -3.93349 | 8.37E-05 | 0.018631 |
| ENSRNOG00000037371 | *Xaf1* | 103.1105 | -1.66917 | 0.425426 | -3.92352 | 8.73E-05 | 0.018695 |
| ENSRNOG00000016322 | *Camk2n1* | 18116.9 | -1.47914 | 0.377657 | -3.91663 | 8.98E-05 | 0.018977 |
| ENSRNOG00000014453 | *Anxa5* | 2677.973 | -0.78675 | 0.201636 | -3.90182 | 9.55E-05 | 0.019646 |
| ENSRNOG00000015859 | *Chdh* | 82.24286 | -1.17432 | 0.300881 | -3.90296 | 9.5E-05 | 0.019646 |
| ENSRNOG00000001427 | *Orai2* | 4943.709 | 0.661517 | 0.17006 | 3.889893 | 0.0001 | 0.019729 |
| ENSRNOG00000005669 | *Ca8* | 390.5946 | -1.43167 | 0.367969 | -3.89073 | 9.99E-05 | 0.019729 |
| ENSRNOG00000016897 | *Rlbp1* | 1439.417 | -0.88709 | 0.228317 | -3.88535 | 0.000102 | 0.019729 |
| ENSRNOG00000021468 | *Grm8* | 108.9223 | -2.09147 | 0.537944 | -3.8879 | 0.000101 | 0.019729 |
| ENSRNOG00000027264 | *Dagla* | 6229.52 | 1.251852 | 0.321975 | 3.888042 | 0.000101 | 0.019729 |
| **LPS vs. CUS** | | | | | | | |
| ENSRNOG00000061639 |  | 46.051654 | 3.431289623 | 0.617083435 | 5.560495439 | 2.69E-08 | 0.000408599 |
| ENSRNOG00000001036 | *Rsph10b* | 880.499551 | 1.551537825 | 0.300845844 | 5.157251972 | 2.51E-07 | 0.000965474 |
| ENSRNOG00000006674 | *Rflnb* | 178.3244968 | 1.154453372 | 0.223968322 | 5.154538646 | 2.54E-07 | 0.000965474 |
| ENSRNOG00000015554 | *Ankdd1a* | 161.2166148 | 1.991291367 | 0.384216255 | 5.18273587 | 2.19E-07 | 0.000965474 |
| ENSRNOG00000005882 | *Tle1* | 2075.440038 | 1.189817284 | 0.233396066 | 5.097846348 | 3.44E-07 | 0.001043604 |
| ENSRNOG00000007030 | *Epha7* | 3329.050413 | 1.513689347 | 0.302340959 | 5.00656396 | 5.54E-07 | 0.001402711 |
| ENSRNOG00000039759 | *Gpr34* | 1146.162377 | -0.940559196 | 0.19337203 | -4.863987809 | 1.15E-06 | 0.002496291 |
| ENSRNOG00000007302 | *Fbn1* | 780.284739 | 1.458123157 | 0.309794292 | 4.706746358 | 2.52E-06 | 0.004569255 |
| ENSRNOG00000013036 | *Epha8* | 832.0532305 | 2.200766751 | 0.469650067 | 4.685971331 | 2.79E-06 | 0.004569255 |
| ENSRNOG00000025274 | *Hexb* | 4974.310175 | -0.508390852 | 0.108857202 | -4.670254643 | 3.01E-06 | 0.004569255 |
| ENSRNOG00000004249 | *Tlr7* | 290.8621438 | -0.91247214 | 0.199446758 | -4.575016166 | 4.76E-06 | 0.006575224 |
| ENSRNOG00000046776 | *Msx3* | 26.73331695 | 2.409372664 | 0.530031019 | 4.54572011 | 5.47E-06 | 0.006929696 |
| ENSRNOG00000002930 | *Ppl* | 528.9596871 | 3.19194073 | 0.710181851 | 4.49453999 | 6.97E-06 | 0.007308366 |
| ENSRNOG00000012795 | *Cables1* | 1011.539931 | -1.247724559 | 0.27892091 | -4.473399135 | 7.70E-06 | 0.007308366 |
| ENSRNOG00000013304 | *Arg1* | 662.2911807 | 1.823791849 | 0.407616807 | 4.474280303 | 7.67E-06 | 0.007308366 |
| ENSRNOG00000054695 | *Calcrl* | 739.4975639 | -0.844257834 | 0.188375449 | -4.481782729 | 7.40E-06 | 0.007308366 |
| ENSRNOG00000027230 | *Fhod3* | 481.2655545 | -2.026823231 | 0.456039584 | -4.444401977 | 8.81E-06 | 0.007874753 |
| ENSRNOG00000001427 | *Orai2* | 4943.709391 | 0.751861457 | 0.170125447 | 4.419453219 | 9.90E-06 | 0.00834981 |
| ENSRNOG00000016977 | *Calb2* | 532.4148012 | -1.092191607 | 0.25131222 | -4.345955029 | 1.39E-05 | 0.01108565 |
| ENSRNOG00000040266 | *Cdkl4* | 84.57590178 | 2.059378132 | 0.475702342 | 4.329131787 | 1.50E-05 | 0.01136884 |
| ENSRNOG00000013907 | *Sall1* | 1119.629856 | -0.657660663 | 0.152699022 | -4.306908149 | 1.66E-05 | 0.011429882 |
| ENSRNOG00000017409 | *Wnt6* | 25.80452896 | -5.827865642 | 1.351481759 | -4.31220444 | 1.62E-05 | 0.011429882 |
| ENSRNOG00000049959 | *Igsf21* | 563.1432265 | -1.218733939 | 0.285800294 | -4.264285114 | 2.01E-05 | 0.013243701 |
| ENSRNOG00000011977 | *Sema5a* | 5717.815662 | 1.3589393 | 0.319550555 | 4.252658241 | 2.11E-05 | 0.013369348 |
| ENSRNOG00000026087 | *Igfn1* | 47.0559962 | -1.575476329 | 0.371302432 | -4.243108023 | 2.20E-05 | 0.013393372 |
| ENSRNOG00000002595 | *Dpp10* | 450.5432684 | -1.553878235 | 0.368080461 | -4.221572179 | 2.43E-05 | 0.013664087 |
| ENSRNOG00000008465 | *Tmem176b* | 3258.007313 | -0.781478966 | 0.185487862 | -4.213100295 | 2.52E-05 | 0.013664087 |
| ENSRNOG00000031443 | *Havcr2* | 285.2416936 | -0.899700776 | 0.21318201 | -4.220340993 | 2.44E-05 | 0.013664087 |
| ENSRNOG00000007027 | *Hgf* | 282.4444347 | -1.362591907 | 0.325688709 | -4.183724728 | 2.87E-05 | 0.014519232 |
| ENSRNOG00000029165 | *Stx1a* | 4386.499087 | -1.417535326 | 0.338667708 | -4.185622937 | 2.84E-05 | 0.014519232 |
| ENSRNOG00000038202 | *Calml4* | 565.2687889 | 1.418497802 | 0.340008437 | 4.171948832 | 3.02E-05 | 0.014797294 |
| ENSRNOG00000007089 | *Lgmn* | 3816.42379 | -0.474865713 | 0.114031822 | -4.164326276 | 3.12E-05 | 0.014822247 |
| ENSRNOG00000010266 | *Cd180* | 258.2558902 | -0.782558449 | 0.188743503 | -4.146147737 | 3.38E-05 | 0.015459905 |
| ENSRNOG00000052080 | *Camk2b* | 61094.42518 | 0.922299112 | 0.222733363 | 4.140821559 | 3.46E-05 | 0.015459905 |
| ENSRNOG00000020652 | *Tgfb1* | 410.5715941 | -0.945432779 | 0.228944985 | -4.129519505 | 3.64E-05 | 0.015775823 |
| ENSRNOG00000008086 | *Dpf3* | 166.775412 | 1.029833164 | 0.251153823 | 4.100408079 | 4.12E-05 | 0.017400781 |
| ENSRNOG00000054614 |  | 202.2083686 | -1.233944366 | 0.302415151 | -4.080299417 | 4.50E-05 | 0.018463965 |
| ENSRNOG00000007300 | *C1qtnf6* | 97.94693514 | 1.291400233 | 0.318058904 | 4.060254924 | 4.90E-05 | 0.018933943 |
| ENSRNOG00000048187 | *Epop* | 351.8980259 | -1.88157883 | 0.463869031 | -4.056271717 | 4.99E-05 | 0.018933943 |
| ENSRNOG00000056714 | *Sla* | 155.8861384 | -1.190343366 | 0.293239982 | -4.059280584 | 4.92E-05 | 0.018933943 |
| **EPA vs. CUS** | | | | | | | |
| ENSRNOG00000001036 | *Rsph10b* | 880.499551 | 1.763084973 | 0.300515961 | 5.866859668 | 4.44E-09 | 7.75E-05 |
| ENSRNOG00000007030 | *Epha7* | 3329.050413 | 1.424829615 | 0.302313602 | 4.713084711 | 2.44E-06 | 0.012667845 |
| ENSRNOG00000013036 | *Epha8* | 832.0532305 | 2.196214217 | 0.469528784 | 4.677485799 | 2.90E-06 | 0.012667845 |
| ENSRNOG00000015554 | *Ankdd1a* | 161.2166148 | 1.826776549 | 0.384009197 | 4.757116663 | 1.96E-06 | 0.012667845 |
| ENSRNOG00000002930 | *Ppl* | 528.9596871 | 3.254850905 | 0.710024625 | 4.584138056 | 4.56E-06 | 0.013256496 |
| ENSRNOG00000061639 |  | 46.051654 | 2.860178105 | 0.619582126 | 4.616301836 | 3.91E-06 | 0.013256496 |
| ENSRNOG00000005809 | *Arhgdib* | 525.6784584 | -0.851061368 | 0.188509501 | -4.514686871 | 6.34E-06 | 0.015805492 |
| ENSRNOG00000008182 | *Htra3* | 463.5113397 | -0.989748902 | 0.220869425 | -4.481149449 | 7.42E-06 | 0.016192202 |
| **FLU+LPS vs. CUS** | | | | | | | |
| ENSRNOG00000004554 | *Dcn* | 566.9095907 | -1.509293941 | 0.272147213 | -5.545873216 | 2.92E-08 | 0.000682498 |
| ENSRNOG00000003738 | *Ush2a* | 62.43995164 | 2.493257418 | 0.499433635 | 4.992169616 | 5.97E-07 | 0.006965757 |
| ENSRNOG00000015554 | *Ankdd1a* | 161.2166148 | 2.004091719 | 0.425463654 | 4.710371144 | 2.47E-06 | 0.014424268 |
| ENSRNOG00000061639 |  | 46.051654 | 3.205391125 | 0.675386162 | 4.746012438 | 2.07E-06 | 0.014424268 |
| **FLU+EPA vs. CUS** | | | | | | | |
| ENSRNOG00000021242 | *Adam33* | 7.287877253 | -21.43659697 | 2.324846827 | -9.220649168 | 2.95E-20 | 4.75E-16 |
| ENSRNOG00000006674 | *Rflnb* | 178.3244968 | 1.651526291 | 0.222630788 | 7.418229562 | 1.19E-13 | 9.55E-10 |
| ENSRNOG00000007830 | *Apold1* | 176.3329612 | -2.403072857 | 0.375615296 | -6.397697017 | 1.58E-10 | 8.46E-07 |
| ENSRNOG00000008941 | *Ets1* | 536.0987981 | -1.281069967 | 0.221579331 | -5.7815409 | 7.40E-09 | 2.98E-05 |
| ENSRNOG00000006523 | *Atl2* | 1212.458173 | -0.738841295 | 0.130932711 | -5.642908414 | 1.67E-08 | 5.38E-05 |
| ENSRNOG00000038202 | *Calml4* | 565.2687889 | 1.887813947 | 0.339787954 | 5.555858948 | 2.76E-08 | 7.41E-05 |
| ENSRNOG00000007041 | *Abcg2* | 645.6031228 | -1.566736983 | 0.293923485 | -5.330424627 | 9.80E-08 | 0.000197069 |
| ENSRNOG00000019885 | *Magi3* | 1449.846597 | -0.870604799 | 0.162857875 | -5.345794917 | 9.00E-08 | 0.000197069 |
| ENSRNOG00000008336 | *Tnfrsf11b* | 374.0627582 | -2.401394771 | 0.457197513 | -5.252423081 | 1.50E-07 | 0.000222926 |
| ENSRNOG00000016294 | *Cd4* | 188.7141825 | -1.721591306 | 0.32794523 | -5.249630576 | 1.52E-07 | 0.000222926 |
| ENSRNOG00000023969 | *Herc6* | 279.8592841 | -1.432743617 | 0.271916818 | -5.26905112 | 1.37E-07 | 0.000222926 |
| ENSRNOG00000027456 | *Cdc42bpg* | 579.6898268 | 1.583306847 | 0.304484855 | 5.199952713 | 1.99E-07 | 0.000246721 |
| ENSRNOG00000045558 | *Cd34* | 352.9899336 | -2.412552268 | 0.462794439 | -5.213010496 | 1.86E-07 | 0.000246721 |
| ENSRNOG00000003098 | *Prom1* | 724.6918943 | -1.840493633 | 0.355175678 | -5.181924743 | 2.20E-07 | 0.000252392 |
| ENSRNOG00000018770 | *Pmaip1* | 297.5975034 | -1.60290194 | 0.311992389 | -5.137631556 | 2.78E-07 | 0.00029844 |
| ENSRNOG00000011154 | *Adgrf5* | 1101.337575 | -2.128513091 | 0.416711879 | -5.107877166 | 3.26E-07 | 0.000327631 |
| ENSRNOG00000054809 |  | 74.75496388 | -3.604114712 | 0.708083141 | -5.089959788 | 3.58E-07 | 0.000338968 |
| ENSRNOG00000011912 | *Tmem38a* | 4861.301534 | 0.65929698 | 0.131460545 | 5.015170003 | 5.30E-07 | 0.000449664 |
| ENSRNOG00000047493 | *Slco1a2* | 1902.082244 | -1.795073874 | 0.357957883 | -5.014762797 | 5.31E-07 | 0.000449664 |
| ENSRNOG00000004072 | *Myo1c* | 313.2742176 | -1.163108096 | 0.237823398 | -4.890637775 | 1.01E-06 | 0.000808601 |
| ENSRNOG00000025604 | *Atad2* | 146.3063049 | -1.337470945 | 0.275129423 | -4.861242872 | 1.17E-06 | 0.000893769 |
| ENSRNOG00000000940 | *Flt1* | 1222.890992 | -1.99887156 | 0.415583787 | -4.809791968 | 1.51E-06 | 0.000925721 |
| ENSRNOG00000009269 | *Cga* | 21.5939622 | -17.58030209 | 3.658105789 | -4.805848466 | 1.54E-06 | 0.000925721 |
| ENSRNOG00000012722 | *Ppdpf* | 2035.27742 | 0.828724735 | 0.172615942 | 4.800974489 | 1.58E-06 | 0.000925721 |
| ENSRNOG00000027230 | *Fhod3* | 481.2655545 | -2.206034902 | 0.457054761 | -4.826631489 | 1.39E-06 | 0.000925721 |
| ENSRNOG00000033940 | *Adgrl4* | 292.0829858 | -1.73188327 | 0.361037975 | -4.796955976 | 1.61E-06 | 0.000925721 |
| ENSRNOG00000059894 | *Hmmr* | 100.1195488 | -1.212455662 | 0.252187978 | -4.807745678 | 1.53E-06 | 0.000925721 |
| ENSRNOG00000061639 |  | 46.051654 | 3.011936418 | 0.622612177 | 4.837580327 | 1.31E-06 | 0.000925721 |
| ENSRNOG00000009521 |  | 191.6699903 | 1.204090215 | 0.25169672 | 4.783893148 | 1.72E-06 | 0.000953927 |
| ENSRNOG00000046863 | *C1ql2* | 5582.444048 | 2.259806757 | 0.473421119 | 4.773354346 | 1.81E-06 | 0.000971743 |
| ENSRNOG00000043098 | *Mt2A* | 642.6043846 | 1.500921838 | 0.314963658 | 4.765381023 | 1.88E-06 | 0.000978361 |
| ENSRNOG00000018481 | *Zpr1* | 1807.496142 | 0.520896132 | 0.109455908 | 4.758958583 | 1.95E-06 | 0.000978445 |
| ENSRNOG00000000465 | *Slc39a7* | 3243.550027 | 0.498183218 | 0.104876103 | 4.750207182 | 2.03E-06 | 0.000990795 |
| ENSRNOG00000020173 | *Tie1* | 421.1874057 | -2.053176852 | 0.434490229 | -4.725484525 | 2.30E-06 | 0.001086394 |
| ENSRNOG00000011162 | *Smco4* | 284.5717408 | 1.273034654 | 0.271628365 | 4.686677896 | 2.78E-06 | 0.001241055 |
| ENSRNOG00000013304 | *Arg1* | 662.2911807 | 1.911525871 | 0.407775532 | 4.687691437 | 2.76E-06 | 0.001241055 |
| ENSRNOG00000042888 | *Cdyl2* | 54.67535616 | -1.32858239 | 0.28550386 | -4.653465591 | 3.26E-06 | 0.001419408 |
| ENSRNOG00000002730 | *Rgs5* | 351.8416834 | -2.199517059 | 0.474071487 | -4.639631614 | 3.49E-06 | 0.00147787 |
| ENSRNOG00000016496 | *Ctsc* | 207.9588455 | -1.120211192 | 0.242336529 | -4.622543689 | 3.79E-06 | 0.001487591 |
| ENSRNOG00000024904 | *Pla2g4e* | 159.5844329 | -1.733604072 | 0.374743759 | -4.626105252 | 3.73E-06 | 0.001487591 |
| ENSRNOG00000046776 | *Msx3* | 26.73331695 | 2.47048331 | 0.533462473 | 4.631034861 | 3.64E-06 | 0.001487591 |
| ENSRNOG00000010047 | *Ddit4l* | 333.0496402 | 1.900852481 | 0.412190967 | 4.61158209 | 4.00E-06 | 0.001530909 |
| ENSRNOG00000010047 | *Ddit4l2* | 333.0496402 | 1.900852481 | 0.412190967 | 4.61158209 | 4.00E-06 | 0.001530909 |
| ENSRNOG00000007027 | *Hgf* | 282.4444347 | -1.506872569 | 0.327573133 | -4.600110384 | 4.22E-06 | 0.001580065 |
| ENSRNOG00000032436 | *Tmod3* | 584.6479919 | -0.915653849 | 0.19950007 | -4.589741998 | 4.44E-06 | 0.001622875 |
| ENSRNOG00000046829 | *Kdr* | 199.1018945 | -1.765709058 | 0.385695452 | -4.577987757 | 4.69E-06 | 0.001678617 |
| ENSRNOG00000051891 | *Hist2h4a* | 16.62774487 | 2.720845571 | 0.595125909 | 4.571882236 | 4.83E-06 | 0.001690718 |
| ENSRNOG00000051891 | *Hist1h2ao* | 16.62774487 | 2.720845571 | 0.595125909 | 4.571882236 | 4.83E-06 | 0.001690718 |
| ENSRNOG00000051891 | *LOC100912564* | 16.62774487 | 2.720845571 | 0.595125909 | 4.571882236 | 4.83E-06 | 0.001690718 |
| ENSRNOG00000051891 | *LOC102551184* | 16.62774487 | 2.720845571 | 0.595125909 | 4.571882236 | 4.83E-06 | 0.001690718 |
| ENSRNOG00000007300 | *C1qtnf6* | 97.94693514 | 1.456211494 | 0.318852627 | 4.567036218 | 4.95E-06 | 0.00169345 |
| ENSRNOG00000038574 | *Rassf9* | 101.4201723 | -1.961178212 | 0.432458359 | -4.534952724 | 5.76E-06 | 0.001931349 |
| ENSRNOG00000005882 | *Tle1* | 2075.440038 | 1.054893606 | 0.233624559 | 4.515336958 | 6.32E-06 | 0.001994408 |
| ENSRNOG00000013324 | *Cdh5* | 248.7080682 | -2.42145572 | 0.535658572 | -4.520520804 | 6.17E-06 | 0.001994408 |
| ENSRNOG00000020377 | *Cideb* | 38.76888861 | 2.11150248 | 0.467618831 | 4.515435097 | 6.32E-06 | 0.001994408 |
| ENSRNOG00000001348 | *Erp29* | 2628.482766 | 0.74523837 | 0.165676414 | 4.498156075 | 6.85E-06 | 0.002120952 |
| ENSRNOG00000008323 | *Pitpnm3* | 1179.428134 | -2.1067399 | 0.470363797 | -4.478958448 | 7.50E-06 | 0.002194329 |
| ENSRNOG00000015967 | *Sh3bgrl3* | 5381.287863 | 0.798345392 | 0.17797325 | 4.485760586 | 7.27E-06 | 0.002194329 |
| ENSRNOG00000028841 | *Mt1m* | 1351.425359 | 1.279387924 | 0.285623543 | 4.479280352 | 7.49E-06 | 0.002194329 |
| ENSRNOG00000021899 | *Tmem115* | 1462.507276 | 0.389989688 | 0.08738425 | 4.462928814 | 8.08E-06 | 0.002322905 |
| ENSRNOG00000009037 | *Sulf1* | 145.9085383 | -2.02699636 | 0.455240632 | -4.452582255 | 8.48E-06 | 0.002394975 |
| ENSRNOG00000013269 | *Tnfsf10* | 120.3672437 | -2.526994474 | 0.568016825 | -4.448802156 | 8.64E-06 | 0.002395482 |
| ENSRNOG00000001544 | *Cyyr1* | 671.7972197 | -1.72548894 | 0.389998267 | -4.424350282 | 9.67E-06 | 0.002590281 |
| ENSRNOG00000005900 | *Slc2a6* | 631.6159005 | 0.680538266 | 0.153929805 | 4.421094826 | 9.82E-06 | 0.002590281 |
| ENSRNOG00000006893 | *Ppm1k* | 1126.915113 | -0.75914044 | 0.171638739 | -4.422896853 | 9.74E-06 | 0.002590281 |
| ENSRNOG00000017890 | *Crhbp* | 260.5593482 | -1.142449693 | 0.259083399 | -4.409582771 | 1.04E-05 | 0.002687808 |
| ENSRNOG00000007956 | *Styx* | 73.74133037 | -1.247508314 | 0.28399086 | -4.392776283 | 1.12E-05 | 0.002858204 |
| ENSRNOG00000028436 | *Rprml* | 2664.623055 | 1.259099891 | 0.286973133 | 4.387518369 | 1.15E-05 | 0.002882406 |
| ENSRNOG00000000633 | *Rhobtb1* | 152.3681218 | -1.392063678 | 0.317685784 | -4.381888481 | 1.18E-05 | 0.002912415 |
| ENSRNOG00000004351 | *Slc25a29* | 789.8664224 | 0.501367003 | 0.114606765 | 4.374671961 | 1.22E-05 | 0.002964832 |
| ENSRNOG00000007152 | *Bhlhe40* | 1878.36827 | -1.125434246 | 0.258009304 | -4.361990941 | 1.29E-05 | 0.003095145 |
| ENSRNOG00000013201 | *Tex264* | 2929.686502 | 0.475593371 | 0.109851208 | 4.329432334 | 1.49E-05 | 0.003436231 |
| ENSRNOG00000014034 | *Olfml2a* | 207.9712812 | -2.130451773 | 0.491881062 | -4.331233579 | 1.48E-05 | 0.003436231 |
| ENSRNOG00000025143 | *Icam2* | 133.713765 | -1.843496816 | 0.425699976 | -4.33050721 | 1.49E-05 | 0.003436231 |
| ENSRNOG00000048187 | *Epop* | 351.8980259 | -2.011596805 | 0.464975813 | -4.326239661 | 1.52E-05 | 0.003437286 |
| ENSRNOG00000034022 | *Taf2* | 1944.933455 | -0.420965831 | 0.097411432 | -4.321523874 | 1.55E-05 | 0.00346282 |
| ENSRNOG00000007081 | *Xdh* | 185.9339655 | -1.727774416 | 0.400119803 | -4.318142723 | 1.57E-05 | 0.003468115 |
| ENSRNOG00000017975 | *Dnajb13* | 1582.530768 | 1.613942486 | 0.374159388 | 4.313515946 | 1.61E-05 | 0.003493673 |
| ENSRNOG00000014288 | *Fn1* | 792.9509718 | -1.917382811 | 0.445927819 | -4.299760474 | 1.71E-05 | 0.003668151 |
| ENSRNOG00000011987 | *Cd2ap* | 213.4406195 | -1.704117404 | 0.398178553 | -4.279782002 | 1.87E-05 | 0.003960606 |
| ENSRNOG00000036960 | *Abcc9* | 183.0520332 | -1.843026806 | 0.4311267 | -4.274907596 | 1.91E-05 | 0.003995682 |
| ENSRNOG00000000245 | *Slc16a6* | 239.2162708 | 0.81376384 | 0.191042985 | 4.259585029 | 2.05E-05 | 0.00418443 |
| ENSRNOG00000004221 | *Lgr5* | 22.7215987 | -2.955923355 | 0.694408189 | -4.256751866 | 2.07E-05 | 0.00418443 |
| ENSRNOG00000019689 | *Vwf* | 839.0602377 | -1.798382443 | 0.422545274 | -4.256070428 | 2.08E-05 | 0.00418443 |
| ENSRNOG00000000990 | *Pdap1* | 6332.527572 | 0.720928707 | 0.170718037 | 4.222920565 | 2.41E-05 | 0.004217623 |
| ENSRNOG00000001259 | *Cux2* | 547.7272449 | -1.80488235 | 0.427334197 | -4.223585112 | 2.40E-05 | 0.004217623 |
| ENSRNOG00000001336 | *Orai1* | 492.6948751 | 0.719459947 | 0.169801057 | 4.237075785 | 2.26E-05 | 0.004217623 |
| ENSRNOG00000003296 | *Dck* | 361.8365332 | -0.991133145 | 0.234115584 | -4.233520596 | 2.30E-05 | 0.004217623 |
| ENSRNOG00000003785 | *Usp43* | 21.43869349 | -3.696258228 | 0.872905724 | -4.234430049 | 2.29E-05 | 0.004217623 |
| ENSRNOG00000006749 | *Tmtc3* | 511.9650278 | -0.723912547 | 0.171372676 | -4.224200522 | 2.40E-05 | 0.004217623 |
| ENSRNOG00000010535 | *Cdh18* | 707.2538783 | -0.886988228 | 0.209654044 | -4.230723193 | 2.33E-05 | 0.004217623 |
| ENSRNOG00000012002 | *Iqgap1* | 451.065454 | -1.328686187 | 0.314214202 | -4.228600046 | 2.35E-05 | 0.004217623 |
| ENSRNOG00000012645 | *Mecom* | 93.93182423 | -1.936950667 | 0.455746831 | -4.250058441 | 2.14E-05 | 0.004217623 |
| ENSRNOG00000021013 | *Stx3* | 420.6582012 | -1.062587924 | 0.2514941 | -4.22510081 | 2.39E-05 | 0.004217623 |
| ENSRNOG00000055293 | *Ptprb* | 262.5477417 | -2.358525472 | 0.555838135 | -4.24318758 | 2.20E-05 | 0.004217623 |
| ENSRNOG00000059538 | *Clec2g* | 762.501258 | -1.803466849 | 0.426455117 | -4.228972236 | 2.35E-05 | 0.004217623 |
| ENSRNOG00000010188 | *Satb2* | 503.0841211 | -2.316617649 | 0.548923788 | -4.22029014 | 2.44E-05 | 0.004221256 |
| ENSRNOG00000002848 | *Maoa* | 2518.171315 | -0.610070072 | 0.144639488 | -4.217866642 | 2.47E-05 | 0.004221479 |
| ENSRNOG00000018086 | *Slc22a8* | 344.3008403 | -2.449186801 | 0.581729106 | -4.210184389 | 2.55E-05 | 0.004276627 |
| ENSRNOG00000021682 | *Dcakd* | 1844.506264 | 0.634949194 | 0.150784576 | 4.210969122 | 2.54E-05 | 0.004276627 |
| ENSRNOG00000030715 | *Cfh* | 1877.61545 | -0.819225076 | 0.194834689 | -4.204718784 | 2.61E-05 | 0.004291876 |
| ENSRNOG00000056228 | *Atp10a* | 261.6939162 | -1.348036039 | 0.320581991 | -4.204964955 | 2.61E-05 | 0.004291876 |
| ENSRNOG00000008619 | *Agtrap* | 1776.814736 | 0.480735974 | 0.114511616 | 4.198141564 | 2.69E-05 | 0.004373788 |
| ENSRNOG00000014352 | *Smim12* | 810.4569273 | 0.614312682 | 0.146476704 | 4.193927525 | 2.74E-05 | 0.004411325 |
| ENSRNOG00000015329 | *Kpna2* | 232.3467072 | -0.925968685 | 0.221017336 | -4.189574913 | 2.79E-05 | 0.00445227 |
| ENSRNOG00000018297 | *Ocln* | 387.0194876 | -1.513224018 | 0.36193375 | -4.180942004 | 2.90E-05 | 0.004579402 |
| ENSRNOG00000013798 | *Fnbp1l* | 637.9501517 | -1.721223863 | 0.411993465 | -4.177794086 | 2.94E-05 | 0.004598146 |
| ENSRNOG00000012121 | *Sdf2* | 2919.790454 | 0.59763522 | 0.143141241 | 4.175143475 | 2.98E-05 | 0.004607283 |
| ENSRNOG00000014284 | *Elof1* | 1434.764462 | 0.739132311 | 0.177182802 | 4.17158043 | 3.02E-05 | 0.004635364 |
| ENSRNOG00000011435 | *Osbpl10* | 221.310464 | -1.720578414 | 0.412753015 | -4.168542332 | 3.07E-05 | 0.004653256 |
| ENSRNOG00000002595 | *Dpp10* | 450.5432684 | -1.533590032 | 0.368780937 | -4.158539336 | 3.20E-05 | 0.004694622 |
| ENSRNOG00000004821 | *Sntb1* | 103.5584847 | -1.744264054 | 0.419295184 | -4.159990672 | 3.18E-05 | 0.004694622 |
| ENSRNOG00000014179 | *Rps2* | 5326.598905 | 0.740709867 | 0.177978674 | 4.161789997 | 3.16E-05 | 0.004694622 |
| ENSRNOG00000052376 | *Bend6* | 6563.27103 | -0.576710208 | 0.138696661 | -4.158068427 | 3.21E-05 | 0.004694622 |
| ENSRNOG00000009888 | *Timm8b* | 38.82806078 | 1.497576149 | 0.36135627 | 4.144320362 | 3.41E-05 | 0.004940398 |
| ENSRNOG00000002989 | *Nmt1* | 6899.419328 | 0.631801343 | 0.152739124 | 4.136473536 | 3.53E-05 | 0.004988214 |
| ENSRNOG00000005814 | *NEWGENE_1582771* | 167.4757011 | -1.008966791 | 0.244405511 | -4.128248943 | 3.66E-05 | 0.004988214 |
| ENSRNOG00000005814 | *Katnbl1* | 167.4757011 | -1.008966791 | 0.244405511 | -4.128248943 | 3.66E-05 | 0.004988214 |
| ENSRNOG00000010213 | *Fgd5* | 172.8264415 | -1.734036288 | 0.420245576 | -4.126245198 | 3.69E-05 | 0.004988214 |
| ENSRNOG00000015036 | *Ccn2* | 627.4469516 | -1.413977569 | 0.342360034 | -4.130089459 | 3.63E-05 | 0.004988214 |
| ENSRNOG00000020009 | *Npas4* | 108.9536122 | -2.463002595 | 0.596146898 | -4.131536374 | 3.60E-05 | 0.004988214 |
| ENSRNOG00000029941 | *Grina* | 26049.32016 | 0.630244291 | 0.152715915 | 4.12690643 | 3.68E-05 | 0.004988214 |
| ENSRNOG00000049931 | *LOC288913* | 1594.367668 | 0.646388651 | 0.156132184 | 4.140009037 | 3.47E-05 | 0.004988214 |
| ENSRNOG00000059061 | *Uqcr10* | 5654.889784 | 0.870226002 | 0.210906245 | 4.126127232 | 3.69E-05 | 0.004988214 |
| ENSRNOG00000007227 | *Mien1* | 644.7438377 | 0.707718453 | 0.171952267 | 4.115784377 | 3.86E-05 | 0.005173805 |
| ENSRNOG00000011826 | *Lzts1* | 2804.429114 | -1.513734671 | 0.368506466 | -4.107756069 | 4.00E-05 | 0.005312648 |
| ENSRNOG00000008564 | *Tmem222* | 2220.614802 | 0.514983233 | 0.125767053 | 4.094738818 | 4.23E-05 | 0.005505825 |
| ENSRNOG00000008587 | *Tek* | 625.200133 | -1.48688684 | 0.363051687 | -4.095523841 | 4.21E-05 | 0.005505825 |
| ENSRNOG00000053691 | *Lama5* | 111.9522579 | -1.91311968 | 0.467318385 | -4.093824985 | 4.24E-05 | 0.005505825 |
| ENSRNOG00000002599 | *Grap* | 52.59584438 | -1.807067731 | 0.443649851 | -4.073184579 | 4.64E-05 | 0.005829438 |
| ENSRNOG00000008709 | *Arhgap32* | 6332.748441 | -0.978131305 | 0.239814841 | -4.078693802 | 4.53E-05 | 0.005829438 |
| ENSRNOG00000021735 | *Akr1c15* | 203.8282967 | -1.449091867 | 0.355725773 | -4.073620687 | 4.63E-05 | 0.005829438 |
| ENSRNOG00000024651 | *Greb1* | 72.26991506 | -1.994225968 | 0.489272409 | -4.075901139 | 4.58E-05 | 0.005829438 |
| ENSRNOG00000055023 |  | 347.0729964 | 0.940387778 | 0.231041294 | 4.070215169 | 4.70E-05 | 0.005858473 |
| ENSRNOG00000053058 |  | 101.3147269 | 2.487295132 | 0.612920477 | 4.058104148 | 4.95E-05 | 0.006123199 |
| ENSRNOG00000009802 | *Fam124a* | 845.6280654 | 0.647044287 | 0.15961766 | 4.053713649 | 5.04E-05 | 0.006144785 |
| ENSRNOG00000036816 | *Wls* | 1257.866407 | -0.799865393 | 0.197246246 | -4.055161546 | 5.01E-05 | 0.006144785 |
| ENSRNOG00000002360 | *Gabrg1* | 1671.941893 | -0.993567572 | 0.24601271 | -4.038683911 | 5.38E-05 | 0.006502772 |
| ENSRNOG00000002365 | *Itm2a* | 2048.350903 | -1.290266965 | 0.319939844 | -4.032842394 | 5.51E-05 | 0.006519554 |
| ENSRNOG00000025274 | *Hexb* | 4974.310175 | -0.439610706 | 0.108983638 | -4.033731266 | 5.49E-05 | 0.006519554 |
| ENSRNOG00000042592 | *Rgs10* | 5781.261508 | 1.221220673 | 0.302589283 | 4.035901936 | 5.44E-05 | 0.006519554 |
| ENSRNOG00000025895 | *Cavin2* | 432.2935085 | -1.115978233 | 0.277208637 | -4.025770063 | 5.68E-05 | 0.006669618 |
| ENSRNOG00000001652 | *Erg* | 133.7958696 | -1.782225188 | 0.442914794 | -4.023855635 | 5.73E-05 | 0.00667537 |
| ENSRNOG00000001928 | *Il1rap* | 1298.059565 | 0.915290576 | 0.228619089 | 4.003561473 | 6.24E-05 | 0.007221331 |
| ENSRNOG00000011739 | *Dennd4a* | 1617.344658 | -0.654380744 | 0.163517108 | -4.001909976 | 6.28E-05 | 0.007221331 |
| ENSRNOG00000002141 | *Cd200* | 3750.175607 | -0.511169113 | 0.127873683 | -3.997453587 | 6.40E-05 | 0.007306405 |
| ENSRNOG00000028357 | *Lrrc14b* | 210.5412897 | 0.639064651 | 0.160036058 | 3.993254141 | 6.52E-05 | 0.007384704 |
| ENSRNOG00000001834 | *Mzt2b* | 554.4569934 | 0.613630896 | 0.154483123 | 3.972154899 | 7.12E-05 | 0.0080141 |
| ENSRNOG00000022244 | *Olr1462* | 148.1904851 | 2.162869861 | 0.545205876 | 3.967069978 | 7.28E-05 | 0.008130102 |
| ENSRNOG00000005003 | *Ptprn2* | 5909.990145 | -0.635580436 | 0.16059532 | -3.957652287 | 7.57E-05 | 0.008270407 |
| ENSRNOG00000005132 | *Slc52a3* | 61.57783967 | -1.851523558 | 0.467669622 | -3.959041745 | 7.53E-05 | 0.008270407 |
| ENSRNOG00000011716 | *Degs2* | 124.0546075 | -1.699935733 | 0.429837292 | -3.954835386 | 7.66E-05 | 0.008270407 |
| ENSRNOG00000017905 | *Map1lc3b* | 6064.726661 | 0.459087043 | 0.116077811 | 3.954993974 | 7.65E-05 | 0.008270407 |
| ENSRNOG00000055777 |  | 142.041954 | 1.044721355 | 0.264110033 | 3.955629189 | 7.63E-05 | 0.008270407 |
| ENSRNOG00000021244 | *Hspa12b* | 181.4282595 | -1.65777645 | 0.419644745 | -3.950428238 | 7.80E-05 | 0.008368032 |
| ENSRNOG00000012794 | *Grhpr* | 1486.39409 | 0.8179382 | 0.20715511 | 3.948433609 | 7.87E-05 | 0.008382168 |
| ENSRNOG00000005868 | *Ttc21b* | 261.1105751 | -0.952447702 | 0.241431382 | -3.945003731 | 7.98E-05 | 0.008391906 |
| ENSRNOG00000017659 | *Hs3st2* | 259.27256 | -1.422907973 | 0.360645953 | -3.945442787 | 7.97E-05 | 0.008391906 |
| ENSRNOG00000016362 | *Gpr4* | 86.45470419 | -2.03870961 | 0.517850507 | -3.936869006 | 8.26E-05 | 0.008617366 |
| ENSRNOG00000018243 | *Ubfd1* | 756.7457394 | 0.565596472 | 0.143735194 | 3.934989447 | 8.32E-05 | 0.008617366 |
| ENSRNOG00000020178 | *Cope* | 2999.736915 | 0.577100604 | 0.146696226 | 3.93398398 | 8.35E-05 | 0.008617366 |
| ENSRNOG00000024799 | *Cd93* | 121.7454579 | -2.151917443 | 0.547648789 | -3.929374965 | 8.52E-05 | 0.008728256 |
| ENSRNOG00000000394 | *Srgn* | 571.6287321 | -1.371405058 | 0.350259479 | -3.915397415 | 9.03E-05 | 0.009175166 |
| ENSRNOG00000013179 | *Tinagl1* | 185.0710566 | -1.762094981 | 0.450169022 | -3.914296396 | 9.07E-05 | 0.009175166 |
| ENSRNOG00000057569 | *Ahnak* | 1024.174471 | -1.448621151 | 0.370700548 | -3.907793386 | 9.31E-05 | 0.009366683 |
| ENSRNOG00000004303 | *Timp3* | 983.3192333 | -1.133169745 | 0.290230582 | -3.904377473 | 9.45E-05 | 0.009440956 |
| ENSRNOG00000003888 | *Rgs13* | 238.2829283 | 2.326985382 | 0.598528699 | 3.887842614 | 0.000101139 | 0.010045242 |
| ENSRNOG00000022839 | *Ifit3* | 40.4688516 | -2.7790993 | 0.71669819 | -3.877642415 | 0.000105474 | 0.010411473 |
| ENSRNOG00000002101 | *Paics* | 266.0609765 | -0.78822399 | 0.203865607 | -3.866390222 | 0.000110458 | 0.010837022 |
| ENSRNOG00000047218 | *Clic5* | 209.0185758 | -1.73192565 | 0.448368398 | -3.862729081 | 0.000112127 | 0.010934114 |
| ENSRNOG00000026163 | *Cpt1c* | 2959.317755 | 0.394607324 | 0.102251049 | 3.859200727 | 0.000113758 | 0.011026348 |
| ENSRNOG00000001628 | *Pcp4* | 6226.207926 | 1.189425366 | 0.308968986 | 3.849659411 | 0.000118282 | 0.011327906 |
| ENSRNOG00000012536 | *Sgms1* | 1515.298759 | -0.725614224 | 0.188609945 | -3.847168414 | 0.000119491 | 0.011327906 |
| ENSRNOG00000023337 | *Sema3a* | 14.57117174 | -3.672360593 | 0.955167601 | -3.844729019 | 0.000120686 | 0.011327906 |
| ENSRNOG00000026177 |  | 398.1540538 | 1.407077184 | 0.365918711 | 3.845327226 | 0.000120392 | 0.011327906 |
| ENSRNOG00000054224 | *Bloc1s4* | 722.4745664 | 0.683868061 | 0.177786547 | 3.846568094 | 0.000119784 | 0.011327906 |
| ENSRNOG00000059859 |  | 219.6045706 | -1.221678361 | 0.31782253 | -3.843901069 | 0.000121094 | 0.011327906 |
| ENSRNOG00000021053 | *Lsr* | 228.297886 | -1.57966717 | 0.411631754 | -3.837573645 | 0.000124256 | 0.011490107 |
| ENSRNOG00000022491 | *Wdr76* | 49.46560255 | -1.578086822 | 0.411133725 | -3.838378427 | 0.00012385 | 0.011490107 |
| ENSRNOG00000005715 | *Lgr4* | 972.4092175 | -0.779000404 | 0.203231375 | -3.83307156 | 0.000126553 | 0.011573032 |
| ENSRNOG00000010855 | *Mrpl57* | 996.8876948 | 0.863097015 | 0.225188538 | 3.832775071 | 0.000126706 | 0.011573032 |
| ENSRNOG00000046316 | *Tomm6* | 3311.749137 | 0.799967972 | 0.208781486 | 3.831603976 | 0.000127311 | 0.011573032 |
| ENSRNOG00000003116 | *Dph1* | 1388.260912 | 0.657209172 | 0.171654865 | 3.828666161 | 0.00012884 | 0.011646234 |
| ENSRNOG00000039759 | *Gpr34* | 1146.162377 | -0.740054603 | 0.193486662 | -3.824835241 | 0.00013086 | 0.01176274 |
| ENSRNOG00000051977 | *Mmrn2* | 175.998097 | -1.902357301 | 0.498012494 | -3.819898742 | 0.000133506 | 0.011933995 |
| ENSRNOG00000014169 |  | 36.37954479 | 1.639339051 | 0.430336539 | 3.809434949 | 0.000139285 | 0.01231369 |
| ENSRNOG00000027730 | *Nxpe1* | 49.39349363 | -2.178853811 | 0.571792188 | -3.810569395 | 0.000138647 | 0.01231369 |
| ENSRNOG00000002345 | *Rasgef1b* | 281.0824145 | -1.240822476 | 0.326114974 | -3.804862006 | 0.000141883 | 0.012474873 |
| ENSRNOG00000017786 | *Acta1* | 620.7793408 | 0.665169113 | 0.175092469 | 3.79895902 | 0.000145305 | 0.012677213 |
| ENSRNOG00000019319 | *Fchsd2* | 1265.152926 | -0.579088432 | 0.152470377 | -3.79803896 | 0.000145845 | 0.012677213 |
| ENSRNOG00000019509 | *Fbxl15* | 1430.509992 | 0.833880646 | 0.219624515 | 3.79684685 | 0.000146548 | 0.012677213 |
| ENSRNOG00000014890 | *Mrpl43* | 1906.60951 | 0.629221527 | 0.165988241 | 3.790759642 | 0.000150187 | 0.012922528 |
| ENSRNOG00000040266 | *Cdkl4* | 84.57590178 | 1.812643522 | 0.478513384 | 3.788072771 | 0.00015182 | 0.012993564 |
| ENSRNOG00000005286 | *Coch* | 412.9341506 | -3.260649859 | 0.861725472 | -3.783861526 | 0.000154414 | 0.012996435 |
| ENSRNOG00000008666 | *Etl4* | 1286.155398 | -0.759324428 | 0.200752583 | -3.782389332 | 0.00015533 | 0.012996435 |
| ENSRNOG00000011925 | *Fam83b* | 13.51397118 | -3.337391749 | 0.883049077 | -3.779395546 | 0.00015721 | 0.012996435 |
| ENSRNOG00000012061 | *Prkcb* | 5447.624019 | -0.615565844 | 0.162949456 | -3.777648975 | 0.000158316 | 0.012996435 |
| ENSRNOG00000012681 | *Lgals9* | 416.3869335 | -1.500699079 | 0.397236624 | -3.777846726 | 0.00015819 | 0.012996435 |
| ENSRNOG00000013248 | *Wwc2* | 1261.920905 | -0.409498101 | 0.108258227 | -3.7826049 | 0.000155196 | 0.012996435 |
| ENSRNOG00000020659 | *Mrpl4* | 2204.753023 | 0.523141463 | 0.138465483 | 3.778136275 | 0.000158006 | 0.012996435 |
| ENSRNOG00000048230 | *LOC300308* | 151.754022 | -2.176708555 | 0.5756463 | -3.781329883 | 0.000155993 | 0.012996435 |
| ENSRNOG00000003869 | *Sod3* | 1289.179632 | 0.634156416 | 0.1681064 | 3.772351421 | 0.000161716 | 0.01306642 |
| ENSRNOG00000019811 | *Timm23* | 873.5609962 | 0.562942736 | 0.149249133 | 3.771832535 | 0.000162053 | 0.01306642 |
| ENSRNOG00000019811 | *LOC100362432* | 873.5609962 | 0.562942736 | 0.149249133 | 3.771832535 | 0.000162053 | 0.01306642 |
| ENSRNOG00000026132 | *Trdmt1* | 180.3857419 | -0.956288942 | 0.253459716 | -3.772942529 | 0.000161333 | 0.01306642 |
| ENSRNOG00000054058 | *Osbpl1a* | 5099.776862 | -0.717213598 | 0.190178101 | -3.771273322 | 0.000162417 | 0.01306642 |
| ENSRNOG00000057159 | *Rsl1* | 74.1247242 | -1.375248201 | 0.365431308 | -3.76335626 | 0.000167648 | 0.013420189 |
| ENSRNOG00000001189 | *Sik1* | 147.214959 | -1.079432272 | 0.287745954 | -3.751337798 | 0.000175894 | 0.013599792 |
| ENSRNOG00000009401 | *Lmo2* | 1011.618812 | 0.774118284 | 0.206414856 | 3.750303152 | 0.000176621 | 0.013599792 |
| ENSRNOG00000011589 | *Camk2d* | 1173.796341 | -1.361056597 | 0.36230903 | -3.756617921 | 0.000172225 | 0.013599792 |
| ENSRNOG00000016299 | *Klf4* | 242.6709306 | -1.856174457 | 0.493966362 | -3.757694044 | 0.000171486 | 0.013599792 |
| ENSRNOG00000019247 | *Plekhj1* | 760.0250868 | 0.723083456 | 0.192744351 | 3.751515679 | 0.000175769 | 0.013599792 |
| ENSRNOG00000025416 | *Dpy19l4* | 186.6967362 | -1.037013077 | 0.276442423 | -3.751280518 | 0.000175934 | 0.013599792 |
| ENSRNOG00000028713 | *Acvrl1* | 303.9459409 | -1.732235897 | 0.461748819 | -3.751467955 | 0.000175802 | 0.013599792 |
| ENSRNOG00000036680 | *Notum* | 95.91489342 | -2.125392283 | 0.566732475 | -3.750256739 | 0.000176654 | 0.013599792 |
| ENSRNOG00000012563 | *Arhgap29* | 350.7629407 | -1.384444176 | 0.369642189 | -3.745362997 | 0.000180133 | 0.013736218 |
| ENSRNOG00000019178 | *Taf10* | 2251.979599 | 0.64628368 | 0.172528519 | 3.745952747 | 0.00017971 | 0.013736218 |
| ENSRNOG00000016281 | *Col4a1* | 667.4360368 | -1.42674244 | 0.381424018 | -3.740567906 | 0.000183605 | 0.013930708 |
| ENSRNOG00000020169 | *Gimap8* | 63.25658202 | -1.584078289 | 0.423611388 | -3.739461058 | 0.000184415 | 0.013930708 |
| ENSRNOG00000009414 | *Creld1* | 3327.24464 | 0.420929396 | 0.112774257 | 3.732495393 | 0.000189592 | 0.01425485 |
| ENSRNOG00000005506 | *Arhgef5* | 32.39753444 | -2.274458512 | 0.609982349 | -3.728728403 | 0.000192448 | 0.014402304 |
| ENSRNOG00000018797 | *Myrip* | 3081.604283 | -0.767393059 | 0.205951501 | -3.726086267 | 0.000194476 | 0.014486652 |
| ENSRNOG00000000943 |  | 25.18493284 | 1.751603171 | 0.470490459 | 3.722930267 | 0.000196924 | 0.014553963 |
| ENSRNOG00000017841 | *Psme3ip1* | 2018.962903 | 0.393179461 | 0.10561983 | 3.722591314 | 0.000197189 | 0.014553963 |
| ENSRNOG00000050258 | *Ccnd3* | 1785.965603 | 0.435700024 | 0.117141811 | 3.719423658 | 0.000199678 | 0.014670398 |
| ENSRNOG00000014243 | *Pear1* | 162.3865951 | -1.955226309 | 0.525930171 | -3.717653817 | 0.000201082 | 0.014706374 |
| ENSRNOG00000002215 | *Mylk* | 914.789805 | -0.485292364 | 0.130826029 | -3.709448082 | 0.000207712 | 0.015054409 |
| ENSRNOG00000053035 |  | 7.608683568 | 5.41916405 | 1.460691804 | 3.70999826 | 0.000207261 | 0.015054409 |
| ENSRNOG00000056716 | *Zbtb20* | 1774.218258 | 1.364746519 | 0.368157162 | 3.706967187 | 0.000209756 | 0.01513442 |
| ENSRNOG00000009381 | *Mapk6* | 2110.645824 | -0.550310554 | 0.148690775 | -3.701040332 | 0.000214717 | 0.015423223 |
| ENSRNOG00000019920 | *Doc2a* | 286.1626155 | -1.120837631 | 0.30301549 | -3.698945004 | 0.000216498 | 0.015481977 |
| ENSRNOG00000010031 | *Vtn* | 344.4881349 | -1.936958403 | 0.524158492 | -3.695367783 | 0.000219569 | 0.015563263 |
| ENSRNOG00000050348 | *LOC684270* | 726.2361648 | 0.745761132 | 0.20179152 | 3.695701053 | 0.000219281 | 0.015563263 |
| ENSRNOG00000017409 | *Wnt6* | 25.80452896 | -4.791049625 | 1.297743194 | -3.691831827 | 0.000222645 | 0.015712074 |
| ENSRNOG00000005608 | *Tead4* | 43.23993482 | -1.933202212 | 0.52466073 | -3.684671066 | 0.000228998 | 0.016089872 |
| ENSRNOG00000018639 | *Zfand2b* | 1444.30573 | 0.839028154 | 0.227997839 | 3.679982931 | 0.00023325 | 0.016317328 |
| ENSRNOG00000025287 | *RGD1565611* | 81.48539128 | 1.844085203 | 0.50176523 | 3.675195274 | 0.000237668 | 0.01655442 |
| ENSRNOG00000004206 | *Glrx5* | 2088.911693 | 0.58462802 | 0.159177847 | 3.672797636 | 0.000239909 | 0.016564821 |
| ENSRNOG00000014835 | *Il1rl1* | 36.34225877 | -1.396521896 | 0.380343429 | -3.67173925 | 0.000240905 | 0.016564821 |
| ENSRNOG00000016103 | *Nkd2* | 220.9459166 | -1.616147991 | 0.440022544 | -3.672875433 | 0.000239836 | 0.016564821 |
| ENSRNOG00000010917 | *Bmp5* | 13.57935482 | -2.803122196 | 0.764147327 | -3.668300727 | 0.000244168 | 0.016717708 |
| ENSRNOG00000016366 | *Colec12* | 293.0481307 | -1.445966206 | 0.394406722 | -3.666180433 | 0.0002462 | 0.016785432 |
| ENSRNOG00000002697 | *Mtmr1* | 2330.31102 | 0.519547971 | 0.142295534 | 3.651189571 | 0.000261028 | 0.017215482 |
| ENSRNOG00000003772 | *Csrp2* | 458.353737 | -1.030783011 | 0.282327808 | -3.651014825 | 0.000261206 | 0.017215482 |
| ENSRNOG00000004489 | *Adgre5* | 178.1378086 | -1.314137883 | 0.359619368 | -3.654246681 | 0.000257938 | 0.017215482 |
| ENSRNOG00000009329 | *Nr1d1* | 1724.700509 | -0.652276815 | 0.178391113 | -3.656442321 | 0.00025574 | 0.017215482 |
| ENSRNOG00000013994 | *Enpp1* | 148.5106929 | -0.931719108 | 0.255172277 | -3.651333598 | 0.000260882 | 0.017215482 |
| ENSRNOG00000021233 | *Itpa* | 4762.523929 | 0.904065987 | 0.247509664 | 3.652649246 | 0.000259549 | 0.017215482 |
| ENSRNOG00000022868 | *Ell3* | 333.2902025 | 0.823123818 | 0.225368417 | 3.652347696 | 0.000259854 | 0.017215482 |
| ENSRNOG00000027309 |  | 44.70006969 | -2.613493867 | 0.7150148 | -3.655160517 | 0.000257021 | 0.017215482 |
| ENSRNOG00000053070 |  | 2174.153534 | 0.575531431 | 0.157675495 | 3.650100677 | 0.000262138 | 0.017215482 |
| ENSRNOG00000021084 | *Mpeg1* | 1185.262527 | -1.328552106 | 0.3642539 | -3.647324314 | 0.000264985 | 0.017331767 |
| ENSRNOG00000010966 | *Itgb1* | 2963.497867 | -0.606751335 | 0.166406329 | -3.646203477 | 0.000266143 | 0.017337025 |
| ENSRNOG00000000306 | *Smpd2* | 2113.317468 | 1.027745446 | 0.282673956 | 3.635798148 | 0.000277121 | 0.017643997 |
| ENSRNOG00000011719 | *Ngb* | 317.3334436 | 0.986771485 | 0.271292359 | 3.637299218 | 0.000275512 | 0.017643997 |
| ENSRNOG00000014010 | *Gfra2* | 215.3202711 | -1.998094452 | 0.549605542 | -3.635506377 | 0.000277435 | 0.017643997 |
| ENSRNOG00000048174 | *Uqcrq* | 4658.100678 | 0.786608311 | 0.216278354 | 3.637018208 | 0.000275812 | 0.017643997 |
| ENSRNOG00000048733 | *Nup62* | 1690.941399 | 0.550769721 | 0.151379769 | 3.638331102 | 0.00027441 | 0.017643997 |
| ENSRNOG00000050869 | *Cebpd* | 543.78835 | 1.174596255 | 0.322963444 | 3.636932532 | 0.000275904 | 0.017643997 |
| ENSRNOG00000014908 | *Sf3b5* | 1988.745015 | 0.89078112 | 0.245149509 | 3.633623926 | 0.000279468 | 0.017703315 |
| ENSRNOG00000050964 | *LOC100911068* | 39.37340087 | -2.236465184 | 0.615831198 | -3.631620469 | 0.000281647 | 0.017771381 |
| ENSRNOG00000046585 | *Chmp4b* | 1503.145286 | 0.434362897 | 0.119702962 | 3.628672913 | 0.000284882 | 0.017905273 |
| ENSRNOG00000021916 | *Slc16a12* | 130.0909617 | -1.199479871 | 0.330648577 | -3.627657744 | 0.000286004 | 0.017905856 |
| ENSRNOG00000009849 | *Eef1akmt1* | 610.3529827 | 0.617574319 | 0.170385572 | 3.624569333 | 0.000289443 | 0.017912091 |
| ENSRNOG00000014048 | *Cyld* | 1293.571946 | -0.879259598 | 0.24253603 | -3.625274142 | 0.000288655 | 0.017912091 |
| ENSRNOG00000014672 | *Hs3st6* | 6.871577985 | -6.946140267 | 1.916158292 | -3.625034683 | 0.000288923 | 0.017912091 |
| ENSRNOG00000002959 | *Shroom4* | 195.7983687 | -1.313889267 | 0.363076462 | -3.618767405 | 0.00029601 | 0.01809199 |
| ENSRNOG00000026134 |  | 42.27891627 | -1.417901935 | 0.391898329 | -3.618035164 | 0.000296848 | 0.01809199 |
| ENSRNOG00000042274 | *Fbxo31* | 3144.692199 | 0.477982342 | 0.132071799 | 3.619109801 | 0.000295618 | 0.01809199 |
| ENSRNOG00000046109 | *LOC689574* | 1115.829589 | 0.612593104 | 0.169274798 | 3.618926797 | 0.000295827 | 0.01809199 |
| ENSRNOG00000033217 | *Esam* | 454.2529056 | -1.430674265 | 0.395628166 | -3.616209327 | 0.000298949 | 0.01815126 |
| ENSRNOG00000001645 | *Filip1l* | 103.0825216 | -1.622920459 | 0.449157664 | -3.613253405 | 0.000302379 | 0.018229561 |
| ENSRNOG00000019949 | *Mrps12* | 732.9368121 | 0.638558077 | 0.176731876 | 3.613146045 | 0.000302504 | 0.018229561 |
| ENSRNOG00000008301 | *Tagln2* | 682.4459561 | -1.382724387 | 0.382842906 | -3.611727854 | 0.000304164 | 0.01826117 |
| ENSRNOG00000042667 |  | 118.8503971 | -0.935370692 | 0.259258775 | -3.607865121 | 0.000308727 | 0.01846623 |
| ENSRNOG00000004819 | *Porcn* | 3156.419341 | 0.703321626 | 0.195137465 | 3.604236766 | 0.000313072 | 0.018587902 |
| ENSRNOG00000052795 | *Itpr3* | 107.299195 | -1.617206488 | 0.448629798 | -3.604768327 | 0.000312432 | 0.018587902 |
| ENSRNOG00000012058 | *Egflam* | 191.7785964 | -1.710445637 | 0.474745782 | -3.602866425 | 0.000314727 | 0.018617507 |
| ENSRNOG00000009063 | *Dnajc15* | 1387.769871 | 0.808486458 | 0.224821402 | 3.596127637 | 0.000322989 | 0.018829344 |
| ENSRNOG00000038864 |  | 35.20345958 | 1.921458058 | 0.534154333 | 3.597196428 | 0.000321666 | 0.018829344 |
| ENSRNOG00000055259 | *Phldb3* | 17.72803368 | -2.719676236 | 0.756242621 | -3.596301185 | 0.000322774 | 0.018829344 |
| ENSRNOG00000056212 | *Flt3lg* | 96.19332031 | 1.34145813 | 0.372997378 | 3.596427772 | 0.000322617 | 0.018829344 |
| ENSRNOG00000054194 |  | 51.17415255 | 1.236883285 | 0.344042046 | 3.59515152 | 0.000324203 | 0.018831854 |
| ENSRNOG00000016571 | *Ngf* | 166.2442697 | 0.902334205 | 0.251063941 | 3.594041427 | 0.000325588 | 0.018844286 |
| ENSRNOG00000011422 |  | 1060.821541 | -0.501176906 | 0.139503454 | -3.592577031 | 0.000327424 | 0.018882614 |
| ENSRNOG00000003887 | *Lgi2* | 956.4759591 | -0.678546838 | 0.18904492 | -3.589341821 | 0.000331514 | 0.01905021 |
| ENSRNOG00000001143 | *Cit* | 1178.971231 | -1.453468034 | 0.405184455 | -3.587176196 | 0.000334278 | 0.019140713 |
| ENSRNOG00000054155 |  | 158.8777393 | -1.412836795 | 0.393968045 | -3.58617104 | 0.000335569 | 0.019146467 |
| ENSRNOG00000019352 | *Emc6* | 1078.760238 | 0.620313504 | 0.173108627 | 3.58337719 | 0.00033918 | 0.019216234 |
| ENSRNOG00000057806 | *Trpm7* | 818.5702 | -0.974669035 | 0.271969376 | -3.583745532 | 0.000338702 | 0.019216234 |
| ENSRNOG00000017720 | *Mxd1* | 1089.987407 | -0.538852248 | 0.150476617 | -3.580969974 | 0.000342321 | 0.019326122 |
| ENSRNOG00000001185 | *RGD1311899* | 11732.51426 | 0.618041469 | 0.172820311 | 3.576208519 | 0.000348614 | 0.019441859 |
| ENSRNOG00000006898 | *Mrps16* | 1465.542602 | 0.790025261 | 0.220938766 | 3.575765705 | 0.000349204 | 0.019441859 |
| ENSRNOG00000021719 | *Slfn5* | 460.7208893 | -1.422405698 | 0.397617225 | -3.577324143 | 0.00034713 | 0.019441859 |
| ENSRNOG00000031313 |  | 437.9514261 | 1.067618959 | 0.298436834 | 3.577369945 | 0.000347069 | 0.019441859 |
| ENSRNOG00000011058 | *Utrn* | 1558.92664 | -1.098730781 | 0.307489979 | -3.573224673 | 0.000352612 | 0.019496655 |
| ENSRNOG00000020455 | *Cst6* | 2674.133204 | 1.237911706 | 0.346401295 | 3.57363475 | 0.00035206 | 0.019496655 |
| ENSRNOG00000015304 | *Tmem160* | 2136.97014 | 0.783862194 | 0.219876193 | 3.565016218 | 0.000363834 | 0.019979809 |
| ENSRNOG00000029773 | *Atm* | 895.2636029 | -0.631836215 | 0.177212237 | -3.565420915 | 0.000363273 | 0.019979809 |

**Supplementary Table S4.** Statistical data for pathways enriched in subiculum in experimental groups vs. control group or in experimental groups vs. stress group using the KEGG- enrichment analysis that was performed on normalized and log2-transformed counts by general applicable gene set enrichment for pathway analysis (GAGE) package, using two-tailed t-test for group comparison of differential expression of gene sets False discovery rate was set at 0.01(6) for control comparison and 0.02 for CUS comparison. p.geomean – geometric mean of the individual p-values from multiple single array based gene set tests, stat.mean - mean of the individual statistics from multiple single array based gene set tests. Pathways are listed as sorted by q value. CUS – chronic unpredictable stress, FLU – fluoxetine, EPA - eicosapentaenoic acid, LPS – lipopolysaccharide.

| **Pathway code and name** | **p.geomean** | **stat.mean** | **p.val** | **q.val** | **set.size** |
| --- | --- | --- | --- | --- | --- |
| **CUS vs. Control Down** | | | | | |
| rno03040 Spliceosome | 0.001116 | -2.77084 | 1.37E-06 | 0.000415 | 130 |
| **CUS vs. Control Up** | | | | | |
| rno03010 Ribosome | 1.87E-06 | 3.180916 | 6.59E-08 | 1.99E-05 | 179 |
| rno05140 Leishmaniasis | 0.0021 | 2.846318 | 6.75E-07 | 0.000102 | 67 |
| rno05145 Toxoplasmosis | 0.002734 | 2.688692 | 2.16E-06 | 0.000217 | 102 |
| rno04380 Osteoclast differentiation | 0.004974 | 2.554533 | 5.79E-06 | 0.000437 | 119 |
| rno04621 NOD-like receptor signaling pathway | 0.007651 | 2.394708 | 1.88E-05 | 0.001134 | 157 |
| rno05152 Tuberculosis | 0.00891 | 2.287501 | 4.13E-05 | 0.001835 | 157 |
| rno05323 Rheumatoid arthritis | 0.011466 | 2.292509 | 4.25E-05 | 0.001835 | 77 |
| rno04611 Platelet activation | 0.01045 | 2.141895 | 0.000119 | 0.004475 | 119 |
| rno04145 Phagosome | 0.013019 | 2.082059 | 0.000169 | 0.005672 | 165 |
| rno03320 PPAR signaling pathway | 0.014661 | 2.065361 | 0.000205 | 0.006191 | 77 |
| rno01200 Carbon metabolism | 0.011211 | 2.04309 | 0.000235 | 0.006456 | 112 |
| rno04510 Focal adhesion | 0.007959 | 2.015244 | 0.000269 | 0.006685 | 196 |
| rno04933 AGE-RAGE signaling pathway in diabetic complications | 0.015498 | 2.008801 | 0.000288 | 0.006685 | 99 |
| rno04217 Necroptosis | 0.023356 | 1.973863 | 0.000334 | 0.007079 | 139 |
| rno00260 Glycine. serine and threonine metabolism | 0.016751 | 2.009176 | 0.000352 | 0.007079 | 37 |
| rno00190 Oxidative phosphorylation | 0.002601 | 2.003442 | 0.000389 | 0.007341 | 121 |
| rno05321 Inflammatory bowel disease (IBD) | 0.022603 | 1.956095 | 0.00042 | 0.007468 | 53 |
| rno05020 Prion diseases | 0.010007 | 1.899032 | 0.000541 | 0.008456 | 255 |
| rno04066 HIF-1 signaling pathway | 0.027278 | 1.896729 | 0.000548 | 0.008456 | 105 |
| rno05133 Pertussis | 0.028273 | 1.898905 | 0.00056 | 0.008456 | 68 |
| rno04062 Chemokine signaling pathway | 0.025022 | 1.814902 | 0.000879 | 0.01243 | 174 |
| rno00010 Glycolysis / Gluconeogenesis | 0.027772 | 1.82734 | 0.000905 | 0.01243 | 59 |
| rno05132 Salmonella infection | 0.026767 | 1.799239 | 0.000949 | 0.012461 | 243 |
| rno04060 Cytokine-cytokine receptor interaction | 0.020816 | 1.776098 | 0.001098 | 0.013819 | 236 |
| rno04932 Non-alcoholic fatty liver disease (NAFLD) | 0.028614 | 1.766863 | 0.001172 | 0.014159 | 149 |
| rno05150 Staphylococcus aureus infection | 0.031979 | 1.744943 | 0.001382 | 0.016051 | 83 |
| **FLU vs. Control Down** | | | | | |
| rno03010 Ribosome | 2E-07 | -3.73431 | 4.18E-10 | 1.26E-07 | 179 |
| rno03040 Spliceosome | 0.000367 | -3.24524 | 1.91E-08 | 2.89E-06 | 130 |
| rno04110 Cell cycle | 0.008494 | -2.13112 | 0.000131 | 0.013154 | 122 |
| **FLU vs. Control Up** | | | | | |
| rno04020 Calcium signaling pathway | 0.001913 | 2.751187 | 1.12E-06 | 0.000337 | 231 |
| rno04724 Glutamatergic synapse | 0.007743 | 2.219188 | 7.25E-05 | 0.010942 | 111 |
| **LPS vs. Control Down** | | | | | |
| rno04110 Cell cycle | 0.00492 | -2.27823 | 4.9E-05 | 0.014792 | 122 |
| **LPS vs. Control Up** | | | | | |
| rno05020 Prion diseases | 0.00045 | 2.409313 | 1.94E-05 | 0.005844 | 255 |
| **EPA vs. Control Down** | | | | | |
| rno03040 Spliceosome | 0.001324 | -2.50647 | 1.12E-05 | 0.003382 | 130 |
| **EPA vs. Control Up** | | | | | |
| - | - | - | - | - | - |
| **FLU+LPS vs. Control Down** | | | | | |
| rno03040 Spliceosome | 0.000551 | -3.12431 | 7.54E-06 | 0.002277 | 130 |
| **FLU+LPS vs. Control Up** | | | | | |
| - | - | - | - | - | - |
| **FLU+EPA vs. Control Down** | | | | | |
| rno04512 ECM-receptor interaction | 0.003045549 | -2.575844553 | 5.91257E-06 | 0.001146623 | 83 |
| rno04360 Axon guidance | 0.000688939 | -2.532777495 | 7.59353E-06 | 0.001146623 | 178 |
| rno04070 Phosphatidylinositol signaling system | 0.005153775 | -2.174235937 | 0.000107815 | 0.010853397 | 92 |
| rno04974 Protein digestion and absorption | 0.011945634 | -2.108683635 | 0.000153851 | 0.011615737 | 94 |
| rno04110 Cell cycle | 0.006491793 | -2.046110298 | 0.000236615 | 0.014291571 | 122 |
| rno04512 ECM-receptor interaction | 0.003045549 | -2.575844553 | 5.91257E-06 | 0.001146623 | 83 |
| **FLU+EPA vs. Control Up** | | | | | |
| rno03010 Ribosome | 4.53E-20 | 8.968605 | 2.43E-46 | 7.33782E-44 | 179 |
| rno00190 Oxidative phosphorylation | 3.7E-10 | 6.322148 | 1.24E-25 | 1.86961E-23 | 121 |
| rno05020 Prion diseases | 1.51E-06 | 4.672288 | 6.39E-16 | 6.43285E-14 | 255 |
| rno05012 Parkinson's disease | 3.89E-06 | 4.281548 | 1.32E-13 | 9.99705E-12 | 234 |
| rno05016 Huntington's disease | 2.61E-05 | 4.006004 | 3E-12 | 1.81443E-10 | 279 |
| rno04932 Non-alcoholic fatty liver disease (NAFLD) | 9.38E-05 | 3.749106 | 7.58E-11 | 3.8137E-09 | 149 |
| rno01200 Carbon metabolism | 0.00032 | 3.412904 | 3.1E-09 | 1.33769E-07 | 112 |
| rno05014 Amyotrophic lateral sclerosis (ALS) | 0.000493 | 3.223331 | 1.38E-08 | 5.21762E-07 | 342 |
| rno05010 Alzheimer's disease | 0.001748 | 2.896199 | 2.9E-07 | 9.7198E-06 | 351 |
| rno00010 Glycolysis / Gluconeogenesis | 0.002472 | 2.860425 | 6.26E-07 | 1.88916E-05 | 59 |
| rno01230 Biosynthesis of amino acids | 0.009807 | 2.32021 | 3.65E-05 | 0.001002333 | 73 |
| rno03050 Proteasome | 0.006454 | 2.343971 | 4.36E-05 | 0.001097172 | 46 |
| rno04146 Peroxisome | 0.011007 | 2.277768 | 4.72E-05 | 0.001097585 | 84 |
| rno00630 Glyoxylate and dicarboxylate metabolism | 0.012548 | 2.287215 | 6.46E-05 | 0.001393473 | 30 |
| rno00140 Steroid hormone biosynthesis | 0.010314 | 2.223382 | 7.77E-05 | 0.001563877 | 63 |
| rno00280 Valine. leucine and isoleucine degradation | 0.012153 | 2.206523 | 8.96E-05 | 0.00169083 | 53 |
| rno00260 Glycine. serine and threonine metabolism | 0.016156 | 2.123065 | 0.000165 | 0.002876338 | 37 |
| rno00640 Propanoate metabolism | 0.016158 | 2.128827 | 0.000171 | 0.002876338 | 33 |
| rno05322 Systemic lupus erythematosus | 0.018077 | 2.072023 | 0.000184 | 0.002926358 | 101 |
| rno00620 Pyruvate metabolism | 0.023662 | 1.983002 | 0.000383 | 0.005777395 | 43 |
| rno00590 Arachidonic acid metabolism | 0.023741 | 1.886482 | 0.000629 | 0.009042865 | 64 |
| **FLU vs. CUS Down** | | | | | |
| rno03010 Ribosome | 6.88E-14 | -7.64599 | 5.67E-37 | 1.71E-34 | 179 |
| rno00190 Oxidative phosphorylation | 2.4E-05 | -3.65238 | 5.14E-10 | 7.76E-08 | 121 |
| rno05012 Parkinson's disease | 0.000174 | -3.14819 | 3.63E-08 | 3.65E-06 | 234 |
| rno05140 Leishmaniasis | 0.002775 | -2.80947 | 8.61E-07 | 6.5E-05 | 67 |
| rno04145 Phagosome | 0.003617 | -2.60376 | 3.79E-06 | 0.000229 | 165 |
| rno05144 Malaria | 0.003044 | -2.55568 | 9.56E-06 | 0.000481 | 48 |
| rno05020 Prion diseases | 0.002185 | -2.40696 | 1.77E-05 | 0.000764 | 255 |
| rno05323 Rheumatoid arthritis | 0.012573 | -2.24684 | 5.87E-05 | 0.002216 | 77 |
| rno04932 Non-alcoholic fatty liver disease (NAFLD) | 0.008409 | -2.17258 | 9.57E-05 | 0.003211 | 149 |
| rno00260 Glycine. serine and threonine metabolism | 0.01259 | -2.19198 | 0.000109 | 0.003291 | 37 |
| rno05145 Toxoplasmosis | 0.018678 | -2.08496 | 0.000168 | 0.004601 | 102 |
| rno05322 Systemic lupus erythematosus | 0.018991 | -2.06535 | 0.000191 | 0.004813 | 101 |
| rno05150 Staphylococcus aureus infection | 0.016696 | -2.05492 | 0.000215 | 0.004986 | 83 |
| rno05310 Asthma | 0.022106 | -2.06501 | 0.000289 | 0.006232 | 20 |
| rno04621 NOD-like receptor signaling pathway | 0.024133 | -1.92615 | 0.000448 | 0.008931 | 157 |
| rno05321 Inflammatory bowel disease (IBD) | 0.027741 | -1.93113 | 0.000473 | 0.008931 | 53 |
| rno00590 Arachidonic acid metabolism | 0.032888 | -1.81974 | 0.000901 | 0.015635 | 64 |
| rno04612 Antigen processing and presentation | 0.023633 | -1.82068 | 0.000932 | 0.015635 | 72 |
| rno05164 Influenza A | 0.033614 | -1.78943 | 0.001012 | 0.01607 | 150 |
| rno05320 Autoimmune thyroid disease | 0.018752 | -1.81676 | 0.001064 | 0.01607 | 54 |
| rno04672 Intestinal immune network for IgA production | 0.034895 | -1.78465 | 0.001186 | 0.016366 | 40 |
| rno01200 Carbon metabolism | 0.014885 | -1.77673 | 0.001192 | 0.016366 | 112 |
| rno05330 Allograft rejection | 0.020283 | -1.77959 | 0.001361 | 0.017871 | 48 |
| **FLU vs. CUS Up** | | | | | |
| rno04070 Phosphatidylinositol signaling system | 0.002884 | 2.788419 | 9.15E-07 | 0.000144 | 92 |
| rno04360 Axon guidance | 0.002213 | 2.77186 | 9.52E-07 | 0.000144 | 178 |
| **LPS vs. CUS Down** | | | | | |
| rno05145 Toxoplasmosis | 0.003641 | -2.52348 | 8.18E-06 | 0.001347 | 102 |
| rno05140 Leishmaniasis | 0.004667 | -2.5223 | 8.92E-06 | 0.001347 | 67 |
| **LPS vs. CUS Up** | | | | | |
| - | - | - | - | - | - |
| **EPA vs. CUS Down** | | | | | |
| rno03010 Ribosome | 6.96E-08 | -4.28398 | 4.98E-13 | 1.5E-10 | 179 |
| rno05323 Rheumatoid arthritis | 0.004165 | -2.66016 | 2.75E-06 | 0.000416 | 77 |
| rno05140 Leishmaniasis | 0.004836 | -2.58308 | 5.34E-06 | 0.000538 | 67 |
| rno00190 Oxidative phosphorylation | 0.001654 | -2.18848 | 0.000114 | 0.008574 | 121 |
| rno04060 Cytokine-cytokine receptor interaction | 0.01262 | -2.00647 | 0.000272 | 0.016433 | 236 |
| **EPA vs. CUS Up** | | | | | |
| - | - | - | - | - | - |
| **FLU+LPS vs. CUS Down** | | | | | |
| rno03010 Ribosome | 2.99E-09 | -5.74512 | 2E-15 | 6.03E-13 | 179 |
| rno05140 Leishmaniasis | 0.002596 | -2.83132 | 4.17E-05 | 0.0063 | 67 |
| **FLU+LPS vs. CUS Up** | | | | | |
| - | - | - | - | - | - |
| **FLU+EPA vs. CUS Down** | | | | | |
| rno04510 Focal adhesion | 5.88E-05 | -3.63509 | 2.6E-10 | 7.86E-08 | 196 |
| rno04512 ECM-receptor interaction | 6.45E-05 | -3.54385 | 1.74E-09 | 2.63E-07 | 83 |
| rno04151 PI3K-Akt signaling pathway | 0.001167 | -2.92016 | 2.41E-07 | 2.43E-05 | 324 |
| rno04933 AGE-RAGE signaling pathway in diabetic complications | 0.004219 | -2.53902 | 7.04E-06 | 0.000532 | 99 |
| rno05145 Toxoplasmosis | 0.007228 | -2.4474 | 1.33E-05 | 0.000805 | 102 |
| rno04621 NOD-like receptor signaling pathway | 0.008092 | -2.37199 | 2.21E-05 | 0.001114 | 157 |
| rno04974 Protein digestion and absorption | 0.001949 | -2.34724 | 3.93E-05 | 0.001696 | 94 |
| rno04611 Platelet activation | 0.006097 | -2.27743 | 4.76E-05 | 0.001798 | 119 |
| rno04010 MAPK signaling pathway | 0.01024 | -2.20882 | 6.88E-05 | 0.001958 | 284 |
| rno02010 ABC transporters | 0.011916 | -2.24002 | 7.1E-05 | 0.001958 | 49 |
| rno04371 Apelin signaling pathway | 0.010845 | -2.21227 | 7.13E-05 | 0.001958 | 133 |
| rno01521 EGFR tyrosine kinase inhibitor resistance | 0.010532 | -2.14939 | 0.000121 | 0.002947 | 78 |
| rno04724 Glutamatergic synapse | 0.00661 | -2.13391 | 0.000133 | 0.002947 | 111 |
| rno04015 Rap1 signaling pathway | 0.008291 | -2.11425 | 0.000137 | 0.002947 | 207 |
| rno04360 Axon guidance | 0.003032 | -2.08704 | 0.000177 | 0.003567 | 178 |
| rno05222 Small cell lung cancer | 0.017472 | -2.03601 | 0.000239 | 0.004506 | 90 |
| rno04020 Calcium signaling pathway | 0.005828 | -2.00106 | 0.000293 | 0.004795 | 231 |
| rno04540 Gap junction | 0.011051 | -2.01145 | 0.0003 | 0.004795 | 84 |
| rno05146 Amoebiasis | 0.016846 | -2.00338 | 0.000302 | 0.004795 | 87 |
| rno05161 Hepatitis B | 0.021089 | -1.94636 | 0.000399 | 0.006029 | 147 |
| rno01522 Endocrine resistance | 0.013454 | -1.91814 | 0.00052 | 0.007161 | 93 |
| rno04620 Toll-like receptor signaling pathway | 0.025713 | -1.90465 | 0.000536 | 0.007161 | 83 |
| rno04725 Cholinergic synapse | 0.00864 | -1.91193 | 0.000545 | 0.007161 | 111 |
| rno04072 Phospholipase D signaling pathway | 0.017941 | -1.88564 | 0.00059 | 0.007424 | 147 |
| rno04914 Progesterone-mediated oocyte maturation | 0.02414 | -1.85995 | 0.000707 | 0.008538 | 89 |
| rno05032 Morphine addiction | 0.012548 | -1.85894 | 0.000757 | 0.008565 | 91 |
| rno05142 Chagas disease (American trypanosomiasis) | 0.029892 | -1.84167 | 0.000766 | 0.008565 | 100 |
| rno04730 Long-term depression | 0.029119 | -1.83222 | 0.000864 | 0.009268 | 58 |
| rno04062 Chemokine signaling pathway | 0.023332 | -1.80815 | 0.000917 | 0.009268 | 174 |
| rno04668 TNF signaling pathway | 0.032469 | -1.80436 | 0.000946 | 0.009268 | 106 |
| rno05205 Proteoglycans in cancer | 0.016155 | -1.80368 | 0.000951 | 0.009268 | 197 |
| rno04931 Insulin resistance | 0.026694 | -1.78151 | 0.001102 | 0.010205 | 104 |
| rno04064 NF-kappa B signaling pathway | 0.029837 | -1.77929 | 0.001115 | 0.010205 | 93 |
| rno04014 Ras signaling pathway | 0.028782 | -1.73637 | 0.001365 | 0.012129 | 223 |
| rno04390 Hippo signaling pathway | 0.026636 | -1.70776 | 0.001646 | 0.013874 | 153 |
| rno04610 Complement and coagulation cascades | 0.02232 | -1.72178 | 0.001654 | 0.013874 | 78 |
| rno04658 Th1 and Th2 cell differentiation | 0.041134 | -1.69311 | 0.001802 | 0.014432 | 84 |
| rno04024 cAMP signaling pathway | 0.022108 | -1.68776 | 0.001816 | 0.014432 | 208 |
| rno04713 Circadian entrainment | 0.027828 | -1.6663 | 0.002143 | 0.016381 | 94 |
| rno00534 Glycosaminoglycan biosynthesis - heparan sulfate / heparin | 0.041827 | -1.69707 | 0.00217 | 0.016381 | 24 |
| rno04380 Osteoclast differentiation | 0.045012 | -1.62152 | 0.002599 | 0.018698 | 119 |
| rno04068 FoxO signaling pathway | 0.044186 | -1.62141 | 0.0026 | 0.018698 | 122 |
| rno04514 Cell adhesion molecules (CAMs) | 0.023214 | -1.62005 | 0.002699 | 0.018957 | 154 |
| rno04970 Salivary secretion | 0.039651 | -1.61576 | 0.002803 | 0.019242 | 75 |
| **FLU+EPA vs. CUS Up** | | | | | |
| rno03010 Ribosome | 7.06E-14 | 6.712771 | 1.08E-27 | 3.28E-25 | 179 |
| rno00190 Oxidative phosphorylation | 5.91E-07 | 4.880524 | 1.5E-16 | 2.27E-14 | 121 |
| rno05014 Amyotrophic lateral sclerosis (ALS) | 1.77E-05 | 3.902496 | 1.03E-11 | 1.01E-09 | 342 |
| rno05016 Huntington's disease | 2.75E-05 | 3.884531 | 1.33E-11 | 1.01E-09 | 279 |
| rno05012 Parkinson's disease | 6.52E-05 | 3.803676 | 3.32E-11 | 2E-09 | 234 |
| rno05020 Prion diseases | 0.000525 | 3.176303 | 2.32E-08 | 1.17E-06 | 255 |
| rno03040 Spliceosome | 4.8E-05 | 3.235012 | 3.16E-08 | 1.36E-06 | 130 |
| rno03050 Proteasome | 0.002326 | 2.781989 | 1.89E-06 | 7.15E-05 | 46 |
| rno05010 Alzheimer's disease | 0.004894 | 2.337587 | 2.76E-05 | 0.000925 | 351 |
| rno03013 RNA transport | 0.002418 | 2.331785 | 3.43E-05 | 0.001036 | 161 |
| rno04932 Non-alcoholic fatty liver disease (NAFLD) | 0.010099 | 2.289421 | 4.04E-05 | 0.00111 | 149 |
| rno00980 Metabolism of xenobiotics by cytochrome P450 | 0.007811 | 2.280644 | 5.13E-05 | 0.001292 | 70 |
| rno05204 Chemical carcinogenesis | 0.01686 | 1.912128 | 0.00054 | 0.012539 | 80 |
| rno03008 Ribosome biogenesis in eukaryotes | 0.019163 | 1.863833 | 0.000744 | 0.016053 | 79 |
